# Additive and partially dominant effects from genomic variation contribute to rice heterosis

**DOI:** 10.1101/2024.07.16.603817

**Authors:** Zhiwu Dan, Yunping Chen, Wei Zhou, Yanghong Xu, Junran Huang, Yi Chen, Jizhou Meng, Guoxin Yao, Wenchao Huang

**Author notes:** These authors contributed equally to this article (Zhiwu Dan, Yunping Chen).

## Abstract

Heterosis, or hybrid vigor, describes the superior performance of F_1_ hybrids relative to their parents. Despite its significant importance in crop breeding, the molecular mechanisms underlying heterosis remain debated, mainly attributable to discrepancies across genotypes, traits, tissues, populations, developmental stages, growth environments, and species. In this study, we systematically identified heterosis-associated genes and metabolites from parental molecular differences in rice and functionally validated three genes for heterosis of seedling length. We incorporated these heterosis-associated molecules into network modules and explained the variance of heterosis. The predominant inheritance patterns of these molecules were additive and partially dominant effects, namely at mid-parent levels or values between mid-parent and parental levels, respectively. These two genetic effects contributed to heterosis of 17 agronomic traits in rice, including grain yield and plant height across developmental stages. They also explained yield heterosis in diverse hybrid populations and distinct growth environments in both rice and maize, as well as biomass heterosis in *Arabidopsis*. Notably, additive and partially dominant effects were associated with parental genomic variants in rice, and the number of these variants correlated significantly with heterosis of agronomic traits. Furthermore, we demonstrated the significant impact of parental genomic, transcriptomic, and metabolomic variation in phenylpropanoid biosynthesis on heterosis for seedling length/plant height. Unlike classical heterosis models primarily focused on genomic sequence variation, our findings provide quantitative insights from genomic downstream information into the molecular mechanisms of plant heterosis, highlighting their potential for improving breeding efficiency of hybrid crops.

## INTRODUCTION

Heterosis describes the phenomenon where heterozygous F_1_ hybrids outperform their parents (Birchler et al., 2006; Birchler et al., 2010; Shull, 1948). The exploitation of heterosis to enhance crop yield represents a landmark innovation in modern agriculture for mitigating food shortages (Hochholdinger and Baldauf, 2018). Several well-known models have been proposed to explain heterosis, including dominance (pseudo-overdominance) (Bruce, 1910; Davenport, 1908; Jones, 1917; Li et al., 2015; Xiao et al., 2021), overdominance (Crow, 1948; East, 1936), and epistasis (Li et al., 1997; Powers, 1944; Yu et al., 1997), emphasizing complementation, heterozygosity, and interactions of loci at the genomic level, respectively. However, the molecular mechanisms underpinning crop heterosis remain controversial, largely due to diversity in factors such as genotypes, phenotypes, tissues, hybrid populations, developmental stages, growth environments, and species.

To quantitatively describe the inheritance patterns of hybrid phenotypes and molecular levels of F_1_ hybrids, geneticists used four key terms: additive effect (mid-parent value, MPV), partially dominant effect (between MPV and parental values), dominant effect (parental values), and overdominant effect (beyond parental values) (Lisec et al., 2011; Schnable and Springer, 2013). Accumulating evidence suggests a connection between additive effect and heterosis. For example, the near-additive expression of *SINGLE FLOWER TRUSS* drives strong yield heterosis in tomato (Krieger et al., 2010). Moreover, additive effect of transcripts, proteins, metabolites, and component traits has been implicated as the genetic basis of heterosis in maize, rice, and tomato (Dan et al., 2021; Dan et al., 2019; Dan et al., 2020; Dan et al., 2015; Dan et al., 2016; Wang et al., 2014; Williams, 1959; Zhou et al., 2019b). Partially dominant effect has also been suggested to play a role in heterosis (Jones, 1917; Powers, 1944; Sun et al., 2023; Williams, 1959).

In this study, we systematically identified heterosis-associated genes and metabolites for seedling length in rice and analyzed the proportions and contributions of the four inheritance patterns. Our results demonstrate that additive and partially dominant effects predominantly account for plant heterosis, a finding consistent across diverse traits, hybrid populations, developmental stages, growth environments, omics data types, and even species.

## RESULTS

### Heterosis-associated genes identified from parental transcriptomic differences

To identify heterosis-associated genes (HAGs) in rice, we cultivated cultivars Yuetai B (YB, *indica*), Balilla (*japonica*), and their reciprocal F_1_ hybrids under sterile conditions. Seedling length and weight were measured at 5, 10, and 15 days after sowing (DAS) (Figure 1A and Supplemental Table 1). The two traits were positively correlated in parents and F_1_ hybrids and seedling length was selected for subsequent analysis (Supplemental Figure 1A). Unexpectedly, one F_1_ hybrid (YB×Balilla) exhibited positive better-parent heterosis (BPH) as early as 5 DAS (Figure 1A). Both F_1_ hybrids exhibited positive BPH at 15 DAS. Moreover, genotype, stage, and their interaction explained nearly all variance in seedling length (adjusted R^2^ = 0.99; Supplemental Figure 1B), confirming minimal environmental influence on phenotypic changes.

**Figure 1.**
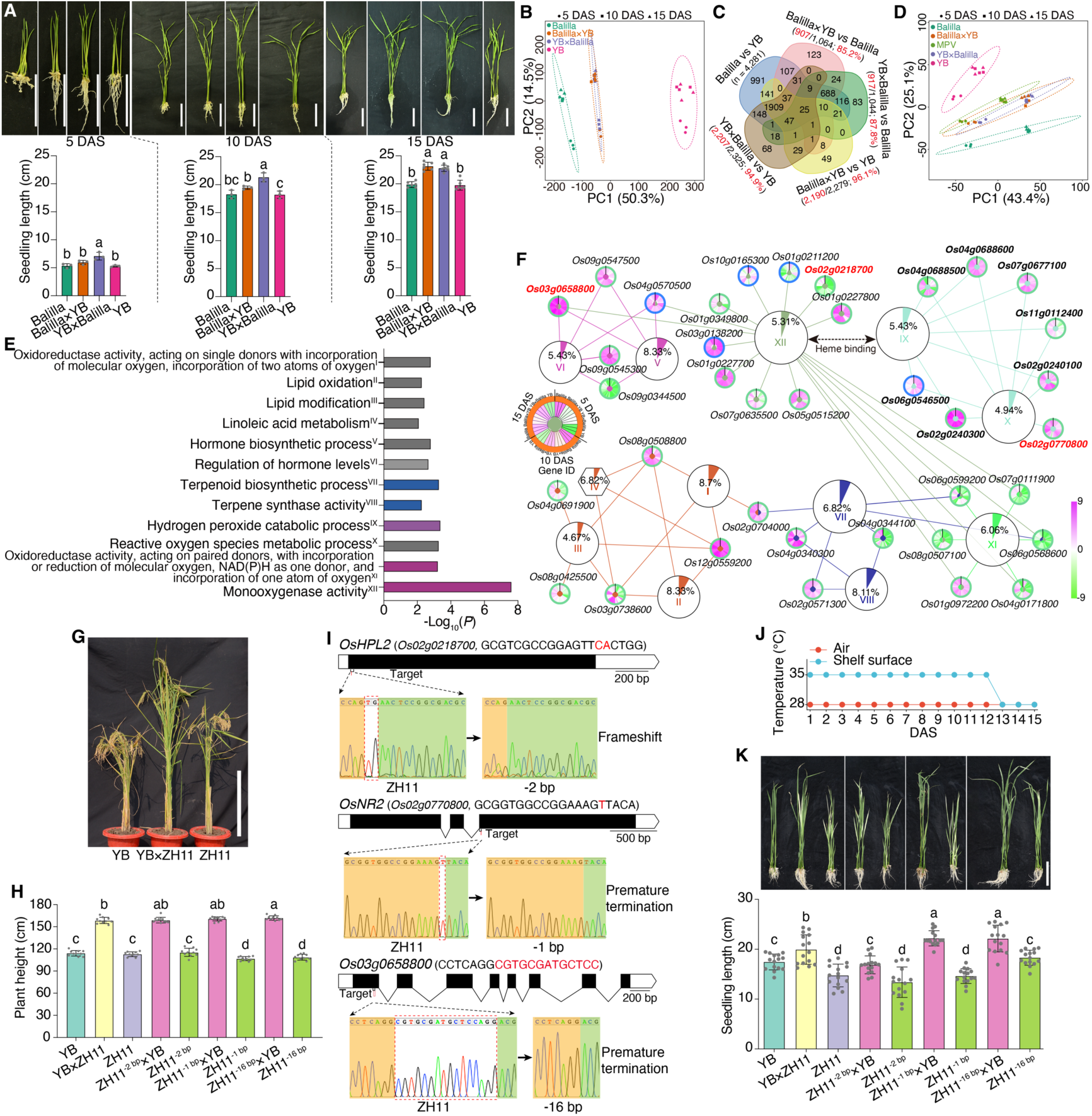
Identification and functional validation of heterosis-associated genes for seedling length in rice. **(A)** Seedlings and seedling length of Balilla, Yuetai B (YB), and their reciprocal F_1_ hybrids at 5, 10, and 15 days after sowing (DAS). Scale bar, 5 cm. **(B)** Principal Component Analysis (PCA) of Balilla, YB, and F_1_ hybrids based on the normalized expression levels of 22,194 genes. **(C)** Venn diagram for differentially expressed genes (DEGs) of parental inbred lines and F_1_ hybrids at 15 DAS. Red texts indicate the number and percentage of DEGs derived from parental differences. **(D)** PCA score plot of Balilla, YB, F_1_ hybrids, and mid-parent values (MPVs) based on 626 heterosis-associated genes. **(E)** Twelve Gene Ontology (GO) terms of 626 heterosis-associated genes. Colored terms denote those overlapping with parental differential terms. **(F)** Network of 37 heterosis-associated genes. Percentages indicate the proportion of genes associated with a GO term. Green/blue outer circles denote up-/downregulation in F_1_ hybrids relative to the parents. **(G)** Plant morphology of YB, Zhonghua11 (ZH11), and their F_1_ hybrid at maturity. Scale bar, 50 cm. **(H)** Plant height of YB, ZH11, ZH11 mutants, and their F_1_ hybrids. n = 10 plants. **(I)** Gene structures and mutation sites of *OsHPL2*, *OsNR2*, and *Os03g0658800* in ZH11 mutant lines. Target sequences and their locations are indicated. **(J)** Daily temperature profile over the 15-day experimental period. **(K)** Seedlings and seedling length of YB, ZH11, ZH11 mutants, and their F_1_ hybrids at 15 DAS. Scare bar, 5 cm. n = 15 seedlings. Analysis of variance with least significant difference used in post hoc test was implemented to compare differences in **A**, **H**, and **K**. The significance level was set at 0.05.

We subsequently performed transcriptomic analysis on these seedling samples and used the normalized expression levels of 22,194 genes for principal component analysis (PCA; Supplemental Table 2 and Supplemental Dataset 1). Though the two parental inbred lines (ILs) had no difference in seedling length at each stage, they located at different regions of the plot, inferring the existence of gene expression heterogeneity among rice ILs. The F_1_ hybrids occupied an intermediate position between the parents, with a closer alignment to Balilla and greater number of differentially expressed genes (DEGs) to YB (Figure 1B and Supplemental Figure 1C). The number of DEGs among the two ILs and F_1_ hybrids increased over time (Supplemental Figure 1D-E), implicating the involvements of more genes during development. Unlike the pooled 16,350 non-DEGs (Supplemental Figure 1F), the 5,844 DEGs explained 66.6% of sample variance in genotypes along principal component 1 (PC1; Supplemental Figure 1G). Furthermore, the F_1_ hybrids deviated slightly from “additive hybrids” (calculated as MPVs), suggesting contributions from additive and non-additive effects to hybrid transcriptomes.

The DEGs were then treated as candidates underlying phenotypic changes between F_1_ hybrids and their parents. The parental ILs harbored the most DEGs at each stage. Crucially, 83.5-97.1% (mean 92.38%) of hybrid-parent DEGs originated from parental expression differences (Figure 1C and Supplemental Figure 2A-B). With Balilla showing higher expression levels than that of YB across stages, the F_1_ hybrids showed more up-regulated than down-regulated DEGs relative to each parent (Supplemental Figure 2C). Furthermore, the significantly enriched Gene Ontology (GO) terms for hybrid-parent comparisons largely overlapped with those for parental ILs (Supplemental Figure 2D-I), indicating that parental expression differences predominantly shape hybrid transcriptomes.

To focus on genes mainly affected by genotype rather than developmental stage, we grouped DEGs using analysis of variance (ANOVA; Supplemental Figure 3A). We identified 1,476 DEGs responsive to genotype, stage, and their interaction. After excluding 275 DEGs specific to ILs or reciprocal F_1_ hybrids (Supplemental Figure 3B), 1,201 DEGs remained. These DEGs retained the sample distribution patterns of the full gene set (Supplemental Figure 3C), with 79.27% up-regulated in F_1_ hybrids (Supplemental Figure 3D). These genes participated in 17 GO terms, ten of which overlapped with parental differential terms (Supplemental Figure 3E).

Correlation analysis between expression levels of the 1,201 DEGs and seedling length of ILs and hybrids identified 626 genes with significant correlations (*P* < 0.05). These genes also contributed substantially to seedling length in partial least squares (PLS) analysis (Supplemental Figure 3F-G). Moreover, correlation coefficients strengthened in F_1_ hybrids versus parents when shifting from the full gene set to these core genes (Supplemental Figure 3H-J), validating them as HAGs.

Unexpectedly, PCA of the 626 HAGs positioned F_1_ hybrids near MPVs (Figure 1D). Among these, 483 HAGs were up-regulated in F_1_ hybrids at one or more stages (Supplemental Figure 3K), inferring phased up-regulation drives heterosis. Most up-regulated HAGs correlated positively with seedling length, and about 50% of down-regulated correlated negatively (Supplemental Figure 3L). GO enrichment analysis of 499 positively and 127 negatively correlated HAGs revealed 37 HAGs in 12 terms (Figure 1E). These formed a tightly connected network bridged by heme binding (Figure 1F), demonstrating functional coherence of genes involved in seedling length heterosis.

### Functional validation of three heterosis-associated genes for seedling length

To functionally validate the identified HAGs, we used CRISPR-Cas9 technology to generate mutants for three genes in Zhonghua 11 (ZH11, *japonica*), whose F_1_ hybrids with YB exhibited strong heterosis for plant height (Figure 1G-H). A 2-bp deletion of nucleotide was generated in exonic region of *OsHPL2* (Figure 1I), which is a gene encoding hydroperoxide lyase (Chehab et al., 2006). A 1-bp deletion of T in exon 3 was detected in gene *OsNR2*, which encodes as a NADH/NADPH-dependent NO_3_^-^ reductase (Gao et al., 2019). The third mutated gene *Os03g0658800* was putatively annotated as a cytochrome P450 and a 16-bp deletion of found in the first exon. The mutation in *OsHPL2* caused frameshift variant and those in other two genes brought premature termination codons.

After selfing to the T_4_ generations, the three mutant lines (designated ZH11^-2^ ^bp^, ZH11^-1^ ^bp^, and ZH11^-16^ ^bp^) were crossed with YB to generate F_1_ hybrids. We first cultivated ZH11, YB, and their F_1_ hybrids (YB×ZH11) under the same conditions used previously for YB, Balilla, and their hybrids: 28 °C with 24-hour light. Under these conditions, ZH11 grew faster than both YB and the F_1_ hybrids, exhibiting significantly longer seedlings at 15 DAS (Supplemental Figure 4A-B). We then tested a 16-hour light/8-hour dark photoperiod at 28 °C, but again found that the seedling length of ZH11 was longer than the F_1_ hybrids as early as 10 DAS (Supplemental Figure 4C-D). This indicated that photoperiod had a minimal effect on seedling length under these conditions.

We next investigated the role of temperature by placing the seedlings on a heated shelf with a surface temperature of approximately 35 °C for 12 days (Figure 1J). Under this higher temperature regime, F_1_ hybrids of YB and ZH11 displayed heterosis for seedling length at 15 DAS (Figure 1K), demonstrating a strong influence of temperature on heterosis. Along with the wild-type F_1_ hybrids, we also cultivated the F_1_ hybrids derived from the ZH11 mutant lines. Compared to the wild-type F_1_ hybrids, the length of YB×ZH11^-2^ ^bp^ hybrids was shorter and displayed a phenotype similar to YB, showing a dominant effect. In contrast, seedling length of YB×ZH11^-1^ ^bp^ and YB×ZH11^-16^ ^bp^ hybrids was both longer than the wild-type hybrids, indicating that mutations in different genes lead to diverse effects on heterosis. Subsequently, we transplanted these seedlings to the field and measured their final plant height. Unlike the observations at the seedling stage, the plant height of YB×ZH11^-2^ ^bp^ and YB×ZH11^-1^ ^bp^ hybrids showed no significant difference from the wild-type hybrids, suggesting roles of developmental stage and environment played in the manifestation of heterosis. However, the YB×ZH11^-16^ ^bp^ hybrids remained significantly taller than the wild-type hybrids, exhibiting an overdominant effect and indicating a robust role of the gene *Os03g0658800* in promoting heterosis.

### Heterosis-associated metabolites from parental metabolomic variation

To identify heterosis-associated metabolites, we performed untargeted metabolite profiling analysis on the above-mentioned seedling samples of YB, Balilla, and their F_1_ hybrids. PCA of metabolite profiles revealed that the F_1_ hybrids clustered intermediately between their parents but positioned closer to YB (Figure 2A, Supplemental Figure 5A, and Supplemental Dataset 2). Differential metabolite (DM) analysis of ILs and F_1_ hybrids showed that, unlike DEGs, the number of DMs decreased during development (Supplemental Figure 5B-C). Genotypic variation was clearly indicated along PC1 based on pooled DMs, consistent with the transcriptomic patterns (Supplemental Figure 5D-E). Meanwhile, the DMs were generally up-regulated in reciprocal F_1_ hybrids, with no significant difference between YB and Balilla (Supplemental Figure 5F). Although the proportions were lower than in transcriptomic data (Figure 2B), most metabolomic differences between F_1_ hybrids and their parents (average 84.0%) originated from parental variation (Supplemental Figure 5G-I). Pathway enrichment analysis of DMs showed that the majority of significantly enriched metabolic pathways distinguishing F_1_ hybrids from parents were also divergent between the two parents (Figure 2C and Supplemental Figure 6), further supporting that those parental molecular differences primarily determined hybrid-parent molecular variation.

**Figure 2.**
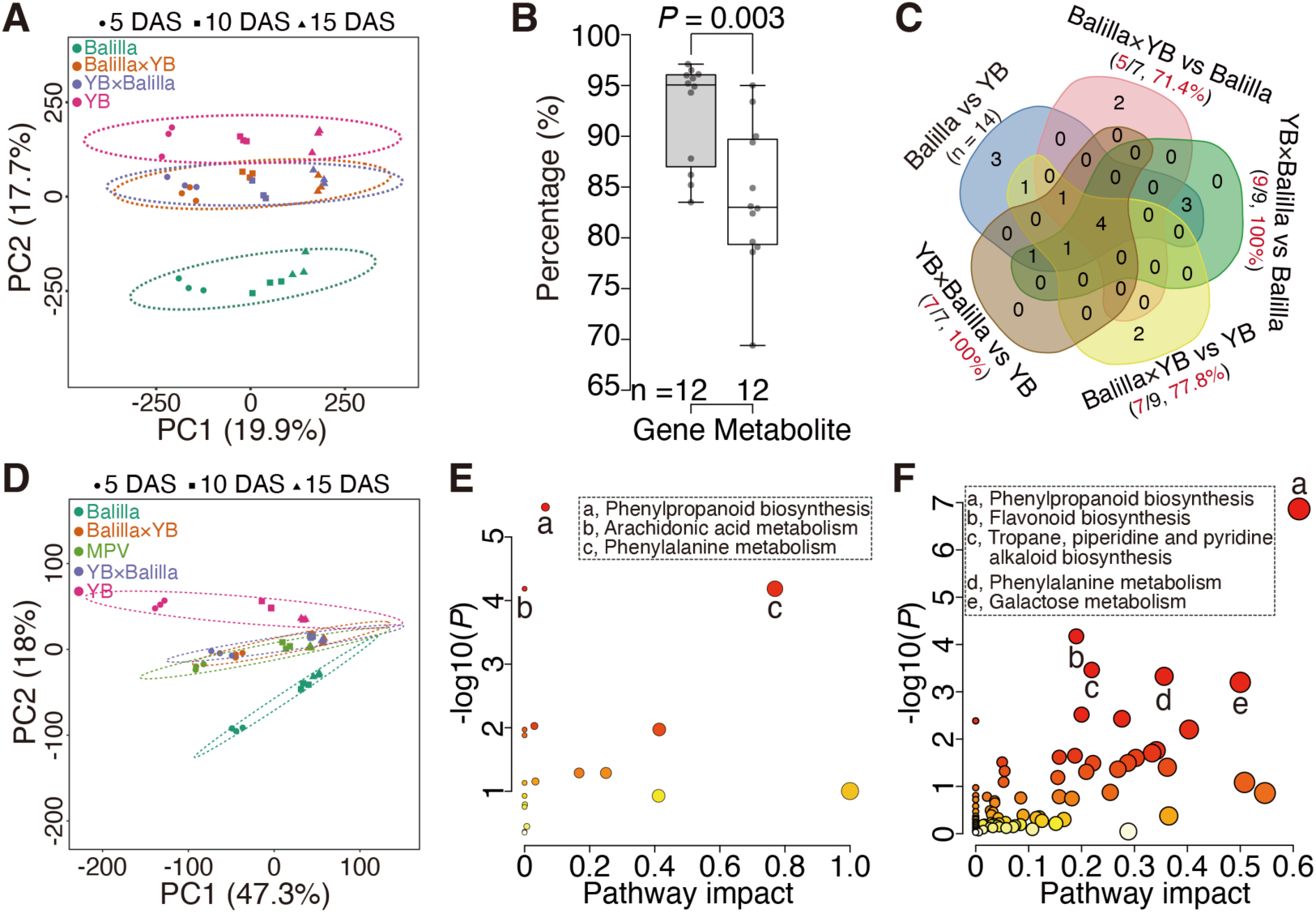
Identification of heterosis-associated metabolites for seedling length in rice. **(A)** PCA score plot of Balilla, YB, and their F_1_ hybrids based on normalized levels of 10,522 metabolites. **(B)** Percentages of differentially expressed genes/metabolites between F_1_ hybrids and parents derived from parental differences. *P* value is for independent samples *t* test. **(C)** Venn diagram for differential metabolic pathways (positive ion mode) of parental inbred lines and F_1_ hybrids at 15 days after sowing (DAS). **(D)** PCA score plot based on 645 heterosis-associated metabolites. **(E)** Enriched pathways of heterosis-associated metabolites (positive ion mode). Three significantly enriched pathways are labeled. **(F)** Integrated pathway enrichment analysis of heterosis-associated genes and metabolites. Five significantly enriched pathways are indicated.

Combining ANOVA, PLS, and correlation analyses identified 645 heterosis-associated metabolites (HAMs) of seedling length (Supplemental Figure 7). With these metabolites, the F_1_ hybrids exhibited metabolite levels closely resembling “additive hybrids” (Figure 2D). Based on annotation information with *mummichog* algorithm (Li et al., 2013), we found eight HAMs were significantly enriched in three pathways: phenylpropanoid biosynthesis, arachidonic acid metabolism, and phenylalanine metabolism (Figure 2E). After chemically annotated metabolites from these pathways were added for correlation analysis, we found that they were positively or negatively correlated with seedling length (Supplemental Figure 8), indicating distinct contributions from metabolites within the same pathway. Notably, three chemically annotated metabolites from phenylpropanoid biosynthesis (4-hydroxycinnamic acid, L-phenylalanine, and trans-ferulic acid) all showed significant correlations with seedling length in F_1_ hybrids, but not in their parents (Supplemental Figure 9).

Joint pathway analysis integrating the 626 HAGs and 645 HAMs identified five significantly enriched pathways (Figure 2F). Phenylpropanoid biosynthesis was the most significant pathway, containing eight metabolites and ten genes. Among these genes, nine encoded peroxidases, seven of which were included in the 37 previously identified HAGs (Supplemental Figure 10). This finding underscored the combined contributions of transcriptomic and metabolomic changes in phenylpropanoid biosynthesis to seedling length heterosis.

### Integrating parental differential molecules into network modules explains the variance of heterosis

To validate contributions of the pooled HAGs and HAMs to heterosis, we analyzed phenotypic and omics data from nine additional reciprocal F_1_ hybrid pairs, each exhibiting varying level of BPH for seedling length (Figure 3A-B). Neither PCA nor partial least squares-discriminant analysis (PLS-DA) successfully separated hybrids into high-, low-, and no-BPH groups using the 626 HAGs (Supplemental Figure 11A-D). The 645 HAMs clearly distinguished the high-BPH group from others without overfitting but failed to classify low- and no-BPH groups (Supplemental Figure 11E-H). This indicated that HAGs or HAMs specific to a single hybrid pair had limited predictive power for heterosis in F_1_ hybrids with divergent genetic background.

**Figure 3.**
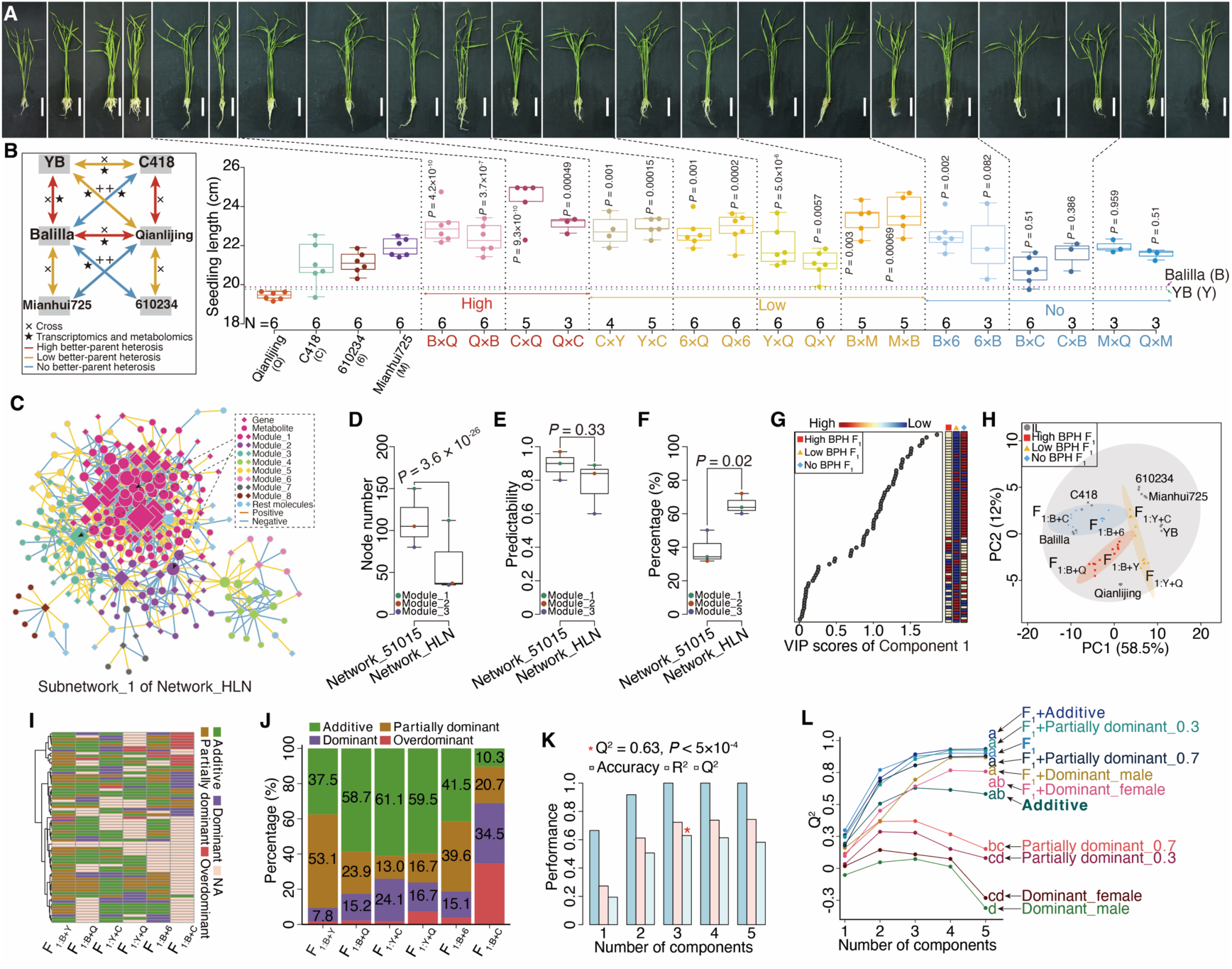
Inheritance patterns and predictabilities of heterosis-associated molecules for seedling length. **(A)** Seedlings and seedling length of four inbred lines and nine pairs of reciprocal F_1_ hybrids. Seedling length was compared between F_1_ hybrid and its better-parent. *P* values indicate least significant difference in post hoc test of analysis of variance. **(B)** Cross scheme of six inbred lines. **(C)** Subnetwork_1 of Network_HLN integrating heterosis-associated genes and metabolites. The subnetwork contains eight modules and 868 edges. Node size reflects node degree. **(D-F)** Comparison of Subnetwork_1 between Network_51015 and Network_HLN. Node number (**D**; weighted), predictability (**E**), and percentage of Component 1 (**F**) of top three modules are compared. *P* values are for independent samples *t* test. **(G)** Scores of variable importance in projections (VIP) of Component 1 for 68 heterosis-associated molecules across F_1_ hybrids with high-, low-, or no better-parent heterosis (BPH). **(H)** PCA score plot of six inbred lines and their reciprocal F_1_ hybrids based on 68 heterosis-associated molecules. **(I)** Inheritance patterns of 68 molecules in six pairs of reciprocal F_1_ hybrids. NA = unclassifiable inheritance patterns. **(J)** Percentages of four inheritance patterns. **(K)** Performance of additive effect. **(L)** Predictabilities (Q^2^) of hybrid metabolite profiles (F_1_), individual inheritance patterns, and their combinations for seedling length heterosis. Analysis of variance was performed with the least significant difference used in post hoc test.

We next integrated HAGs and HAMs into Network_51015, whose largest subnetwork (subnetwork_1) comprised seven modules and 2,138 edges (Supplemental Figure 12A). Unlike the decreasing trend in parents for the top three modules, coefficients of module levels (average node levels) and seedling length maintained greater than 0.97 in F_1_ hybrids (Supplemental Figure 12B-D). Notably, PLS-DA separated F_1_ hybrids into distinct groups, with Module_2 showing high predictability (Q^2^ = 0.97; Supplemental Figure 12E-J). This result indicated that integrating HAGs and HAMs into network modules enhanced the predictability of seedling length heterosis.

We identified 967 DEGs and 990 DMs across six ILs of the ten reciprocal F_1_ hybrid pairs. The DEGs displayed better separation in PLS-DA than PCA, with predictability comparable to DMs (Supplemental Figure 13A-H). Combining gene expression and metabolite levels separated the three groups along PC1 and PC4 but caused overfitted in PLS-DA (Supplemental Figure 13I-L). We then constructed Network_HLN through integrating these DEGs and DMs, with the largest subnetwork (subnetwork_1) contained eight modules and 868 edges (Figure 3C). Though the top three modules in subnetwork_1 from Network_HLN had fewer nodes than those in Network_51015 (Figure 3D), their predictabilities were comparable (Figure 3E and Supplemental Figure 14A-F), and Component 1 explained greater variance in Network_HLN (Figure 3F). Combining Module_1 and Module_2 (149 nodes) increased predictability to 0.94, but adding Module_3 provided no further improvement (Supplemental Figure 14G-J). Thus, integrating parental transcriptomic and metabolomic differences into network modules effectively explained the variance in seedling length heterosis.

### Heterosis-associated molecules primarily exhibit additive and partially dominant effects

We further identified heterosis-associated molecules specific to seedling length by applying PLS analysis to Module_1 and Module_2 of Network_HLN. This yielded 68 high-contribution molecules (29 metabolites and 39 genes) that effectively separated the three groups with high predictability (Q^2^ = 0.90; Supplemental Figure 14K-L). However, these molecules did not exhibit continuous expression or abundance across the high-, low-, and no-BPH groups along either Component 1 or Component 2, which together explained 75.1% of group variance (Figure 3G and Supplemental Figure 15). Additionally, F_1_ hybrids within the same heterotic group clustered distinctly and exhibited divergent molecular levels (Supplemental Figure 16), highlighting molecular heterogeneity among F_1_ hybrids with similar degrees of heterosis. Heatmap and PCA analyses further revealed that hybrid molecular profiles were typically intermediate between the parent levels or aligned with one parent (Figure 3H), demonstrating quantitative inheritance of heterosis-associated molecules from parents to hybrids.

We next characterized the inheritance patterns of these molecules, observing all four genetic types: additive, partially dominant, dominant, and overdominant effects (Figure 3I). Additive and partially dominant effects collectively accounted for over 70% of molecular-level variation in most F_1_ hybrids (Figure 3J and Supplemental Figure 17A). This predominance persisted when genes and metabolite were analyzed separately and when restricted to the top 50 contributing molecules (Supplemental Figure 17B-D). The additive effect showed outstanding predictability for seedling length heterosis (Q^2^ = 0.63, *P* < 5×10^-4^; Figure 3K). Partially dominant (PD) effects were subdivided into PD_0.3 and PD_0.7 subtypes (Supplemental Figure 18), with respective predictability of 0.37 (*P* < 5×10^-4^) and 0.29 (*P* < 5×10^-4^; Figure 3L). In contrast, predictabilities of dominant effects were non-significant for the female parent (Q^2^ = 0.14, *P* = 0.40) and marginal for the male parent (Q^2^ = 0.07, *P* = 0.015; Figure 3L). Importantly, the predictability of additive effect was statistically indistinguishable from that of hybrid profiles, and the additions of additive, partially dominant, or dominant effects to hybrid profiles did not improve predictability (Figure 3L). Furthermore, the combined proportion of additive and partially dominant effects positively correlated with seedling length heterosis (Supplemental Figure 19). Together, these results underscored the predominant roles of additive and partially dominant effects in heterosis at both transcriptomic and metabolomic levels.

### Additive and partially dominant effects contribute to heterosis in rice, maize, and *Arabidopsis*

Since seedling length and plant height represent the same trait measured at different developmental stages in rice, we quantified plant height for above wild type F_1_ hybrids across four stages (28 days post-transplanting to maturation). Significant positive correlations existed between heterosis for seedling length and heterosis for plant height (Supplemental Figure 20A-B), inferring shared genetic mechanisms underlying heterosis along development. Additionally, correlation coefficients declined from early to mature stages, suggesting increasing environmental influences on heterosis manifestation.

To evaluate the contribution of different inheritance patterns to heterosis for plant height and other agronomic traits, we then constructed a hybrid population using a complete diallel cross design. The 17 parental ILs represented genetic diversity within a panel of 223 accessions encompassing *indica*, intermediate types, and *japonica* (Figure 4A-B, Supplemental Figure 20C-D, and Supplemental Dataset 3). We measured 17 agronomic traits (yield, yield components, and yield-related traits) for the parental ILs and their 245 F_1_ hybrids. Fourteen traits, including yield per plant, tiller number, and plant height, displayed higher values in F_1_ hybrids compared to parents (Supplemental Figure 20E-F). We calculated BPH and revealed negative average BPH for only four traits (Figure 4C), confirming superior fitness in F_1_ hybrids. Trait correlation patterns differed between parents and hybrids: yield per plant tightly clustered with seed setting rate, grain weight, and heading date in parental ILs, but with plant height in F_1_ hybrids (Supplemental Figure 20G-H). Notably, heterosis for the 17 traits displayed correlation patterns similar to those of the parental ILs (Figure 4D). These results indicated that crossing genetically divergent ILs achieved positive heterosis for most agronomic traits, with heterosis correlations mirroring parental phenotypic correlations.

**Figure 4.**
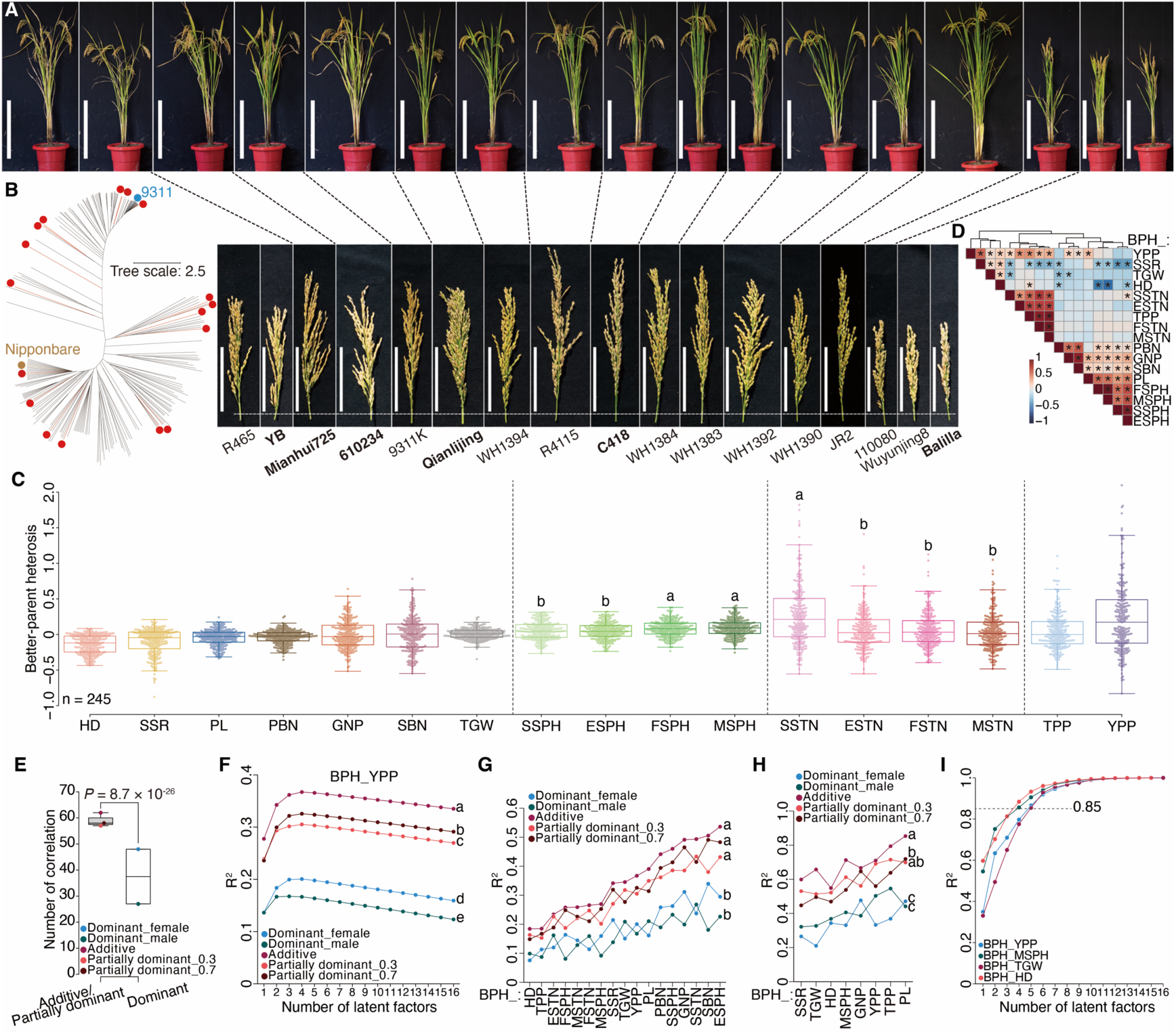
Additive and partially dominant effects contribute to heterosis of 17 agronomic traits in rice. **(A)** Plants and panicles of 17 inbred lines (ILs) used for complete diallel-crosses. Scale bars represent 50 cm and 10 cm for plants and panicles, respectively. The six ILs used in Figure 3B are in bold. **(B)** Neighbour-joining tree of 223 ILs based on InDels. The 17 selected ILs are with red dots. **(C)** Better-parent heterosis (BPH) of 17 traits in 245 F_1_ hybrids. **(D)** Correlation heatmap for BPH of 17 traits. Asterisks indicate significant Pearson correlations (*P* < 0.05). **(E)** Number of significant correlations between different inheritance patterns of five chemically annotated metabolites and heterosis of 17 traits. *P* value indicates independent samples *t* tests (weighted). **(F)** R^2^ values (adjusted) with varying number of latent factors in partial least squares analysis on better-parent heterosis for grain yield per plant (BPH_YPP). The 1,306 differential metabolites analyzed from 17 ILs were used as predictor with different inheritance patterns. **(G)** Top R^2^ values of models for predicting heterosis of 17 traits. **(H)** Top R^2^ values for predicting heterosis of eight traits across environments. **(I)** R^2^ values for predicting heterosis of four traits in test-cross F_1_ hybrids. Analysis of variance was performed with the least significant difference used in post hoc test in **C**, **F**, **G**, and **H**. Abbreviations: secondary branch number = SBN, grain number per panicle = GNP, tiller number per plant = TPP, seed setting rate = SSR, heading date = HD, thousand grain weight = TGW, seedling stage tiller number = SSTN, elongation stage tiller number = ESTN, flowering stage tiller number = FSTN, maturation stage tiller number = MSTN, seedling stage plant height = SSPH, elongation stage plant height = ESPH, flowering stage plant height = FSPH, maturation stage plant height = MSPH, panicle length = PL, and primary branch number = PBN.

Using metabolite profiles from the 17 ILs, we selected five chemically annotated metabolites from phenylpropanoid biosynthesis to investigate associations between their inheritance patterns and heterosis for plant height (Supplemental Figure 21A-F and Supplemental Dataset 4). Significant correlations emerged between plant height heterosis and four inheritance patterns of these metabolites, except for the male dominant effect of L-phenylalanine (negative ion mode) (Supplemental Figure 21G). Among the five metabolites, 4-hydroxycinnamic acid, which has been linked to seedling length heterosis in our above result and plant height heterosis in previously reported findings (Dan et al., 2021), correlated significantly with plant height heterosis across stages with all inheritance patterns. Meanwhile, correlation coefficients of additive levels of 4-hydroxycinnamic acid and heterosis for plant height decreased during later developmental stages, coinciding with significant increases in heterosis at these stages (Supplemental Figure 21H). Importantly, additive and partially dominant effects of these metabolites correlated more strongly with plant height heterosis than dominant effects (Supplemental Figure 21I), a pattern consistent across an additional 13 traits (Figure 4E). These results showed that phenylpropanoid biosynthesis was associated with heterosis for plant height at the population level and additive and partially dominant effects might be the predominant contributors for heterosis of multiple agronomic traits.

We next analyzed DMs among the 17 ILs. Using 1,306 DMs with different inheritance patterns, we predicted heterosis for each trait via PLS. Additive effect yielded higher adjusted R^2^ values in predictive models compared to dominant and partially dominant effects (Figure 4F and Supplemental Figure 22). The maximum R^2^ values for predicting the 17 traits were significantly greater for additive effect than dominant effects and comparable to partially dominant effects (Figure 4G). When re-measuring eight representative traits for the 17 ILs and 36 F_1_ hybrids from the diallel crosses under different growth conditions, we observed significant differences in almost all traits influenced by environmental effects (Supplemental Figure 23A). However, heterosis remained stable for six of these traits (Supplemental Figure 23B). Consistent with the above result, the highest R² values for predicting heterosis of these traits were greater for additive effect than dominant effects and similar to partially dominant effects (Figure 4H and Supplemental Figure 24). These findings confirmed additive and partially dominant effects as the main inheritance patterns underlying heterosis of the 17 agronomic traits, independent of environmental variation.

To assess the application potential of these findings in hybrid breeding programs, we created a test-cross population, with a Honglian-type cytoplasmic male sterile line (Yuetai A) as female and 211 recombinant inbred lines (RILs, derived from hybrids of the above-mentioned 223 ILs) as male parents (Supplemental Figure 25A-B). Trait values (yield per plant, maturation stage plant height, and heading date) differed significantly between test crosses and parents (Supplemental Figure 25C), and trait correlation patterns also differed (Supplemental Figure 25D-F). For heterosis of these four traits, BPH was negative only for heading date (Supplemental Figure 25G). Metabolite profiles from 107 RILs (selected for varied yield heterosis) and Yuetai A, which were profiled alongside the 17 ILs, were used for PLS modeling (Supplemental Figure 25H-I). Despite significant differences in heterosis for yield per plant and heading date between the diallel and test-cross populations (Supplemental Figure 25G), the values of predictive models using additive DMs achieved R^2^ > 0.85 with five latent factors (Figure 4I). Thus, the DMs from representative ILs served as robust biomarkers for predicting heterosis in test-cross hybrids.

To further explore the contributions of additive and partially dominant effects to maize heterosis, we analyzed field fresh weight data from two hybrid populations exhibiting significant different heterosis (Supplemental Figure 26A-B) (de Abreu et al., 2017). PLS analyses utilized root and leave metabolite profiles from 24 Dent lines, 25 Flint lines, and 326 Dent × Flint F_1_ hybrids (Supplemental Figure 26C-J). Unexpectedly, additive and partially dominant effects yielded R^2^ values (maximum) comparable to models using hybrid metabolite profiles directly and significantly higher than dominant effects (Supplemental Figure 26K). Similarly, for grain dry matter yield and grain dry matter content in 291 maize F_1_ hybrids (Schrag et al., 2018), additive and partially dominant effects showed R^2^ values comparable to hybrid profiles and higher than dominant effects (Supplemental Figure 27). Additionally, we analyzed maize grain yield heterosis for a third hybrid population (Supplemental Figure 28A-B), with the 66 F_1_ hybrids showing strong heterosis (Wang et al., 2023). We used expression data of 84 DEGs, which were primarily core genes in pan-genome analysis (Supplemental Figure 28C), from 12 representative maize ILs. Consistent with metabolite results, additive and partially dominant effects produced greater R^2^ values than dominant effects for predicting yield heterosis (Supplemental Figure 28D-G). These findings further supported the predominant roles of additive and partially dominant effects in maize heterosis.

Similarly, in *Arabidopsis*, phenotypic data from Col-0, Per-1, and their F_1_ hybrids across developmental stages revealed robust heterosis for cotyledon biomass and first true leaf size (Liu et al., 2021b) (Supplemental Figure 29A and Supplemental Figure 29K). Using our HAG screening strategy in rice, we identified HAGs for each phenotype (Supplemental Figure 29B-I and Supplemental Figure 29L-S). Unexpectedly, the hybrid profiles strongly associated with additive and partially dominant effects throughout development (Supplemental Figure 29J and Supplemental Figure 29T).

### Associations of parental genomic variants with additive/partially dominant effects and heterosis

Because both transcriptomic and metabolomic differences in parental ILs were connected to heterosis, we next performed genomic sequencing of the 17 rice ILs to investigate associations between parental genomic variants and heterosis. Genetic diversity analysis revealed two distinct groups of these ILs based on variants including single-nucleotide polymorphisms (SNPs), insertions and deletions (InDels), copy number variants (CNVs), and structural variants (SVs) (Figure 5A-F, Supplemental Figure 30-32, and Supplemental Table 3). Using the Nipponbare genome as a reference, genomic variants in each pair of ILs were categorized as common or unique, whose number correlated significantly with trait heterosis (Figure 5G). The unique variants showed stronger correlations with heterosis than common variants (Figure 5H and Supplemental Figure 33A-B). Specifically, the unique variants significantly correlated with heterosis for seed setting rate, tiller number (elongation stage), plant height (across four stages), and panicle length. Meanwhile, unique SNPs and InDels exhibited stronger heterosis correlation than CNVs and SVs (Figure 5I), likely reflecting limitations of short-read sequencing in resolving large structural variants. Notably, the unique SNPs and InDels correlated significantly with yield heterosis, while CNVs and SVs did not. For heterosis, increased number of unique SNPs or InDels correlated with higher heterosis for grain number but lower heterosis for seed setting rate (Figure 5J and Supplemental Figure 33C-E), revealing barriers of reproductive isolation in the exploitation of yield heterosis between *indica* and *japonica* subspecies (Dan et al., 2014). Consequently, the inverse relationship between grain number and seed setting rate heterosis produced a parabolic pattern in yield heterosis (Figure 5K and Supplemental Figure 33F-H). These results showed that parental genomic variants were tightly linked to heterosis and highlighted the genomic complexity underlying heterosis of polygenic traits like yield.

**Figure 5.**
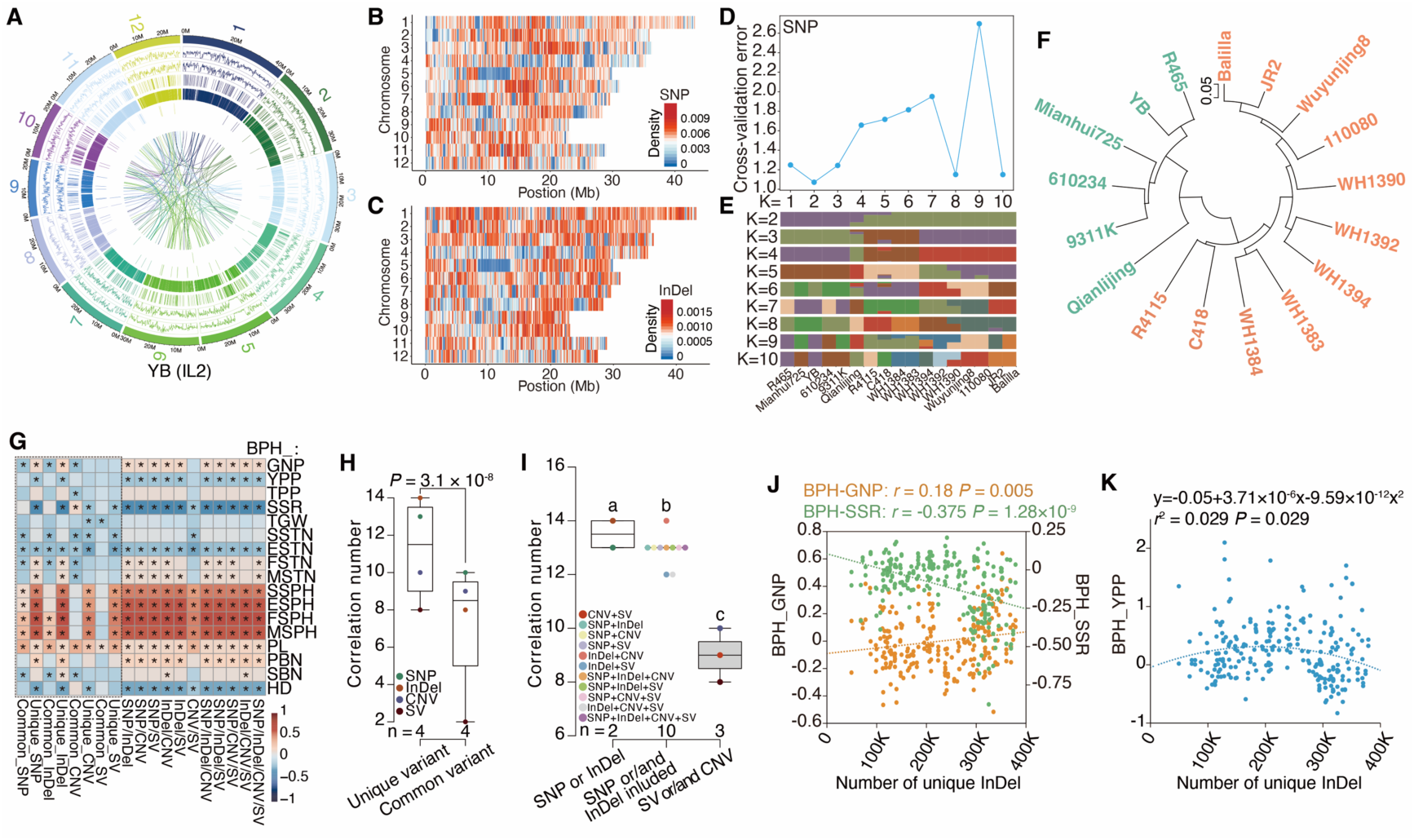
Parental genomic variants correlate with heterosis of agronomic traits in rice. **(A)** Circos visualization of genomic variants in Yuetai B (YB). Chromosome position, SNP density, InDel density, CNV duplication position, CNV deletion position, SV insertion position, SV deletion position, and SV inversion position are listed from outer to inner circles. **(B-C)** Heatmaps for SNP (**B**) and InDel (**C**) density of Yuetai B. **(D)** Cross-validation errors with K values ranging from 1 to 10 using SNPs from 17 inbred lines (ILs). **(E)** Population structure of 17 ILs based on SNPs. **(F)** Neighbour-joining tree of 17 ILs based on SNPs. **(G)** Correlation heatmap between parental genomic variants (common/unique) and heterosis of 17 traits. Asterisks indicate significant Spearman correlations (*P* < 0.05). **(H)** Number of significant correlations between parental genomic variants and trait heterosis. *P* value indicates independent samples *t* test (weighted). **(I)** Number of significant correlations between parental unique variants and trait heterosis. Analysis of variance was performed with the least significant difference used in post hoc test (weighted). **(J)** Correlations between parental unique InDels and heterosis for grain number per plant/seed setting rate. *P* value indicates Spearman rank correlation. **(K)** Quadratic regression between parental unique InDels and heterosis for grain yield per plant. Abbreviations: better-parent heterosis = BPH, yield per plant = YPP, secondary branch number = SBN, grain number per panicle = GNP, tiller number per plant = TPP, seed setting rate = SSR, heading date = HD, thousand grain weight = TGW, seedling stage tiller number = SSTN, elongation stage tiller number = ESTN, flowering stage tiller number = FSTN, maturation stage tiller number = MSTN, seedling stage plant height = SSPH, elongation stage plant height = ESPH, flowering stage plant height = FSPH, maturation stage plant height = MSPH, panicle length = PL, and primary branch number = PBN.

Compared to heterosis for yield, yield components, and yield-related traits, heterosis for plant height, which was less affected by reproductive isolation of subspecies, showed greater correlation coefficients with parental unique variants (Figure 5G). To study the effect of variants in genes from a specific pathway on trait heterosis, we focused on sequences of six HAGs from phenylpropanoid biosynthesis (previously linked to seedling length) in 12 of the 17 ILs. Sequence variants were detected in all the six genes between YB and Balilla. With YB as the reference, SNPs dominated the variants, with several InDels (Figure 6A and Supplemental Figure 34-38). Unexpectedly, presence and absence variants occurred only in *Os09g0262000*, whose function is putatively annotated as cinnamoyl CoA reductase. Sequence variants of each gene clustered the 12 ILs into two or three groups (Figure 6B and Supplemental Figure 34-38), which were close to *indica* and *japonica* groups, implicating the roles of these genes in rice subspeciation. Notably, sequence variants in *Os09g0262000*, particularly two presence variants in the fifth exon, shifted Balilla into the *indica* group (Figure 6B). Using all six genes, the 12 ILs contained two haplotypes (Figure 6C-D). Due to the variants in *Os09g0262000*, Balilla was grouped with six *indica* ILs in haplotype-1. Unexpectedly, the average plant height heterosis of F_1_ hybrids with heterozygous haplotypes (haplotype-1_haplotype-2) were significantly higher than their homozygous counterparts (haplotype-1 and haplotype-2; Figure 6E). These hybrids also exhibited significantly higher heterosis for grain number and primary/secondary branch number, lower heterosis for heading date, and no yield heterosis (Figure 6F-G and Supplemental Figure 39). In combination of results from transcriptomic and metabolomic data, we concluded that phenylpropanoid biosynthesis contributed to heterosis for seedling length/plant height in rice.

**Figure 6.**
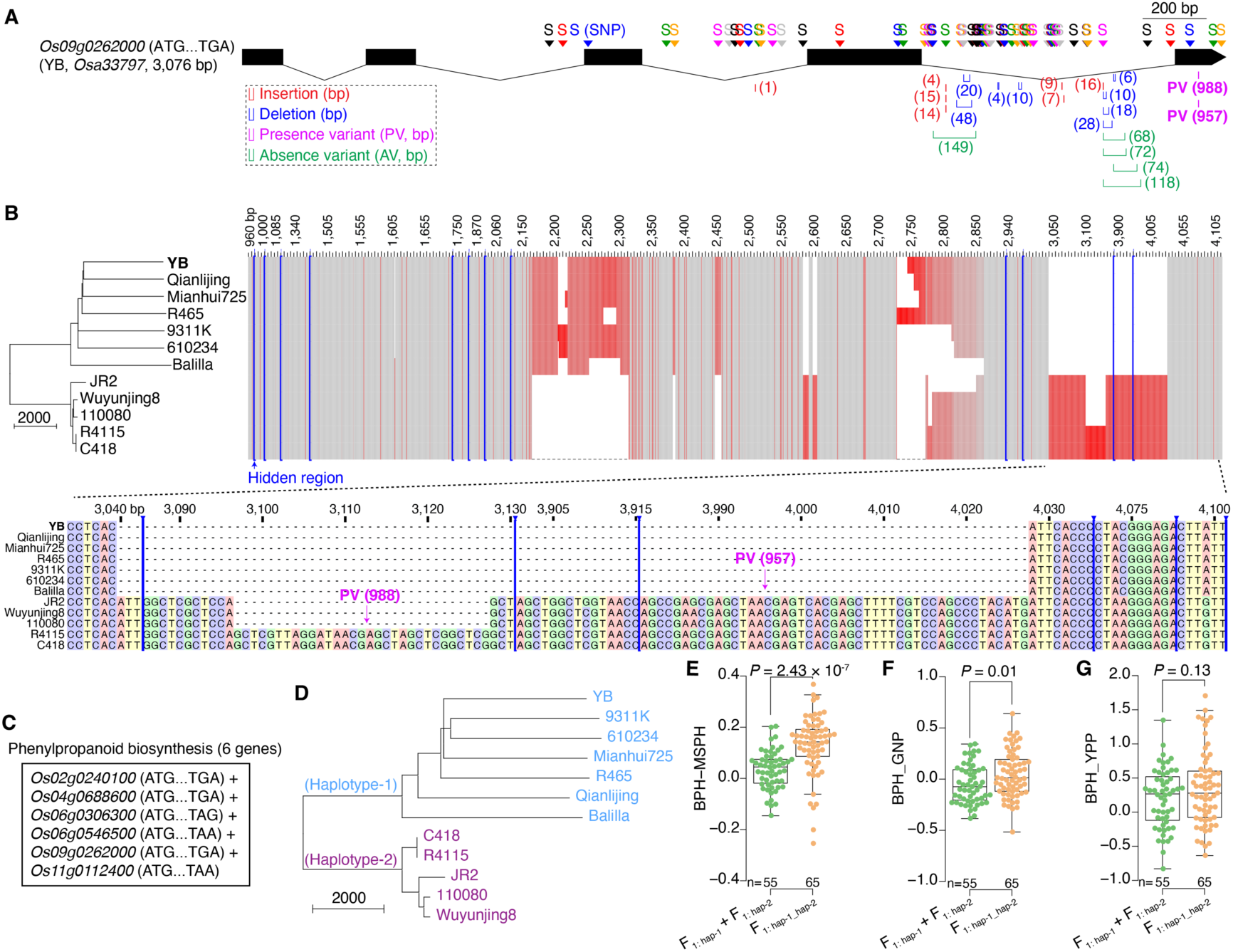
Genomic variants in phenylpropanoid biosynthesis genes contribute to heterosis for plant height. **(A)** Gene structure of *Os09g0262000*. With Yuetai B (YB) as the reference, genomic variants (SNPs, insertions, deletions, presence variants (PVs), and absence variants (AVs)) detected in 11 inbred lines are indicated. **(B)** Clustering of 12 inbred lines based on sequence variants of *Os09g0262000*. Red blocks represent the presence of variants. Detailed sequences of two PVs and two SNPs in the fifth exon are shown. **(C)** Six phenylpropanoid biosynthesis genes. **(D)** Haplotypes of 12 inbred lines based on sequence variants of the six genes. **(E-G)** Comparisons of better-parent heterosis for maturation stage plant height (BPH_MSPH; **E**), grain number per panicle (BPH_GNP; **F**), and grain yield per plant (BPH_YPP; **G**) between F_1_ hybrids with heterozygous (haplotype-1_haplotype-2) versus homozygous haplotypes (haplotype-1 and haplotype-2). *P* values indicate independent samples *t* test.

To investigate associations of parental genomic variants and different inheritance patterns, we used the top 50 HAMs per trait based on the established predictive models. Additive and partially dominant effects correlated more strongly with heterosis of the 17 traits than dominant effects (Supplemental Figure 40). Importantly, additive and partially dominant effects showed stronger associations with the number of parental unique variants across all variant types (Figure 7A-D). Overall, parental genomic variants were tightly associated with both heterosis and additive/partially dominant effects.

**Figure 7.**
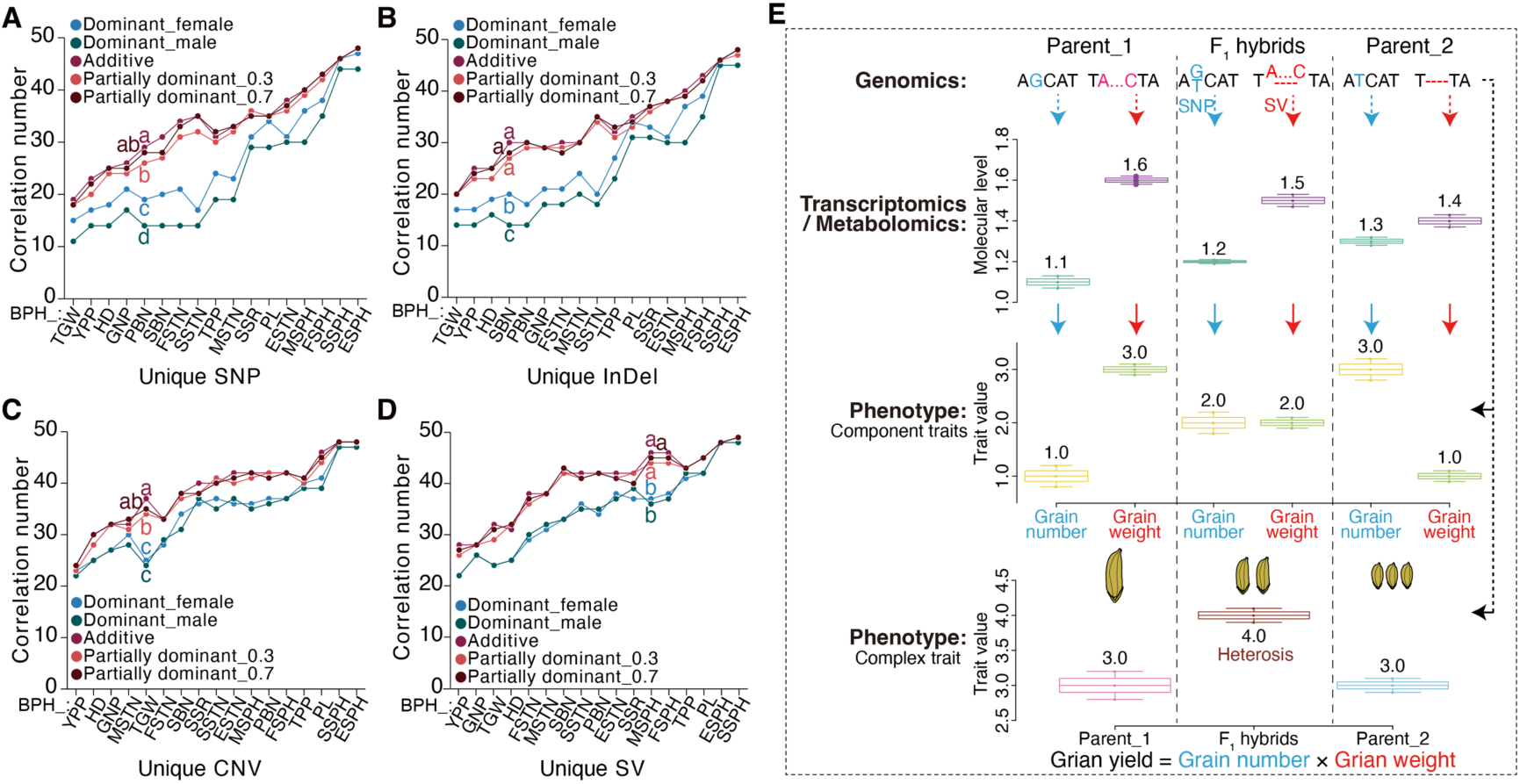
Associations of parental multi-omics variation, different inheritance patterns, and heterosis of component/complex traits in rice. **(A-D)** Number of significant correlations between parental unique variants and the top 50 heterosis-associated metabolites per trait (with different inheritance patterns). Analysis of variance (weighted) was performed with the least significant difference used in post hoc test. **(E)** A model of additive effect from genomic variants on yield heterosis in rice. To simplify interpretation, grain yield is determined by grain number and grain weight, with SNP and SV related with each trait, respectively. Due to additive effect from genomic variants in parents, the F_1_ hybrids showed intermediate transcriptomic and metabolomic levels. Accordingly, in F_1_ hybrids displayed mid-parent values for grain number and grain weight. With multiplicative interaction of the two yield components, positive yield heterosis occurred in F_1_ hybrids, with equal yield in parents. Abbreviations: better-parent heterosis = BPH, yield per plant = YPP, secondary branch number = SBN, grain number per panicle = GNP, tiller number per plant = TPP, seed setting rate = SSR, heading date = HD, thousand grain weight = TGW, seedling stage tiller number = SSTN, elongation stage tiller number = ESTN, flowering stage tiller number = FSTN, maturation stage tiller number = MSTN, seedling stage plant height = SSPH, elongation stage plant height = ESPH, flowering stage plant height = FSPH, maturation stage plant height = MSPH, panicle length = PL, and primary branch number = PBN.

Finally, with consistent transcriptomic evidence for rice heterosis (Dan et al., 2024b), we proposed a model integrating genomic, transcriptomic, and metabolomic variation to illustrate how the main inheritance patterns (e.g., additive effect) contribute to heterosis of component and complex traits in rice (Figure 7E). For simplicity, grain yield was modeled as a function of grain number (SNP-associated) and grain weight (SV-associated), assuming equal parental yield. Genomic variants drove differential gene expression and metabolite levels in parents (Parent_1 and Parent_2). Due to additive effect, the F_1_ hybrids yielded intermediate transcriptomic and metabolomic levels. Since trait values scaled linearly with molecular levels, grain number and grain weight in F_1_ hybrids also displayed mid-parent values. The additive effects of two yield components, which differed reciprocally in parents, then generated positive yield heterosis in F_1_ hybrids.

## DISCUSSION

The genetic mechanisms underlying heterosis have been debated for over a century without definitive consensus. The mysterious molecular bases of heterosis may stem from the multitude of molecules associated with heterosis, including genes, metabolites, and proteins (Birchler et al., 2006; Birchler et al., 2010; Huang et al., 2016; Schnable and Springer, 2013). Accurate identification of these molecules remains challenging, hinders mechanistic insights; consequently, few heterosis-associated genes with strong function have been reported by far (Krieger et al., 2010; Sun et al., 2023).

To highlight genetic factors in rice heterosis, we minimize environmental effects by employing uniform growth conditions for phenotyping seedling length. This allows precise dissection of genotype- and developmental stage-dependent effects on gene expression and metabolite levels, facilitating the successful screening of heterosis-associated molecules from parental differential molecules. We identify and validate the function of three genes to heterosis for seedling length. However, only one of the three genes also plays a role in heterosis for plant height at the maturation stage. Since these heterosis-associated molecules form tightly connected networks, and specific modules within these networks explain a significant proportion of heterosis variance. Thus, modulating entire pathways (e.g., phenylpropanoid biosynthesis) or key network modules (e.g., Module_1 and Module_2 of Network_HLN), rather than targeting single gene, through genetic engineering technology may enable the precision establishment of heterosis across stages and environments. Furthermore, external application of specific metabolites to F_1_ hybrids at appropriate developmental stages may enhance the degrees of heterosis without genomic alteration, given the significant correlations between the levels of heterosis-associated metabolite and heterosis magnitude.

Additive and partially dominant effects represent the predominant inheritance patterns for heterosis-associated molecules. Notably, the additive effect, corresponding to mid-parent values, exhibits predictability comparable to hybrid profiles in both rice and maize, highlighting its application potential for predicting heterosis of F_1_ hybrids in breeding programs. Indeed, additive effect has predicted trait values and heterosis of diverse agronomic traits for F_1_ hybrids across hybrid populations, developmental stages, and growth environments in rice (Dan et al., 2024a; Dan et al., 2021; Dan et al., 2019; Dan et al., 2020; Dan et al., 2016). However, building robust predictive models for heterosis of complex traits like yield remains challenging. Several key issues should be considered to ensure accuracy of predictive models. Firstly, the accessions used for modeling must capture sufficient genetic diversity to avoid reduced predictability from distant genetic relatedness. Then, for low heritability traits (e.g., tiller number, yield), omics data derived from relevant tissues (e.g., young tillers, young panicles) will ensure accurate predictions (Zhou et al., 2019b). Apart from transcriptomic and metabolomic profiles used in this study, adding other quantitative omics data, including proteomics, ionomics, and metagenomics (Kakoulidou et al., 2024; Price et al., 2017; Xu et al., 2021; Zhang et al., 2023), can improve predictability. Accounting for the important roles of environmental effects on heterosis, predictive models need to integrate phenotypic data from multiple environments (Liu et al., 2021a). Additionally, rapid advances in data processing ability from machine learning and artificial intelligence will further enable robust model development using single or combined strategies.

Unlike classical heterosis models emphasizing modes of genomic loci (Bruce, 1910; Crow, 1948; Davenport, 1908; East, 1936; Jones, 1917; Li et al., 2015; Li et al., 1997; Powers, 1944; Xiao et al., 2021; Yu et al., 1997), our approach focuses on downstream omics data to provide quantitative insight into heterosis. Beyond demonstrating contributions of additive/partially dominant effects at transcriptomic/metabolomic levels, we bridge associations among parental genomic variants, inheritance patterns of heterosis-associated molecules, and heterosis of component/complex traits. Using heterosis for plant height as an example, we detect sequence variants (particularly SVs) in phenylpropanoid biosynthesis genes that cluster parents into distinct haplotypes. The F_1_ hybrids with heterozygous haplotypes exhibit stronger heterosis than those with homozygous haplotypes, suggesting an overdominance model underlying rice heterosis.

The genetic mechanisms of heterosis for yield or other complex traits are more intricate than to heterosis for seedling length/plant height. Given that the F_1_ hybrids with heterozygous haplotypes enhanced heterosis not only for plant height but also grain number (while reducing heterosis for heading date), we speculate that phenylpropanoid biosynthesis genes may interact with known heterosis-regulating genes like *Ghd7* (Wang et al., 2022), *Ghd8* (Li et al., 2016; Sun et al., 2023), and *Hd1* (Zhou et al., 2021b) to influence yield heterosis. Moreover, large-size genomic variants, especially presence variants, play pivotal roles in haplotype divergence and affect heterosis of multiple traits. Thus, assembling parental genomes and accurately identifying SVs will provide a more comprehensive genetic landscape of plant heterosis.

In summary, our study provides multi-omics evidence that additive and partially dominant effects constitute the predominant inheritance patterns underlying heterosis, irrespective of genotype, trait/phenotype, hybrid population, developmental stage, growth environment, omics data type, or even species. Future efforts should leverage accurately assembled pan-genomes and multi-omics integration to dissect the interplay between large-effect genomic variants and the main inheritance patterns in heterosis of complex traits, which will improve the breeding efficiencies of hybrid crops.

## METHODS

### Plant materials and field growth conditions

According to the *indica–japonica* indices of 223 inbred lines (ILs) evaluated by 33 pairs of InDel markers (Lu et al., 2009; Yao and Huang, 2013), 17 ILs, ranging from *indica* to intermediate types to *japonica*, were selected as parents of a hybrid population using a complete diallel cross design (Dan et al., 2016). Moreover, 211 recombinant inbred lines (F_5_ followings) derived from F_1_ individuals of the 223 ILs were crossed with a Honglian-type cytoplasmic male sterile line Yuetai A. Plants of the diallel and test cross populations were planted with different density as previously described at the Hybrid Rice Experimental Base of Wuhan University in Ezhou (Hubei Province, China) in 2012 and 2015 (Dan et al., 2019), respectively.

For the diallel cross population, ten experimental plants were planted with 16.5 cm between plants and 26.4 cm between rows, and two plants of the sterile line, namely Yuetai A, were planted on two sides of each row. To obtain F_1_ hybrids, the young panicles of the female parents that would flower in the next few days were immersed in warm water at 44 °C (for *indica* or near-*indica* ILs) or 46 °C (for *japonica* or near-*japonica* ILs) for 5 min to sterilize the pollens. For the hand-pollination procedure, three to four panicles were combined in a paper bag, and one to two bags from each female parent without pollination were used as controls. For female parents with a high self-fertilization rate, such as R465 and Qianlijing, the pistils in each spikelet were manually removed and pollinated by hand. Three biological replicates of each genotype were randomly distributed in the field. The middle plants of each row, ranging from three to ten plants, were selected for the collection of phenotypic data for 17 agronomic traits. Plant height, measured from the base of the plant to the highest part of the plant using meter rulers, was recorded from the seedling to mature stages. The four time points included 28 days after transplanting (8 July, 2012), 42 days after transplanting (22 July, 2012), the time of flowering, and the time of maturation, which were defined as the seedling stage plant height, elongation stage plant height, flowering stage plant height, and maturation stage plant height, respectively. The tiller number per plant was also counted together with the plant height at four different stages, which were defined as the seedling stage tiller number, elongation stage tiller number, flowering stage tiller number, and maturation stage tiller number. The heading date was the number of days between the heading time when a plant from a biological replicate started flowering and the date of sowing (13 May, 2012). At maturity, all panicles from each plant were harvested and dried in the sunshine for approximately one week and then dried in ovens at 42 °C for two days before yield and yield-related traits were measured. Panicle length was measured from the panicle neck to the top for all panicles from each plant. The primary branch number and secondary branch number per panicle were counted manually for all panicles. Panicles from each plant were counted and treated as the tiller number per plant. After manually removing seeds (with or without grains) from panicles for each individual plant, empty seeds and seeds with grains were mechanically separated and counted using seed counters (PME-1, Shanke Equipment, Shanghai). The grain number per panicle was calculated by dividing the total seed number (sum of empty seeds and seeds with grains) of a plant by the tiller number per plant. The seed setting rate was calculated as the ratio of the number of seeds with grains to the total number of seeds. The weight of 1,000 grains from a plant was defined as the thousand-grain weight, and the weight of the seeds with grains per plant was defined as the grain yield per plant. Due to low germination rate of several hybrid seeds, phenotypic data of 245 (expected 272) F_1_ hybrids were used for analyses.

For the test cross population (211 F_1_ hybrids), each biological replicate contained ten plants (per row) spaced at 16.5 cm × 16.5 cm apart. Three biological replicates of each genotype were placed in three consecutive rows and randomly planted in the field. The phenotypic data of four agronomic traits, namely, heading date, thousand-grain weight, maturation stage plant height, and yield per plant were collected. The heading date was recorded for one biological replicate. The middle five plants per row were selected for measurement of plant height at the maturation stage. Panicles from the five plants were bagged and used to collect data on grain yield per plant and thousand-grain weight. Moreover, eight agronomic traits, namely, heading date, thousand grain weight, panicle length, seed setting rate, grain number per panicle, maturation stage plant height, grain yield per plant, and tiller number per plant, of the 17 ILs and 36 F_1_ hybrids from the diallel cross population were recollected along with the test crosses. Parts of trait values of the diallel and test cross populations have been reported in previous studies (Dan et al., 2021; Dan et al., 2019; Dan et al., 2020; Dan et al., 2015; Dan et al., 2016; Dan et al., 2014), and the complete datasets for all investigated traits are provided in source data.

In the functional validation of three heterosis-associated genes for seedling length/plant height, seedlings of Yuetai B, Zhonghua11, Zhonghua11 mutants, and wild-type/mutant F_1_ hybrids were planted in an experimental field at Wuhan University in 2023. The seedlings per genotype were planted in two consecutive rows with a 16.5 cm spacing. To minimize marginal effects, rows with fewer than ten seedlings were bordered by the male sterile line Yuetai A. Plant height was measured from the middle five plants of each row.

### Planting of rice seedlings for phenotypic and omics analysis

Two batches of rice seedlings were planted for untargeted metabolite profiling analysis. The first batch in 2016 (Batch_2016) included 17 ILs, 107 RILs selected from the 211 RILs, and the cytoplasmic male sterile line Yuetai A. As previously described (Dan et al., 2020), the germinated seeds were sown in soil with a spacing of 2 × 2 cm. The seedlings were grown at 30 °C under a 16 h light/8 h dark photoperiod in a light incubator. Two biological replicates were established for each genotype. On the 15^th^ day, the aboveground parts, excluding approximately 0.5 cm of tissue close to the soil, of approximately ten plants were harvested and washed with ddH_2_O three times. Two biological replicates of the 17 ILs were harvested as two experimental samples for each genotype. Two biological replicates of the 107 RILs and Yuetai A were pooled as one experimental sample for each genotype. The seedlings were first placed in 5 mL tubes (Eppendorf #0030119460) and snap frozen in liquid nitrogen.

The second batch in 2018 (Batch_2018) consisted of ten pairs of hybrid combinations. Seeds of 6 ILs and ten corresponding reciprocal F_1_ hybrids were first dehulled and sterilized with 75% ethanol (1 min) and 0.15% HgCl₂ (10 min). The seeds were then transferred to sterilized half-strength Murashige and Skoog rooting medium (pH = 5.8) supplemented with phytogel. Ten seeds were sown on the surface of 30 mL of medium per glass tube (diameter: 3 cm and height: 20 cm). Three to six biological replicates were set up for each genotype or the same genotype at different time points. Seedlings were grown at 28 °C under 24-hour light in an artificial climate chamber (Zhu Laboratory at the College of Life Sciences, Wuhan University). The seedling length and weight of a pair of hybrid combination containing two ILs (Balilla and Yuetai B) and the corresponding reciprocal F_1_ hybrids were recorded on 5, 10, and 15 days after sowing. Five seedlings from each glass tube were removed from the medium and photographed for subsequent measurement of seedling length using ImageJ (version 1.5.3). After excluding the roots, the seedling bases were washed three times with ddH_2_O and the water on the surface was removed with absorbent paper before weighing. The five seedlings were snap frozen in liquid nitrogen, and photographs and weights were taken and used for subsequent experiments. The remaining nine pairs of hybrid combinations were harvested for seedling length measurements after 15 days of growth. The sealing films on the glass tubes were removed on day 11 and 5 mL of ddH_2_O was added to each tube for normal growth. All the experimental samples were stored at -80 °C. Parts of seedling samples from Batch_2018 were also used to perform RNA-sequencing analysis.

Using the same experimental procedures as Batch_2018, seeds of Yuetai B, Zhonghua11, mutated lines of Zhonghua11 (T_4_), and their wild-type /mutant F_1_ hybrids were sown in glass tubes. Following a laboratory move to the Dangdai Building at Wuhan University, the seedlings of two ILs and their wild-type F_1_ hybrids were first grown at 28 °C in a new artificial climate chamber under 24-hour light or in an intelligent breeding room under a 16-hour light and 8-hour dark photoperiod. The two ILs, mutated lines, and wild-type/mutant F_1_ hybrids were then cultivated in the artificial climate chamber at 28 °C under 24-hour light. The shelf surface placing glass tubes was heated to approximated 35 °C for 12 days. Seedling length was recorded for 15 plants per genotype after 15 days.

### RNA sequencing

Seedlings from each biological replicate were ground into fine powder using mortar and pestle with liquid nitrogen. After total RNA was extracted with TRIzol (Invitrogen), the RNA was first assessed by electrophoresis in 1% agarose gels to check for degradation and contamination states. The concentrations (Qubit2.0 Fluorometer) and integrity (Agilent 2100 bioanalyzer) of each RNA sample were analyzed. DNA library preparation (NEBNext Ultra RNA Library Prep Kit, NEB) with index codes and sequencing experiments were performed on the sequencing platform of Novogene Co., Ltd (Beijing, China). Briefly, mRNA was first purified from 3 μg of total RNA using magnetic beads attached to poly(T) oligo. The mRNA with poly(A) tails was randomly fragmented (NEBNext First Strand Synthesis Reaction Buffer) and used to synthesize the first strand of cDNA with the addition of hexamer primers and M-MuLV reverse transcriptase. After the RNA was degraded with RNase H, the second strand of cDNA was synthesized using DNA polymerase I with dNTPs. The strands were then converted to blunt ends using the activities of T4 DNA polymerase and DNA polymerase I. The products were subjected to adenylation, and NEBNext adaptors were added to the products. Approximately 200 bp of cDNA was purified using AMPure XP beads (Beckman Coulter) and treated with 3 μL of USER Enzyme (NEB) at 37 °C for 15 minutes and 95 °C for 5 minutes. PCR was then performed using Phusion High-Fidelity DNA polymerase with universal and index primers. The reaction products were repurified with AMPure XP beads, and the quality of the library was assessed using an Agilent Bioanalyzer 2100. After clustering of the index-coded samples using the cBot Cluster Generation System (TruSeq PE Cluster Kit v3-cBot-HS, Illumina), the libraries were sequenced on the Illumina Hiseq PE150 platform, and 150 bp paired-end reads were acquired with more than 6 G per sample.

The raw reads were first processed by removing adapters, poly N and low-quality reads. The Q20, Q30, and GC contents were calculated for the clean reads. The clean reads were then aligned to the Nipponbare reference genome (Oryza_sativa_Ensembl_43) using Hisat2 (v2.0.5) (Kim et al., 2015) and the numbers of gene reads were counted using featureCounts (v1.5.0-p3) (Liao et al., 2014).

### Normalization and differential analysis of transcriptomics data

Normalization and differential analysis of the transcriptomics data (42,341 genes) for the hybrid pair of Balilla and YB at 5, 10, and 15 DAS were performed using NetworkAnalyst 3.0 (Zhou et al., 2019a) with the following parameters: specify organism, not specified; data type, bulk RNA-seq data (counts); ID type, not specified; gene-level summarization, sum; variance filter, 30; low abundance, 30; filter unannotated genes, yes; normalization, Log2-counts per million; statistical method, DESeq2; Log2 fold change, 2; and adjusted *p-*value, 0.0001. As the numbers of differentially expressed genes between reciprocal F_1_ hybrids of Balilla and YB at 5, 10, and 15 DAS were only 1, 13, and 29, respectively, the differences between reciprocal F_1_ hybrids were not included in the analysis. To analyze the differentially expressed genes of the six ILs, the data matrix containing 42,341 genes was separately normalized (sum, none, and autoscaling) with MetaboAnalyst 4.0 (Chong et al., 2018) and analyzed with ANOVA (adjusted *p*-value = 0.05, post-hoc analysis = Fisher’s LSD). Principal component analysis (PCA) of the transcriptomic and metabolomic data, which were normalized data from NetworkAnalyst 3.0 (Zhou et al., 2019a), was performed at ClustVis (https://biit.cs.ut.ee/clustvis) (Metsalu and Vilo, 2015).

Two-way ANOVA of the 5,844 DEGs was performed using the “Time Series Analysis” (time series and one factor) module of MetaboAnalyst 4.0 (Chong et al., 2018) with the following parameters: ANOVA type, type Ⅰ; interaction, yes; adjusted *p*-value cut-off, 0.05; and correction for multiple testing, false discovery rate.

### Pathway enrichment analysis of the differentially expressed genes

The ClueGO (version 2.5.8) (Bindea et al., 2009) and CluePedia (version 1.5.8) (Bindea et al., 2013) plugins of Cytoscape (version 3.9.0) (Saito et al., 2012) were used for pathway enrichment analysis of the differentially expressed genes. Functional analysis was selected as the analysis mode. The up- and downregulated genes were represented by diamonds (shape 1) and triangles (shape 2), respectively. The GO (ellipse), KEGG (hexagon), and WikiPathways (V) were selected as ontologies/pathways. The GO term fusion was selected and only pathways with pV ≤ 0.01 were shown. The GO tree interval was between 3 (minimum level) and 5 (maximum level). The minimum number and percentage of genes were set to 3 and 4%, respectively, for both clusters. The genes accounting for more than 60% of a cluster were defined as cluster-specific genes. The kappa score of network connectivity was set to 0.4, and enrichment/depletion (two-sided hypergeometric test) with Benjamini‒Hochberg pV correction was set as the statistical option. GO term grouping was selected in the group options, and the leading term was based on the highest significance with fixed colors. The kappa score was selected for term grouping. The initial group size, percentage of genes for group merging and percentage of terms for group merging were set to 1, 50% and 50%, respectively.

### Untargeted metabolite profiling

For the seedling samples in Batch_2016, liquid chromatography-mass spectrometry (LC‒MS) analysis procedures have been previously described (Chen et al., 2024; Dan et al., 2020). In brief, all tissues in each EP tube were first ground into homogenized powder by mortar and pestle with the addition of liquid nitrogen. Eighty milligrams of powder were then transferred to 2 mL EP tubes and the tubes were vortexed for 1 min after the addition of 1 mL of precooled extraction solution (methanol/acetonitrile/water, 2/2/1). A quality control sample was prepared by mixing 10 μL of each experimental sample. The experimental and quality control samples were then subjected to two rounds of ultrasonic treatments (30 min each) and kept at -20 °C for 60 min. The samples were then centrifuged at 14,000 g for 15 min at 4 °C and the supernatants were transferred to clean tubes. After drying the supernatants in a vacuum concentrator, 100 μL of acetonitrile (acetonitrile/water, 1/1) was added to the tubes. The tubes were vortexed and centrifuged at 14,000 g for 15 minutes at 4 °C. The supernatants were transferred to sample tubes for LC‒MS analysis on the platform of Shanghai Applied Protein Technology Co., Ltd.

The experimental and quality control samples were separated on an Agilent 1290 Infinity LC system (Agilent Technologies) equipped with a reverse-phase T3 column (ACQUITY UPLC HSS T3, 1.8 μm, 2.1 mm × 100 mm, Waters). The injection volume was 2 μL per sample. The flow rate was 0.3 mL/min and the column temperature was 25 °C. Regents A and B in positive ion mode were 0.1% formic acid (in water) and 0.1% formic acid (in acetonitrile), respectively. In addition, 0.5 mM ammonium fluoride (Sigma‒Aldrich) and acetonitrile (Merck) were used in negative ion mode. The gradient elution steps were as follows: 0-1 min, 1% B; 1-8 min, linearly increased to 100% B; 8-10 min, held at 100% B; 10-10.1 min, reduced to 1% B; and 10.1-12 min, held at 1% B. Experimental samples were analyzed in a random order, and the quality control sample was injected every nine experimental samples. To obtain MS profiles in both positive and negative electron spray ion modes, the LC system was coupled to an Agilent 6550 iFunnel QTOF mass spectrometer (Agilent Technologies). The MS data were collected from 60 to 1,200 Da. The acquisition rate was four spectra per second, and the cycle time was 250 microseconds. The temperatures of the drying gas and the sheath gas were 250 °C and 400 °C, respectively. The flow rates of the drying gas and the sheath gas were 16 L/min and 12 L/min, respectively. The nebulizer pressure was set at 20 pounds per square inch (psi). The capillary, nozzle, and fragment voltages were 3,000, 0, and 175 V, respectively. Tandem mass spectrometry data were then acquired on the quality control sample using a TripleTOF 6600 mass spectrometer (AB SCIEX). The mass range was between 60 and 1,200 Da. To increase the data acquisition rate of the quality control sample, the sample was injected twice and the mass range was further divided into four sequential windows: 60-200, 190-400, 390-600, 590-1,200 (full range, 60-1,200) Da (for chemical standard library) and 200-600, 590-750, 740-900, 890-1,200 (full range, 200-1,200) Da (for lipid MS/MS library). The mass spectrometry accumulation time was 200 ms per spectrum, and the tandem mass spectra were obtained in an information-dependent manner with high sensitivity. Ion source gas 1, ion source gas 2, and the curtain gas were set at 40, 80, and 30 psi, respectively. The source temperature was 650 °C. The ion spray voltage floating were +5,000 V and -4,500 V, respectively. The declustering potential and collision energy were ± 60 V and 35 ± 15 eV, respectively. The number of candidate ions for monitoring per cycle was ten and the isotope isolation width was 4 Da. The MS/MS accumulation time was set to 50 ms.

The raw MS files (.d) of the experimental and quality control samples were converted to mzXML format using ProteoWizard (version 3.0.6150) (Chambers et al., 2012), and the transformed files were uploaded to XCMS (version 1.46.0) (Gowda et al., 2014) for feature detection, retention time (RT) correction and feature alignment with previously reported parameters (Jia et al., 2018). The 17 ILs formed one group, and the 107 RILs plus Yuetai A formed another group. The features with more than 50% nonzero values in at least one group were retained for further analysis. The error thresholds for m/z and RT were set at 15 ppm and 20 s for the alignment of MS and MS/MS data, respectively. The MS/MS data of the quality control sample were matched against both the local chemical standard library (Shen et al., 2019) and the lipid MS/MS library (Tu et al., 2017), and metabolites were annotated with a mass accuracy of less than 25 ppm. For the seedling samples in Batch_2018, the metabolomics analyses were the same as those in Batch_2016, except that the four sequential windows of the quality control sample were changed to three sequential windows: 60-300, 290-600, and 590-1,200 (full range, 60-1,200) Da. The Metlin library (Domingo-Almenara et al., 2018) was also used for feature annotation.

### Normalization and differential analyses of metabolomics data

The normalization of the metabolomics data for the hybrid combination of Balilla and YB at 5, 10, and 15 DAS followed the same parameters as those used for the transcriptomics data. ANOVA was used to analyze differential metabolites for the six ILs with a data matrix of 15,083 analytes. For ease of interpretation, the word “metabolite” was used in this manuscript to represent analytes with or without annotations. The “statistical analysis (metadata table)” module of MetaboAnalyst 5.0 (Pang et al., 2022) was used to perform two-way ANOVA on the 2,331 differential metabolites with genotype and stage as metadata. One of the three biological replicates of each genotype at each stage was used as a subject. Multiple factors were chosen for the study design. Further data filtering, sample normalization, data transformation, and data scaling were not performed. Type Ⅰ ANOVA was used for two-way independent ANOVA, and two-way repeated ANOVA was used to calculate interactions. The false discovery rate was used to correct for multiple testing, and the adjusted *P* value cut-off was set at 0.05.

To normalize the metabolomics data of the 17 ILs, Yuetai A, and 107 RILs, the intensities of the two biological replicates of the 17 ILs were first averaged and then normalized using MetaboAnalyst 5.0 (Pang et al., 2022) (sum, none, and autoscaling for sample normalization, data transformation, and data scaling). Metabolites with RSD greater than 25% in the QC samples were removed and the interquantile range was selected to filter the features (40%). For the differential metabolite analysis of the 17 ILs, the feature intensities of two biological replicates for each IL were used and ANOVA was performed after filtering 40% of the features and data normalization (sum, none, and autoscaling) using MetaboAnalyst 5.0 (Pang et al., 2022).

### Determination of inheritance patterns for heterosis-associated molecules

To determine the inheritance patterns for heterosis-associated molecules, transcriptomic and metabolic levels were compared for F_1_ hybrids, the corresponding parents, and the average levels of the corresponding parents via ANOVA using MetaboAnalyst 4.0 (Chong et al., 2018). Three biological replicates of reciprocal F_1_ hybrids from two ILs were combined into a hybrid genotype. An additive effect was assigned when the molecular levels of the hybrid genotype were not significantly different from the average levels of the corresponding parents but were significantly different from those of both parents. A partially dominant effect was assigned when the hybrid levels were significantly different from both the average parental levels and the molecular levels of the parent on the same side. A dominant effect was when the hybrid levels were not significantly different from those of the parent but were significantly different from the average levels of the two parents. An overdominant effect was the pattern when the hybrid levels were significantly higher or lower than those of the parent on the same side. The patterns of hybrid levels other than the above four types were defined as unclassified and marked as NA. To quantitatively describe the partially dominant effect (PD), the degrees for the molecules with partially dominant effect of the six hybrid combinations was calculated with the equation PD = (F_1_-P_2_)/(P_1_-P_2_), where F_1_, P_1_, and P_2_ represent the molecular levels of the F_1_ hybrids, parent_1, and parent_2, respectively. Several molecules that had a partially dominant effect in multiple F_1_ hybrids were used multiple times. The degree of the partially dominant effect was divided into two parts, with 0.5 as the threshold. The averages of the two groups were calculated and defined as two types of partially dominant effects (PD_0.3 and PD_0.7). Moreover, the equations PD_0.3 = P_m_+0.3×(P_f_-P_m_) and PD_0.7 = P_m_+0.7×(P_f_-P_m_) were used to represent partially dominant effects based on parental molecular levels.

### Pathway enrichment analysis of differential metabolites

The metabolomics data of the hybrid combination of Balilla and YB at 15 DAS were first used for pathway enrichment analysis. The table with normalized feature intensities and retention times (in seconds) was uploaded to the “functional analysis” module of MetaboAnalyst 5.0 (Pang et al., 2022). Both positive and negative modes were selected for analysis. In positive mode, the mass tolerance was set to 5 ppm. The interquantile range was selected for data filtering because the number of peaks was greater than 5,000. No data were used for sample normalization, data transformation, or data scaling. Both *mummichog* (Li et al., 2013) (*P* value cut-off: default top 10% peaks, version 2.0) and GSEA were used for enrichment analysis. As no pathway was significantly enriched for the differential metabolites between Balilla×Yuetai B and its parent Balilla with the default *P* value cut-off (0.01) in positive mode, the *P* value cut-off was increased to 0.05. The pathway library used was *Oryza sativa japonica* (Japanese rice) [KEGG]. Metabolite currency and adducts were set with the default sets, and pathways or metabolite sets containing at least 3 entries were set. The parameters for the negative mode analysis were the same as those for the positive mode analysis, except for that no data filtering was performed and the *P* value cut-off was set to 0.05. Putatively annotated metabolites and enriched pathways were obtained in each analysis. For the pathway analysis of the 645 DMs using the “Pathway Analysis” module of MetaboAnalyst 5.0 (Pang et al., 2022), feature annotation information was obtained from the results of differential metabolites between Balilla and YB at 15 DAS. The parameters were set as follows: scatter plot, visualization method; hypergeometric test, enrichment method; relative-betweenness centrality, topology analysis; all compounds in the selected pathway library, reference metabolome; and *Oryza sativa japonica* (Japanese rice) (KEGG), pathway library. Three pathways were significantly enriched based on the annotation of eight peaks in positive mode. Moreover, 20 metabolites (23 peaks from both ion modes), which were annotated with local chemical standards, lipids, and Metlin libraries, from the three significantly enriched pathways were also used for analysis. Correlations between the metabolite levels of 26 peaks and seedling length were determined. One peak (M147T118_POS) was annotated as 4-hydroxycinnamic acid in both the local standard library and the result of the pathway enrichment analysis, and four peaks had no concentration in the normalized metabolomics data.

### Integration of genes and metabolites into networks

The normalized transcriptomics and metabolomics data were used to construct networks via OmicsAnalyst (Zhou et al., 2021a). No organism or ID type was specified in the annotation procedure. Features with 50% missing values were excluded and the remaining missing values were replaced by LoDs. The filter threshold for variance and abundance was set to 0. In the comparison step, the genotype was used for grouping, and no data transformation was performed. ANOVA was used for the differential analysis. The fold change cut-off was set at 2.0, and the *P* value cut-off was set at 0.01. To make the data suitable for inter-omics analysis, both the transcriptomics and metabolomics data underwent normalization by sum, variance stabilizing normalization, quantile normalization and range scaling. The statistically significant features were then used to construct an interomics correlation network using the Spearman method. The correlation threshold was set at 0.8, and both positive and negative correlations were included. For Network_51015, 626 genes and 645 metabolites were used, and a total of 213 genes and 198 metabolites were statistically significant. The network contained three subnetworks, and the largest subnetwork with 2,138 edges (subnetwork_1, 61.8% and 38.2% positive and negative correlations, respectively) had 211 genes and 196 metabolites as nodes. The other two subnetworks had two interactions. For Network_HLN, 967 genes and 990 metabolites were used for analysis, and 103 genes and 158 metabolites were retained in the network. Network_HLN contained seven subnetworks, and the largest subnetwork with 868 edges (subnetwork_1) contained 97 genes and 149 metabolites. The remaining six subnetworks had nine edges. The walktrap algorithm was used for module detection. Subnetwork_1 of Network_51015 contained seven modules. The module size and *P* value of each module were 150 (*P* = 2.34×10^-21^, Module_1), 105 (*P* = 1.01×10^-14^, Module_2), 80 (*P* = 2.29×10^-12^, Module_3), 21 (*P* = 0.00303, Module_4), 13 (*P* = 0.452, Module_5), 7 (*P* = 1, Module_6), and 6 (*P* = 0.318, Module_7), respectively. Subnetwork_1 of Network_HLN contained eight modules. The module size and *P* value of each module were 112 (*P* = 6.6×10^-35^, Module_1), 37 (*P* = 2.61×10^-7^, Module_2), 35 (*P* = 1.39×10^-6^, Module_3), 16 (*P* = 3.8×10^-5^, Module_4), 11 (*P* = 0.111, Module_5), 7 (*P* = 0.0592, Module_6), 6 (*P* = 0.0176, Module_7), and 6 (*P* = 0.0309, Module_8), respectively. The subnetworks were imported into Cytoscape (Saito et al., 2012) for display using the compound spring embedder layout. The node size was determined by the node degree.

### Joint omics analysis of differentially expressed genes and metabolites

To integrate the transcriptomics and metabolomics results, the MSU IDs of the 626 DEGs were first converted to gene symbols using the NCBI “Batch Entrez” tool (https://www.ncbi.nlm.nih.gov/sites/batchentrez). Genes with symbols (340 genes) in the *Oryza sativa japonica* group were uploaded to the “Joint pathway” analysis module of MetaboAnalyst 5.0 (Pang et al., 2022). Moreover, the annotation information of the 645 peaks was extracted from the results of the pathway enrichment analysis between Balilla and YB at 15 DAS. The KEGG IDs for 29 compounds in positive mode and 44 compounds in negative mode were uploaded as metabolomics data. “All pathways (integrated)” was selected as the pathway database. The parameters for enrichment analysis, topology measure, and integration method were set as hypergeometric test, degree centrality, and combine queries, respectively.

### Analyses of transcriptomic/metabolic/phenotypic data for inbred lines and F_1_ hybrids in maize and *Arabidopsis*

The field fresh weight, grain dry matter yield, and grain dry matter content of maize were obtained from published data in two previous studies (de Abreu et al., 2017; Schrag et al., 2018). Metabolite profiles of 3.5-day-old roots and leaves at the third leaf stage were also collected for 210 Dent and Flint lines and 363 F_1_ hybrids in the two studies (de Abreu et al., 2017; Schrag et al., 2018). The root and leaf metabolic datasets contained levels of 165 and 81 metabolites, respectively. Since metabolic data were not available for the parents of some of the F_1_ hybrids, the field fresh weight of 326 F_1_ hybrids, which were crossed between 24 Dent lines and 25 Flint lines, were used for analysis. No further data normalization was performed on the metabolic data. As reported in the study (de Abreu et al., 2017), the field fresh weight of the 326 F_1_ hybrids was recorded in two distinct field tests in 2010 (146 F_1_ hybrids) and 2012 (180 F_1_ hybrids). In the PLS analysis, the number was set to 100 to ensure a sufficient number of latent factors, and the number was defined for each regression analysis when the R^2^ value (adjusted) peaked. The R^2^ values (adjusted) corresponding to different numbers of latent factors and the greatest R^2^ values for hybrid profiles and different inheritance patterns were used for comparison analysis via ANOVA. Among the 326 F_1_ hybrids, 291 F_1_ hybrids had recorded data on grain dry matter yield and grain dry matter content in approximately seven agro-ecologically diverse environments (Schrag et al., 2018). The root metabolic data were used for PLS analysis, and the R^2^ values (adjusted) were used for comparison analysis.

The grain yield per plant (in three environments) of 12 maize inbred lines and 66 F_1_ hybrids from the inbred lines with a half-diallel design reported in a previous study (Wang et al., 2023) was used for analysis. The expression levels of 27,608 genes, which were pan and core genes in Zheng58, from the 12 inbred lines were also obtained from the study (Wang et al., 2023). As described in the study (Wang et al., 2023), each genotype of the 12 maize ILs had two sets (not biological replicate) of RNA-seq data, with one set containing equally mixed RNAs from eight tissues and the other from five tissues at different developmental stages. The expression data were processed with MetaboAnalyst 5.0 (Pang et al., 2022), with 40% set as the percentage for filtering. No data transformation, normalization, or scaling was performed. In total, 84 DEGs were identified in the 12 inbred lines by ANOVA, with Fisher’s LSD used in post hoc analysis (*P* value cut-off = 0.05). The average values of the two sets of RNA sequencing data were used to represent the levels of each inbred line. The 84 genes were classified according to the previously reported results (Wang et al., 2023). The number of latent factors was set to 50 in the PLS analysis of better-parent heterosis for grain yield per plant, mid-parent heterosis for grain yield per plant, and grain yield per plant.

The phenotypes of cotyledon and the first true leaf and transcriptomics data in *Arabidopsis* were obtained from a previously published study (Liu et al., 2021b). The transcriptomics data were normalized with above mentioned parameters in NetworkAnalyst 3.0. Differential expression genes between Col-0 and Per-1 at different stages were analyzed with the limma method, with Log2 fold change at 2 and adjusted *p-*value at 0.0001. The pooled differential expression genes across stages were treated as differential expression genes for each tissue. Two-way ANOVA on the normalized levels of DEGs with genotype and stage as metadata was implemented with MetaboAnalyst 5.0 (Pang et al., 2022). The overlapped DEGs of genotype, stage, and their interaction were used for further analysis. Pearson correlation between expression levels of the DEGs and phenotypes of cotyledon and the first true leaf was performed to screen the genes and the significance level was set at 0.05.

### Genome sequencing and analysis of genomic data

Seedlings from the 17 rice ILs were ground into fine powder using liquid nitrogen. The genomic DNA was extracted, and the quality and concentration were assessed by 1% agarose gel electrophoresis and Qubit 3.0 fluorometer. The DNA libraries (with index codes) were prepared and sequenced on the Illumina sequencing platform of Novogene Co., Ltd (Beijing, China). In brief, genomic DNA was randomly fragmented into ∼350 bp size via sonication (Covaris). The strands were then added with adapters and used for PCR amplification. Products were purified with AMPure XP beads (Beckman Coulter) and the concentrations were measured with Agilent 2100 Bioanalyzer. Libraries were sequenced on the Illumina PE150 platform, and the raw data were recorded in FASTQ format (> 15 G per sample). After the clean reads were obtained, the sequences were aligned to the reference genome using the Burrows‒Wheeler Aligner (mem -t 4 -k 32 -M, version 0.7.8-r455) (Li and Durbin, 2009). Dislodged duplications in the BAM files were removed via SAMtools (rmdup, version 1.3.1) (Li et al., 2009). SNPs and InDels in each sample were called with the parameters “-C 50 - mpileup -m 2 -F 0.002 -d 1000” and further filtered to reduce errors (depth of variant > 4 and mapping quality > 20). CNVs were detected by CNVnator (-call 100, version 0.3) (Abyzov et al., 2011), and SVs were detected by BreakDancer (depth of variant > 2, version 1.4.4) (Chen et al., 2009). Variant annotation was implemented with ANNOVAR (Wang et al., 2010). Genomic variants on chromosomes were visualized by Circos (0.69-9) (Krzywinski et al., 2009). To analyze the population structures of the 17 ILs, GATK (version 4.2.0.0) was first used to filter SNPs (QD < 2, FS > 60, MQ < 40, MQRankSum < -12.5, ReadPosRankSum < -8) and InDels (QD < 2, FS > 200, ReadPosRankSum < -20). SNPs and InDels were further filtered with VCFtools (maximum missing value < 10%, minus allele: 0.05, minimum read depth > 4, multiple alleles removed, version 0.1.17) and analyzed with Plink (version 1.90) and Admixture (version 1.3.0). When comparing the SNPs and InDels between each pair of samples of the 17 ILs, the variants were defined as common variants if they shared the same values for five parameters, including CHROM (chromosome), POS (position of the reference), REF (reference base), ALT (alternative base) and GT (genotype). Differential variants were those that were unique to each IL. For CNVs and SVs, variants were defined as common variants if they shared the same values of CHROM, POS1, POS2, TYPE and SIZE. The numbers of common and unique variants were counted, and the sums of unique genomic variants between each pair of ILs were used to represent the genetic distance. Phylogenetic trees (neighbour-joining method) based on one or multiple types of genomic variants in the 17 ILs were generated with MEGA (version 10.2.6) (Stecher et al., 2020).

### Phylogenetic analysis of phenylpropanoid biosynthesis genes

Using genome assemblies from 12 rice inbred lines and available gene family information (Dan et al., 2024b), we extracted sequences for eight of the ten heterosis-associated genes of seedling length, except for *Os04g0688500* and *Os07g0677100*. Among these eight genes, *Os01g0293900* and *Os02g0240300* were not detected in some inbred lines and thus excluded. Consequently, sequences spanning the start-to-stop codons of the six remaining genes were used for analyses. With sequences of YB as the reference, sequence variants were identified for each accession. Exon-intron structure of each gene was visualized using the Exon-Intron Graphic Maker (http://wormweb.org/exonintron) with variants annotated. Phylogenetic trees for individual and concatenated genes were built in Jalview (version 2.11.4.1) (Waterhouse et al., 2009), using E-INS in mafft method. Haplotypes of the 12 inbred lines were classified based on tree topology.

### Partial least squares regression analysis

Partial least squares regression (PLS), which has inherent ability to handle noisy, collinear, and incomplete variables in both predictors and responses (Eriksson et al., 1995; Wold et al., 2001), was applied to build multivariate models for screening heterosis-associated molecules and predicting heterosis of agronomic traits or hybrid phenotypes. To extract enough latent factors, the maximum number was initially set to 100 in SPSS 20. The optimum model for each trait was determined by selecting the highest adjusted R^2^ value, which also served as the measure of predictability. The mean variable importance in projection across all latent factors were used to estimate contributions of predictors, applying a threshold of 1 for screening heterosis-associated molecules of seedling length in rice. With class order taken into account, partial least squares-discriminant analysis (PLS-DA) was performed on F_1_ hybrids with different degrees of seedling length heterosis using MetaboAnalyst 5.0 (Pang et al., 2022). The analysis searched a maximum of five components and employed tenfold cross-validation. To assess model overfitting, the separation distance (B/W) method was used for permutation analysis (2,000 times). The predictability of PLS-DA model was evaluated using the Q^2^.

### Statistical analyses

Better-parent heterosis for each trait was calculated using the equation: BPH_trait = (F_1_ - P_high_)/P_high_, where P_high_ represents the high trait value of both parents. In the calculation of better-parent heterosis for heading date, parents with earlier heading dates were selected as P_high_. The calculated values were multiplied by -1 to represent the heterosis for heading date. Analysis of variance (least significant difference used in post hoc test), independent samples *t* test, paired samples *t* test, Spearman rank correlation, Pearson correlation (two-tailed, *P* value < 0.05) were performed using SPSS 20.0. Venn diagrams were drawn with the tool from http://bioinformatics.psb.ugent.be/webtools/Venn/. The phylogenetic tree of 223 inbred lines was based on genotypes and the tree was constructed using MEGA (Stecher et al., 2020) via the neighbour-joining method. The *indica* genotype, heterozygous genotype, and *japonica* genotype based on InDel markers were represented by 3, 2, and 1, respectively. A similarity matrix of the 223 inbred lines was then derived from MORPHEUS (https://software.broadinstitute.org/morpheus/) using Euclidean distance. For each box generated with BoxPlotR (Spitzer et al., 2014) (http://shiny.chemgrid.org/boxplotr/), the central line represents the median, the upper and lower edges indicate the 75th and 25th percentiles, respectively, and the whiskers are 1.5× the interquartile range away from the 75th and 25th percentiles. Source data are available with this paper in Supplemental Dataset 5.

## Abbreviations

ANOVA: analysis of variance
BPH: better-parent heterosis
CNV: copy number variation
DAS: days after sowing
DEG: differentially expressed gene
DM: differential metabolite
GO: gene ontology
HAG: heterosis-associated gene
HAM: heterosis-associated metabolite
IL: inbred line
InDel: insertions and deletion
MPV: mid-parent value
PC: principal component
PCA: principal component analysis
PLS: partial least squares
PLS-DA: partial least squares-discriminant analysis
RIL: recombinant inbred line
SNP: single-nucleotide polymorphism
SV: structural variation
YA: Yuetai A
YB: Yuetai B
YPP: yield per plant
SBN: secondary branch number
GNP: grain number per panicle
TPP: tiller number per plant
SSR: seed setting rate
HD: heading date
TGW: thousand grain weight
SSTN: seedling stage tiller number
ESTN: elongation stage tiller number
FSTN: flowering stage tiller number
MSTN: maturation stage tiller number
SSPH: seedling stage plant height
ESPH: elongation stage plant height
FSPH: flowering stage plant height
MSPH: maturation stage plant height
PL: panicle length
and PBN: primary branch number.

## DATA AND CODE AVAILABILITY

All data are available in the main text or the materials. The genomics and transcriptomics data reported in this study have been deposited in the NCBI database under BioProject accession PRJNA1122958. The metabolomics data have been deposited in the MetaboLights database under MTBL742 (Dan et al., 2020) and MTBL1344. Source data are available with this paper.

## ACKNOWLEDGEMENTS

The authors thank members of the Zhu lab for assistance in the collection of field phenotypic data. The authors thank Professor Baorong Lu (Fudan University) for the design of the diallel cross population. The authors thank Mr. Caibin Zhu for the fund of rice research. The authors thank Fulin Luo for the planting of rice. The authors thank Dr. Xiaofei Zeng for suggestions.

## AUTHOR CONTRIBUTIONS

Z. D. conceived the project and performed most part of the hybridizations. W. H. collected rice inbred lines. Z. D. and Yunping Chen collected phenotypic data of rice seedlings, prepared seedling samples for transcriptomics and metabolomics experiments, and analysed the omics data. Z. D., W. Z. and G. Y. constructed the diallel cross population and recorded phenotypic data of the hybrid population in 2012. Z. D. and W. H. screened the recombinant inbred lines and constructed the test cross population. Z. D., Y. X., J. H. and J. M. collected phenotypic data of the hybrid population in 2015. G. Y. performed the analysis of 223 inbred lines with InDel markers. Yi Chen participated in the construction of phylogenetic tree for the 223 inbred lines and the preparation of several heatmaps. Z. D., Yunping Chen, and W. H. wrote the manuscript.

## Supplementary information

Additional supporting information may be found online in the Supporting Information section at the end of the article.

Supplemental Figure 1-40.

Supplemental Table 1-3.

Supplemental Dataset 1-5.

## REFERENCES

Abyzov, A., Urban, A.E., Snyder, M., and Gerstein, M. (2011). CNVnator: an approach to discover, genotype, and characterize typical and atypical CNVs from family and population genome sequencing. Genome Res. 21:974–984.

Bindea, G., Galon, J., and Mlecnik, B. (2013). CluePedia Cytoscape plugin: pathway insights using integrated experimental and in silico data. Bioinformatics 29:661–663.

Bindea, G., Mlecnik, B., Hackl, H., Charoentong, P., Tosolini, M., Kirilovsky, A., Fridman, W.H., Pages, F., Trajanoski, Z., and Galon, J. (2009). ClueGO: a Cytoscape plug-in to decipher functionally grouped gene ontology and pathway annotation networks. Bioinformatics 25:1091–1093.

Birchler, J.A., Yao, H., and Chudalayandi, S. (2006). Unraveling the genetic basis of hybrid vigor. Proc. Natl. Acad. Sci. USA 103:12957–12958.

Birchler, J.A., Yao, H., Chudalayandi, S., Vaiman, D., and Veitia, R.A. (2010). Heterosis. Plant Cell 22:2105–2112.

Bruce, A.B. (1910). The Mendelian theory of heredity and the augmentation of vigor. Science 32:627–628.

Chambers, M.C., MacLean, B., Burke, R., Amode, D., Ruderman, D.L., Neumann, S., Gatto, L., Fischer, B., Pratt, B., Egertson, J., et al. (2012). A cross-platform toolkit for mass spectrometry and proteomics. Nat. Biotechnol. 30:918–920

Chehab, E.W., Raman, G., Walley, J.W., Perea, J.V., Banu, G., Theg, S., and Dehesh, K. (2006). Rice HYDROPEROXIDE LYASES with unique expression patterns generate distinct aldehyde signatures in Arabidopsis. Plant Physiol. 141:121–134.

Chen, K., Wallis, J.W., McLellan, M.D., Larson, D.E., Kalicki, J.M., Pohl, C.S., McGrath, S.D., Wendl, M.C., Zhang, Q., Locke, D.P., et al. (2009). BreakDancer: an algorithm for high-resolution mapping of genomic structural variation. Nat. Methods 6:677–681.

Chen, Y., Dan, Z., and Li, S. (2024). *GROWTH REGULATING FACTOR 7*-mediated arbutin metabolism enhances rice salt tolerance. Plant Cell 36:2834–2850.

Chong, J., Soufan, O., Li, C., Caraus, I., Li, S., Bourque, G., Wishart, D.S., and Xia, J. (2018). MetaboAnalyst 4.0: towards more transparent and integrative metabolomics analysis. Nucleic Acids Res. 46:W486–W494.

Crow, J.F. (1948). Alternative hypotheses of hybrid vigor. Genetics 33:477–487.

Dan, Z., Chen, Y., and Huang, W. (2024a). The metabolic pathways underlying dynamic heterosis for plant height are robust biomarkers in rice. bioRxiv:10.1101/2024.1111.1109.622823.

Dan, Z., Chen, Y., and Huang, W. (2024b). Structural variations contribute to genetic diversity and heterosis in rice. bioRxiv:10.1101/2024.1111.1103.621459.

Dan, Z., Chen, Y., Li, H., Zeng, Y., Xu, W., Zhao, W., He, R., and Huang, W. (2021). The metabolomic landscape of rice heterosis highlights pathway biomarkers for predicting complex phenotypes. Plant Physiol. 187:1011–1025.

Dan, Z., Chen, Y., Xu, Y., Huang, J.R., Huang, J.S., Hu, J., Yao, G., Zhu, Y., and Huang, W. (2019). A metabolome-based core hybridisation strategy for the prediction of rice grain weight across environments. Plant Biotechnol. J. 17:906–913.

Dan, Z., Chen, Y., Zhao, W., Wang, Q., and Huang, W. (2020). Metabolome-based prediction of yield heterosis contributes to the breeding of elite rice. Life Sci. Alliance 3:e201900551.

Dan, Z., Hu, J., Zhou, W., Yao, G., Zhu, R., Huang, W., and Zhu, Y. (2015). Hierarchical additive effects on heterosis in rice (*Oryza sativa* L.). Front. Plant Sci. 6:738.

Dan, Z., Hu, J., Zhou, W., Yao, G., Zhu, R., Zhu, Y., and Huang, W. (2016). Metabolic prediction of important agronomic traits in hybrid rice (*Oryza sativa* L.). Sci. Rep. 6:21732.

Dan, Z., Liu, P., Huang, W., Zhou, W., Yao, G., Hu, J., Zhu, R., Lu, B., and Zhu, Y. (2014). Balance between a higher degree of heterosis and increased reproductive isolation: a strategic design for breeding inter-subspecific hybrid rice. PLoS ONE 9:e93122.

Davenport, C.B. (1908). Degeneration, albinism and inbreeding. Science 28:454–455.

de Abreu, E.L.F., Westhues, M., Cuadros-Inostroza, Á., Willmitzer, L., Melchinger, A.E., and Nikoloski, Z. (2017). Metabolic robustness in young roots underpins a predictive model of maize hybrid performance in the field. Plant J. 90:319–329.

Domingo-Almenara, X., Montenegro-Burke, J.R., Ivanisevic, J., Thomas, A., Sidibe, J., Teav, T., Guijas, C., Aisporna, A.E., Rinehart, D., Hoang, L., et al. (2018). XCMS-MRM and METLIN-MRM: a cloud library and public resource for targeted analysis of small molecules. Nat. Methods 15:681–684.

East, E.M. (1936). Heterosis. Genetics 21:375–397.

Eriksson, L., Hermens, J., L. M., Johansson, E., Verhaar, H., J. M., and Wold, S. (1995). Multivariate analysis of aquatic toxicity data with PLS. Aquat. Sci. 57:217–241.

Gao, Z., Wang, Y., Chen, G., Zhang, A., Yang, S., Shang, L., Wang, D., Ruan, B., Liu, C., Jiang, H., et al. (2019). The indica nitrate reductase gene OsNR2 allele enhances rice yield potential and nitrogen use efficiency. Nat. Commun. 10:5207.

Gowda, H., Ivanisevic, J., Johnson, C.H., Kurczy, M.E., Benton, H.P., Rinehart, D., Nguyen, T., Ray, J., Kuehl, J., Arevalo, B., et al. (2014). Interactive XCMS Online: simplifying advanced metabolomic data processing and subsequent statistical analyses. Anal. Chem. 86:6931–6939.

Hochholdinger, F., and Baldauf, J.A. (2018). Heterosis in plants. Curr. Biol. 28:R1089–R1092.

Huang, X., Yang, S., Gong, J., Zhao, Q., Feng, Q., Zhan, Q., Zhao, Y., Li, W., Cheng, B., Xia, J., et al. (2016). Genomic architecture of heterosis for yield traits in rice. Nature 537:629–633.

Jia, H., Shen, X., Guan, Y., Xu, M., Tu, J., Mo, M., Xie, L., Yuan, J., Zhang, Z., Cai, S., et al. (2018). Predicting the pathological response to neoadjuvant chemoradiation using untargeted metabolomics in locally advanced rectal cancer. Radiother Oncol 128:548–556.

Jones, D.F. (1917). Dominance of linked factors as a means of accounting for heterosis. Proc. Natl. Acad. Sci. USA 3:310–312.

Kakoulidou, I., Piecyk, R.S., Meyer, R.C., Kuhlmann, M., Gutjahr, C., Altmann, T., and Johannes, F. (2024). Mapping parental DMRs predictive of local and distal methylome remodeling in epigenetic F1 hybrids. Life Sci. Alliance 7:e202402599.

Kim, D., Langmead, B., and Salzberg, S.L. (2015). HISAT: a fast spliced aligner with low memory requirements. Nat. Methods 12:357–360.

Krieger, U., Lippman, Z.B., and Zamir, D. (2010). The flowering gene *SINGLE FLOWER TRUSS* drives heterosis for yield in tomato. Nat. Genet. 42:459–463.

Krzywinski, M., Schein, J., Birol, I., Connors, J., Gascoyne, R., Horsman, D., Jones, S.J., and Marra, M.A. (2009). Circos: an information aesthetic for comparative genomics. Genome Res. 19:1639–1645.

Li, D., Huang, Z., Song, S., Xin, Y., Mao, D., Lv, Q., Zhou, M., Tian, D., Tang, M., Wu, Q., et al. (2016). Integrated analysis of phenome, genome, and transcriptome of hybrid rice uncovered multiple heterosis-related loci for yield increase. Proc Natl Acad Sci U S A 113:E6026–E6035.

Li, H., and Durbin, R. (2009). Fast and accurate short read alignment with Burrows-Wheeler transform. Bioinformatics 25:1754–1760.

Li, H., Handsaker, B., Wysoker, A., Fennell, T., Ruan, J., Homer, N., Marth, G., Abecasis, G., Durbin, R., and Genome Project Data Processing, S. (2009). The Sequence Alignment/Map format and SAMtools. Bioinformatics 25:2078-2079.

Li, S., Park, Y., Duraisingham, S., Strobel, F.H., Khan, N., Soltow, Q.A., Jones, D.P., and Pulendran, B. (2013). Predicting network activity from high throughput metabolomics. PLoS Comput. Biol. 9:e1003123.

Li, X., Li, X., Fridman, E., Tesso, T.T., and Yu, J. (2015). Dissecting repulsion linkage in the dwarfing gene *Dw3* region for sorghum plant height provides insights into heterosis. Proc. Natl. Acad. Sci. USA 112:11823–11828.

Li, Z., Pinson, S., Park, W., Paterson, A., and Stansel, J. (1997). Epistasis for three grain yield components in rice (*Oryza sativa* L.). Genetics 145:453–465.

Liao, Y., Smyth, G.K., and Shi, W. (2014). featureCounts: an efficient general purpose program for assigning sequence reads to genomic features. Bioinformatics 30:923–930.

Lisec, J., Romisch-Margl, L., Nikoloski, Z., Piepho, H.P., Giavalisco, P., Selbig, J., Gierl, A., and Willmitzer, L. (2011). Corn hybrids display lower metabolite variability and complex metabolite inheritance patterns. Plant J. 68:326–336.

Liu, N., Du, Y., Warburton, M.L., Xiao, Y., and Yan, J. (2021a). Phenotypic plasticity contributes to maize adaptation and heterosis. Mol Biol Evol 38:1262–1275.

Liu, W., He, G., and Deng, X.W. (2021b). Biological pathway expression complementation contributes to biomass heterosis in Arabidopsis. Proc. Natl. Acad. Sci. USA 118:e2023278118.

Lu, B., Cai, X., and Jin, X. (2009). Efficient *indica* and *japonica* rice identification based on the InDel molecular method:its implication in rice breeding and evolutionary research. Progress in Nature Science 19:1241-1252.

Metsalu, T., and Vilo, J. (2015). ClustVis: a web tool for visualizing clustering of multivariate data using Principal Component Analysis and heatmap. Nucleic Acids Res. 43:W566–570.

Pang, Z., Zhou, G., Ewald, J., Chang, L., Hacariz, O., Basu, N., and Xia, J. (2022). Using MetaboAnalyst 5.0 for LC-HRMS spectra processing, multi-omics integration and covariate adjustment of global metabolomics data. Nat. Protoc.

Powers, L. (1944). An expansion of Jones’s theory for the explanation of heterosis. Am. Nat. 78:275–280.

Price, N.D., Magis, A.T., Earls, J.C., Glusman, G., Levy, R., Lausted, C., McDonald, D.T., Kusebauch, U., Moss, C.L., Zhou, Y., et al. (2017). A wellness study of 108 individuals using personal, dense, dynamic data clouds. Nat. Biotechnol. 35:747–756.

Saito, R., Smoot, M.E., Ono, K., Ruscheinski, J., Wang, P.L., Lotia, S., Pico, A.R., Bader, G.D., and Ideker, T. (2012). A travel guide to Cytoscape plugins. Nat. Methods 9:1069–1076.

Schnable, P.S., and Springer, N.M. (2013). Progress toward understanding heterosis in crop plants. Annu. Rev. Plant Biol. 64:71–88.

Schrag, T., Westhues, M., Schipprack, W., Seifert, F., Thiemann, A., Scholten, S., and Melchinger, A.E. (2018). Beyond genomic prediction: combining different types of *omics* data can improve prediction of hybrid performance in maize. Genetics 208:1373–1385.

Shen, X., Wang, R., Xiong, X., Yin, Y., Cai, Y., Ma, Z., Liu, N., and Zhu, Z.-J. (2019). Metabolic reaction network-based recursive metabolite annotation for untargeted metabolomics. Nat. Commun. 10:1516.

Shull, G.H. (1948). What is "heterosis"? Genetics 33:439–446.

Spitzer, M., Wildenhain, J., Rappsilber, J., and Tyers, M. (2014). BoxPlotR: a web tool for generation of box plots. Nat. Methods 11:121–122.

Stecher, G., Tamura, K., and Kumar, S. (2020). Molecular Evolutionary Genetics Analysis (MEGA) for macOS. Mol Biol Evol 37:1237–1239.

Sun, Z., Peng, J., Lv, Q., Ding, J., Chen, S., Duan, M., He, Q., Wu, J., Tian, Y., Yu, D., et al. (2023). Dissecting the genetic basis of heterosis in elite super-hybrid rice. Plant Physiol. 192:307–325.

Tu, J., Yin, Y., Xu, M., Wang, R., and Zhu, Z.J. (2017). Absolute quantitative lipidomics reveals lipidome-wide alterations in aging brain. Metabolomics 14:5.

Wang, B., Hou, M., Shi, J., Ku, L., Song, W., Li, C., Ning, Q., Li, X., Li, C., Zhao, B., et al. (2023). De novo genome assembly and analyses of 12 founder inbred lines provide insights into maize heterosis. Nat. Genet. 55:312–323.

Wang, K., Li, M., and Hakonarson, H. (2010). ANNOVAR: functional annotation of genetic variants from high-throughput sequencing data. Nucleic Acids Res. 38:e164.

Wang, X., Zhou, T., Li, G., Yao, W., Hu, W., Wei, X., Che, J., Yang, H., Shao, L., Hua, J., et al. (2022). A Ghd7-centered regulatory network provides a mechanistic approximation to optimal heterosis in an elite rice hybrid. Plant J. 112:68–83.

Wang, Z., Xue, Z., and Wang, T. (2014). Differential analysis of proteomes and metabolomes reveals additively balanced networking for metabolism in maize heterosis. J. Proteome Res. 13:3987–4001.

Waterhouse, A.M., Procter, J.B., Martin, D.M., Clamp, M., and Barton, G.J. (2009). Jalview Version 2-a multiple sequence alignment editor and analysis workbench. Bioinformatics 25:1189-1191.

Williams, W. (1959). Heterosis and the genetics of complex characters. Nature 184:527–530.

Wold, S., Sjöström, M., and Eriksson, L. (2001). PLS-regression: a basic tool of chemometrics. Chemometrics Intellig. Lab. Syst. 58:109–130.

Xiao, Y., Jiang, S., Cheng, Q., Wang, X., Yan, J., Zhang, R., Qiao, F., Ma, C., Luo, J., Li, W., et al. (2021). The genetic mechanism of heterosis utilization in maize improvement. Genome Biol. 22:148.

Xu, Y., Zhao, Y., Wang, X., Ma, Y., Li, P., Yang, Z., Zhang, X., Xu, C., and Xu, S. (2021). Incorporation of parental phenotypic data into multi-omic models improves prediction of yield-related traits in hybrid rice. Plant Biotechnol. J. 19:261–272.

Yao, G., and Huang, W. (2013). Identifying *indica*-*japonica* characteristic of rice breeding materials with *indica*-*japonica* differentiation InDel markers. Hybrid Rice 28:53-57.

Yu, S.B., Li, J.X., Xu, C.G., Tan, Y.F., Gao, Y.J., Li, X.H., Zhang, Q., and Saghai Maroof, M.A. (1997). Importance of epistasis as the genetic basis of heterosis in an elite rice hybrid. Proc. Natl. Acad. Sci. USA 94:9226–9231.

Zhang, M., Wang, Y., Hu, Y., Wang, H., Liu, Y., Zhao, B., Zhang, J., Fang, R., and Yan, Y. (2023). Heterosis in root microbiota inhibits growth of soil-borne fungal pathogens in hybrid rice. J. Integr. Plant Biol. 65:1059–1076.

Zhou, G., Ewald, J., and Xia, J. (2021a). OmicsAnalyst: a comprehensive web-based platform for visual analytics of multi-omics data. Nucleic Acids Res.

Zhou, G., Soufan, O., Ewald, J., Hancock, R.E.W., Basu, N., and Xia, J. (2019a). NetworkAnalyst 3.0: a visual analytics platform for comprehensive gene expression profiling and meta-analysis. Nucleic Acids Res. 47:W234-W241.

Zhou, P., Hirsch, C.N., Briggs, S.P., and Springer, N.M. (2019b). Dynamic patterns of gene expression additivity and regulatory variation throughout maize development. Mol. Plant 12:410–425.

Zhou, X., Nong, C., Wu, B., Zhou, T., Zhang, B., Liu, X., Gao, G., Mi, J., Zhang, Q., Liu, H., et al. (2021b). Combinations of *Ghd7*, Ghd8, and *Hd1* determine strong heterosis of commercial rice hybrids in diverse ecological regions. J. Exp. Bot. 72:6963-6976.

