## Supplemental figures for "Additive and partially dominant effects from genomic variation contribute to rice heterosis"

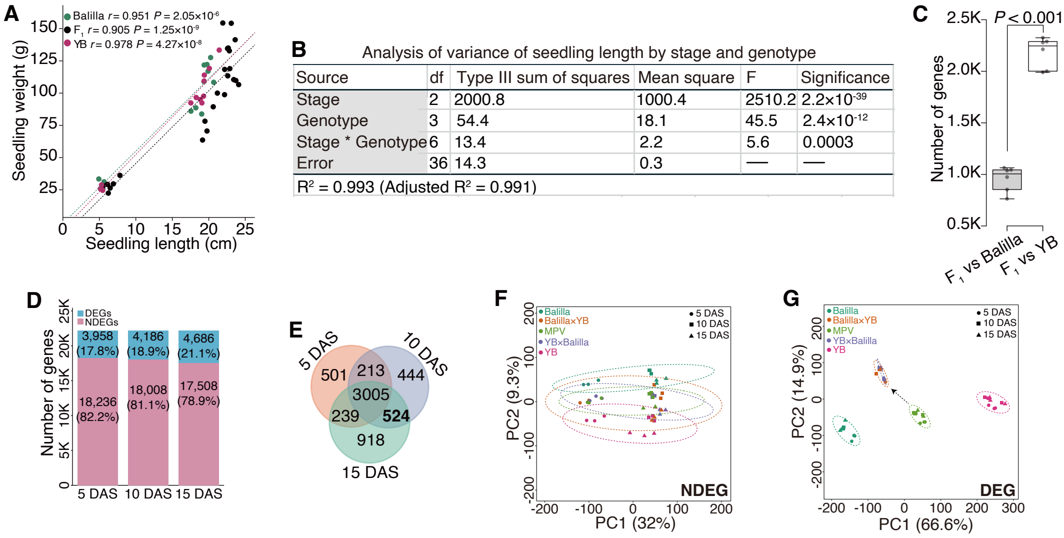


**Supplemental Figure 1. Transcriptomic analysis of seedling samples from two rice inbred lines (Balilla and Yuetai B) and their reciprocal $F_1$ hybrids across three developmental stages.**

**(A)** Correlations between seedling length and seedling weight in parents and $F_1$ hybrids across stages. *P* values indicate Pearson correlation.

**(B)** Analysis of variance (ANOVA) of seedling length by genotype, stage, and their interaction.

**(C)** Number of differentially expressed genes (DEGs) between $F_1$ hybrids and parents. *P* value is for independent samples *t* test (weighted).

**(D)** Number of DEGs and non-differentially expressed genes (NDEGs) at 5, 10, and 15 days after sowing (DAS).

**(E)** Venn diagram of DEGs across stages.

**(F-G)** PCA score plots of Balilla, Yuetai B (YB), $F_1$ hybrids, and mid-parent values (MVP) based on NDEGs (**F**; 16,350 genes) and DEGs (**G**; 5,844 genes).

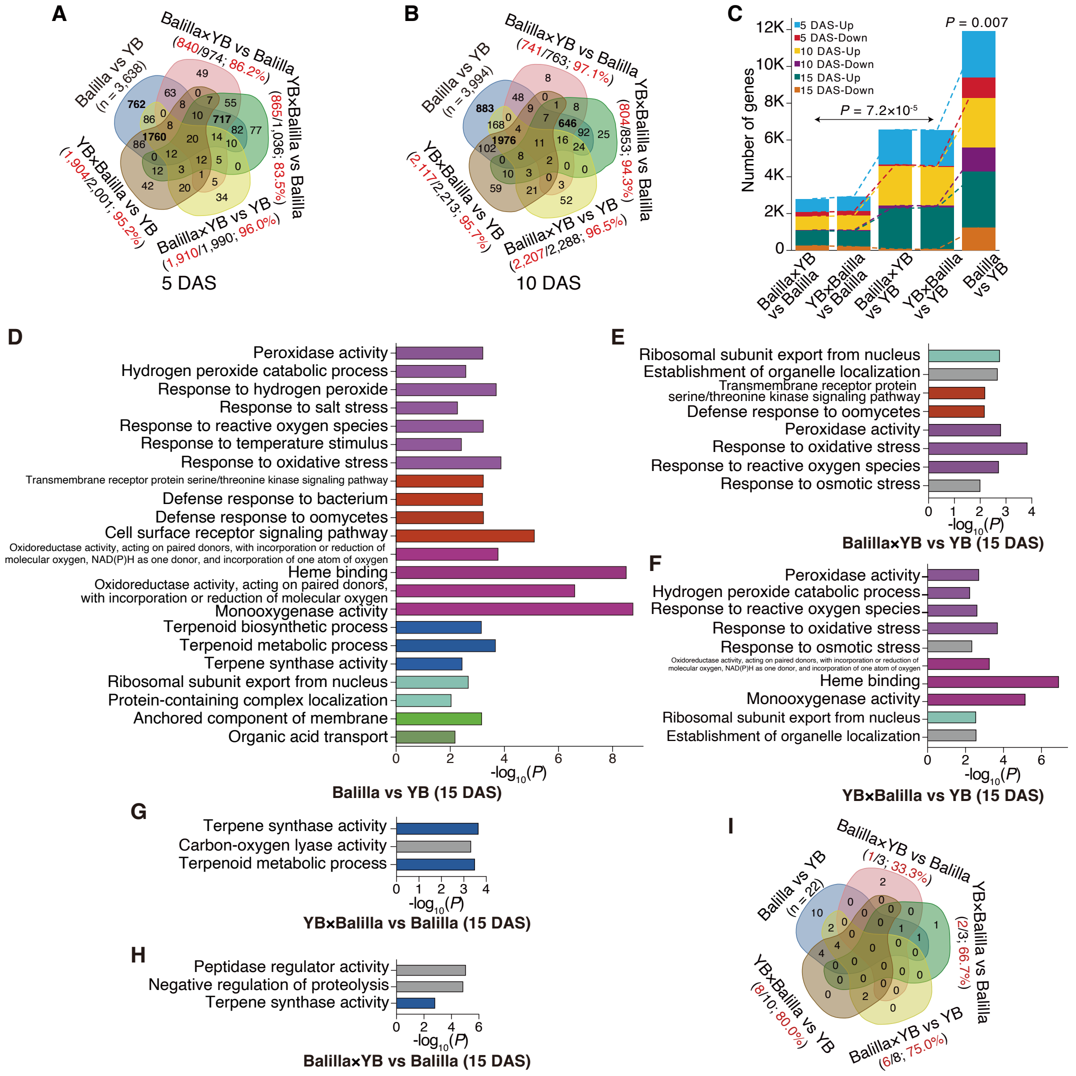


**Supplemental Figure 2. Transcriptomic differences between two inbred lines and between $F_1$ hybrids and corresponding parents.**

**(A-B)** Venn diagrams of differentially expressed genes (DEGs) between inbred lines (Balilla vs YB) and between $F_1$ hybrids and corresponding parents at 5 and 10 days after sowing (DAS). Red number/percentages represent DEGs derived from parental differences.

**(C)** Number of up-/downregulated genes between the two inbred lines and between $F_1$ hybrids and corresponding parents. *P* values indicate paired samples *t* tests.

**(D-H)** Significantly enriched terms between the two inbred lines and between $F_1$ hybrids and their parents at 15 DAS.

**(I)** Venn diagram of differentially enriched terms between the two inbred lines and between $F_1$ hybrids and their parents at 15 DAS. Red number represent differential terms derived from parental differences.

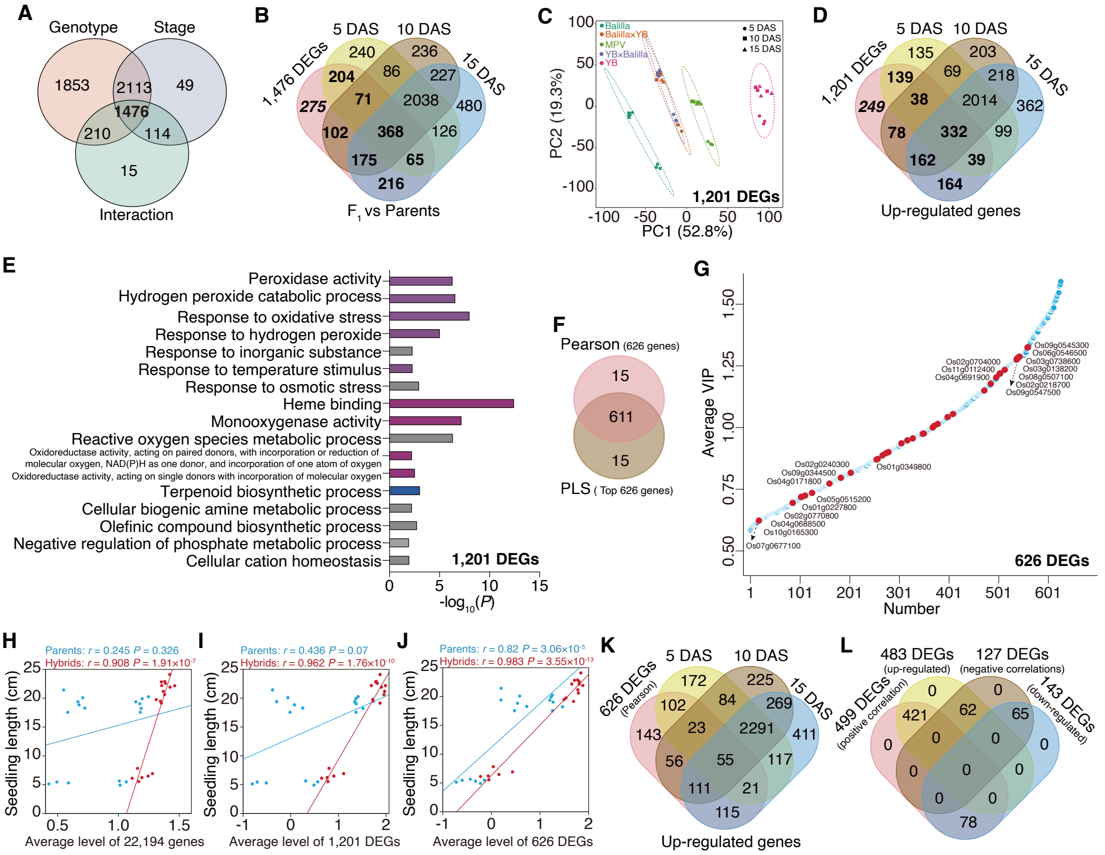


**Supplemental Figure 3. Identification of heterosis-associated genes for seedling length.**

**(A)** Venn diagram of 5,844 differentially expressed genes (DEGs) responsive to genotype, stage, and their interaction.

**(B)** Subset of 1,476 DEGs. These genes included DEGs between inbred lines, $F_1$ hybrids and parents, and reciprocal $F_1$ hybrids. The 275 DEGs specific to inbred lines and reciprocal $F_1$ hybrids were excluded.

**(C)** PCA score plot of Balilla, Yuetai B (YB), $F_1$ hybrids, and mid-parent values (MPVs) based on 1,201 DEGs.

**(D)** Venn diagram of 1,201 DEGs upregulated in $F_1$ hybrids relative to their parents.

**(E)** Significantly enriched GO terms for 1,201 DEGs.

**(F)** Overlap of 626 genes significantly correlated with seedling length (Pearson, $P < 0.05$) and 626 top-contributing genes from partial least squares (PLS) analysis (average values of variable importance in projections).

**(G)** Contributing order of 626 genes in PLS analysis. The 37 heterosis-associated genes (red) are labeled; top/bottom ten genes include IDs.

**(H-J)** Correlations between gene expression and seedling length in parents and $F_1$ hybrids. $P$ values indicate Pearson correlation.

**(K)** Overlap of 626 DEGs upregulated in $F_1$ hybrids relative to their parents.

**(L)** Upregulated genes predominantly had positive correlations with seedling length.

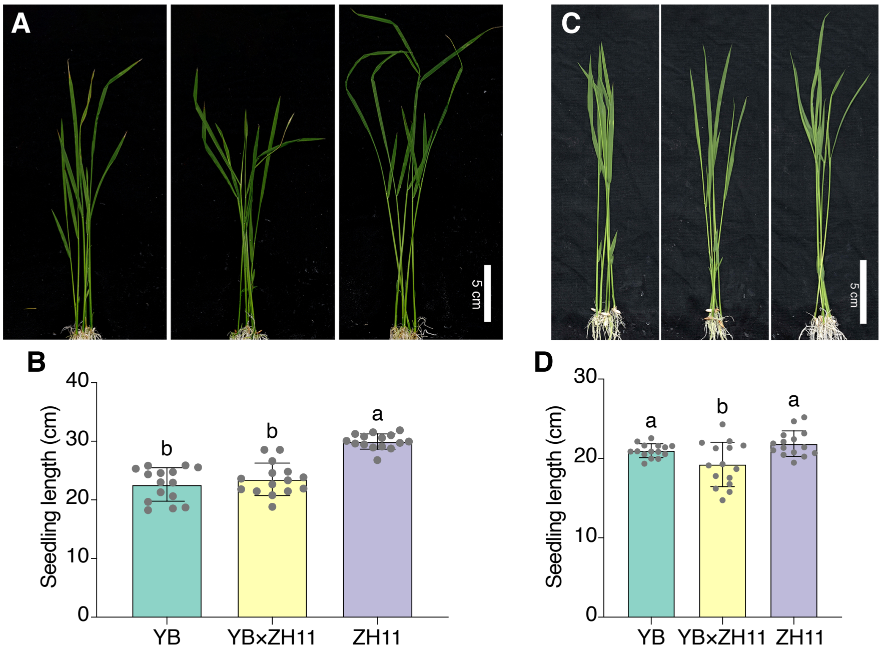


**Supplemental Figure 4. Seedlings and seedling length of YB, Zhonghua11 (ZH11), and their $F_1$ hybrids.**

**(A-B)** Seedlings grown at 28 °C under 24-hour light and their seedling length measured at 15 days after sowing.

**(C-D)** Seedlings grown at 28 °C under 16-hour light/8-hour dark and their seedling length measured at 10 days after sowing. Scare bar, 5 cm. n = 15 seedlings. Analysis of variance was implemented to compare differences.

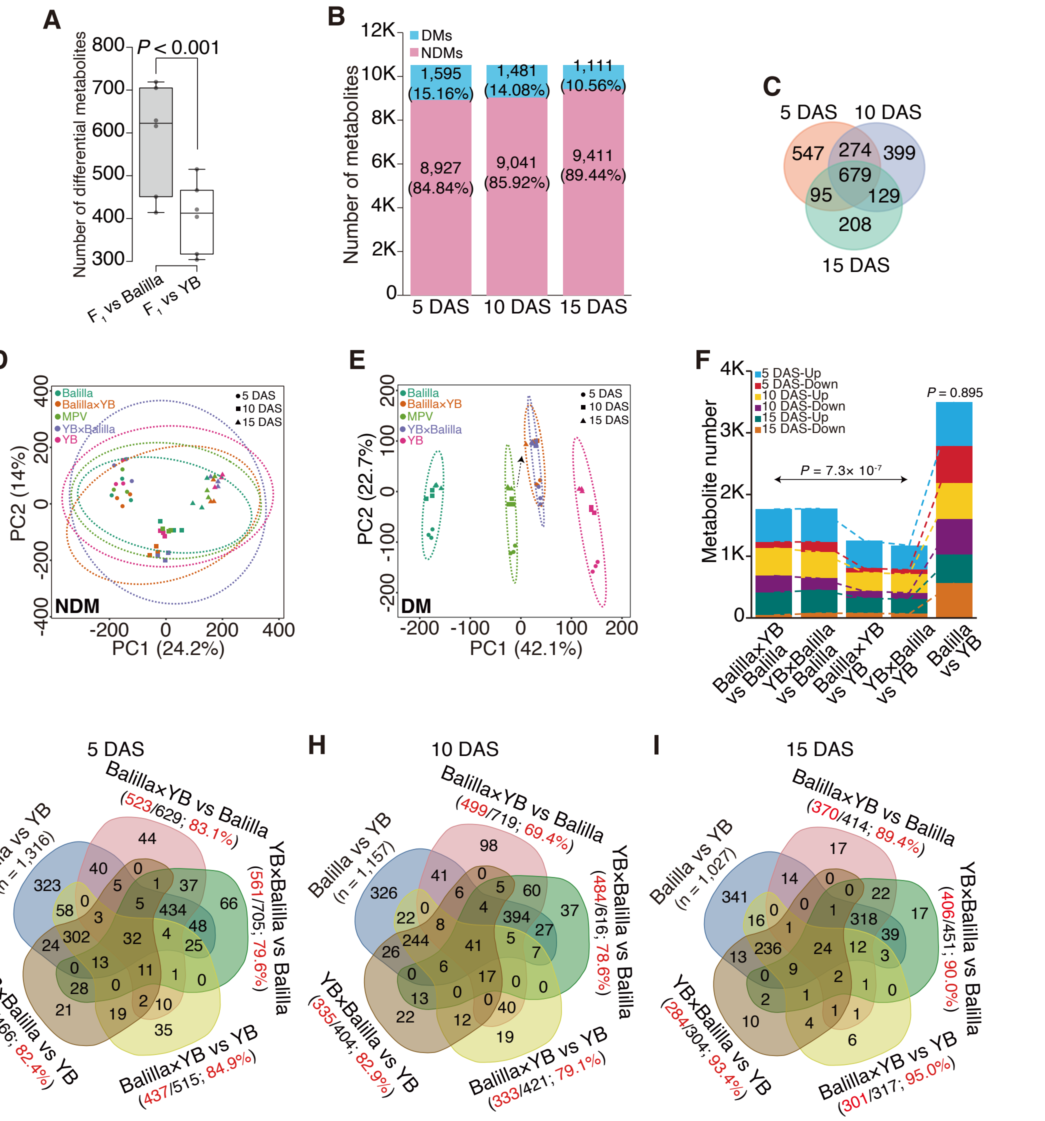

**Supplemental Figure 5. Differential metabolite analysis of two inbred lines and $F_1$ hybrids.**

**(A)** Number of differential metabolites (DMs) between $F_1$ hybrids and parents. *P* value indicates independent samples *t* test. The number are weighted before analysis.

**(B)** Differential and non-differential metabolites (NDMs) at 5, 10, and 15 days after sowing (DAS).

**(C)** Venn diagram of DMs across stages.

**(D-E)** PCA score plots of Balilla, Yuetai B (YB), $F_1$ hybrids, and mid-parent values (MPVs) based on NDMs (**D**; 8,191 metabolites) and DMs (**E**; 2,331 metabolites).

**(F)** Up-/downregulated metabolites between two inbred lines and between $F_1$ hybrids and parents. *P* values indicate paired samples *t* tests.

**(G-I)** Venn diagrams of DMs across stages. Red number/percentages indicate DMs between $F_1$ hybrids and parents derived from parental DMs.

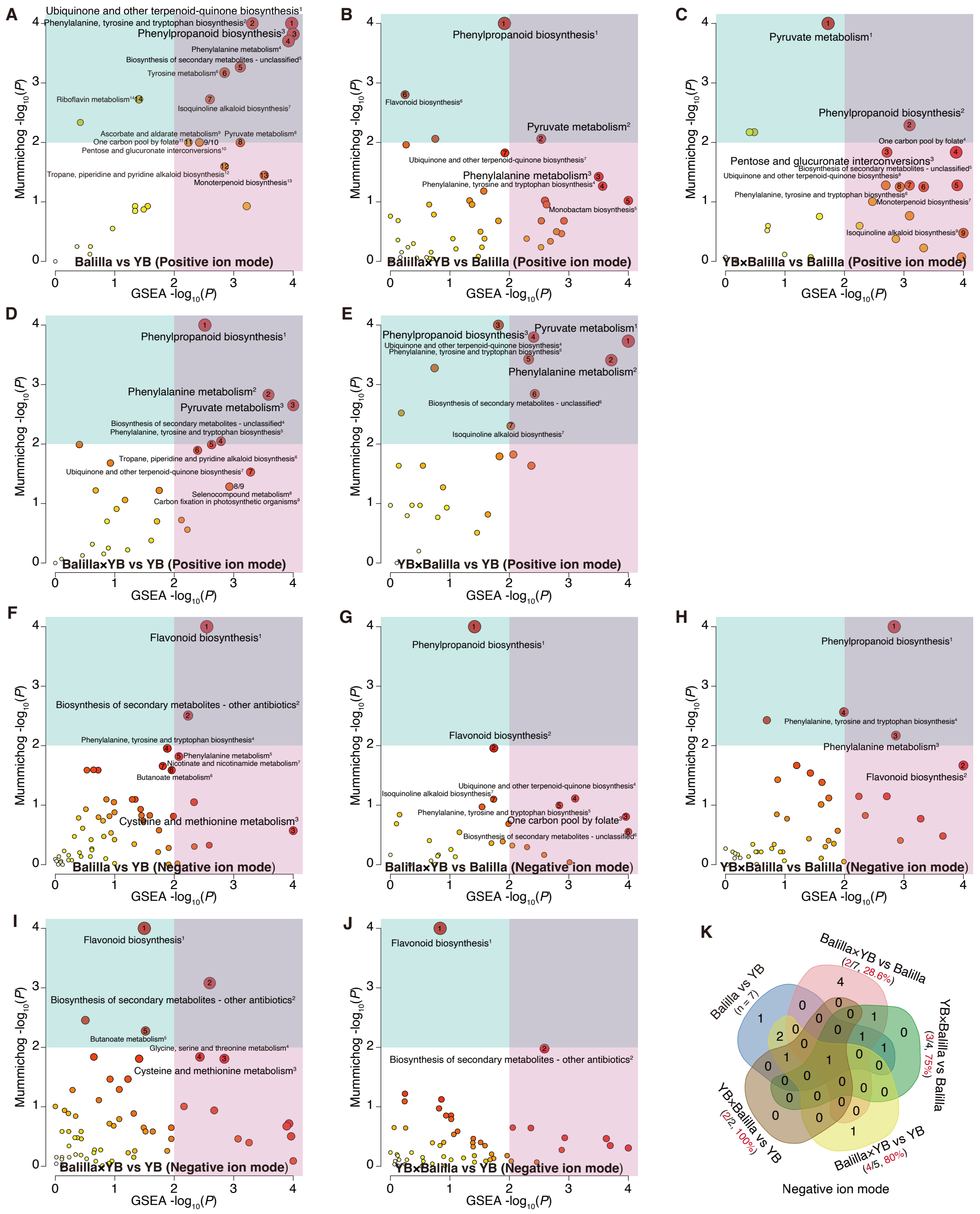


**Supplemental Figure 6. Differential pathways of two inbred lines and $F_1$ hybrids.** Enriched pathways for differential metabolites in positive (**A-E**) and negative (**F-J**) ion modes are displayed. Differential metabolites between Balilla and YB (**A** and **F**), between $F_1$ hybrid (Balilla×YB) and Balilla (**B** and **G**), between $F_1$ hybrid (YB×Balilla) and Balilla (**C** and **H**), between $F_1$ hybrid (Balilla×YB) and YB (**D** and **I**), and between $F_1$ hybrid (YB×Balilla) and YB (**E** and **J**), were used for enrichment analysis.

**(K)** Venn diagram of significantly enriched pathways (negative ion mode). Red number/percentages indicate differential pathways derived from parental differences.

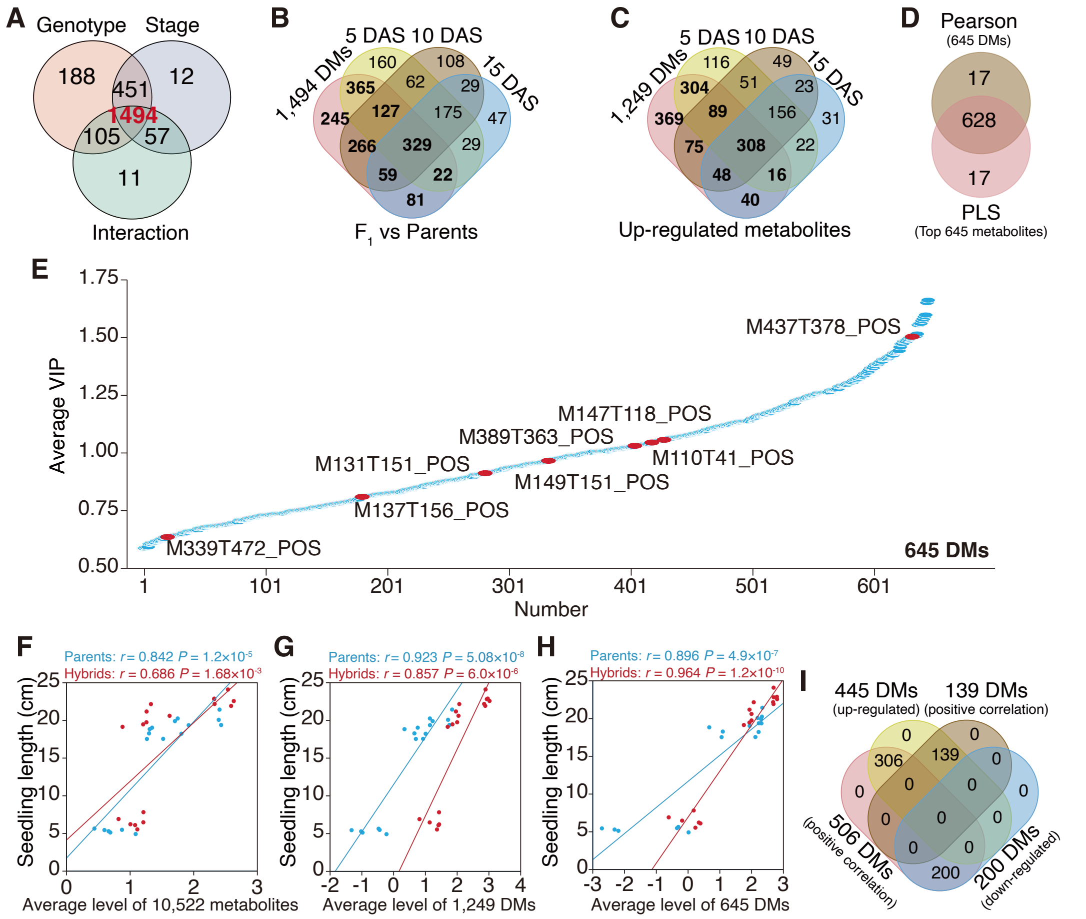


**Supplemental Figure 7. Identification of heterosis-associated metabolites for seedling length.**

**(A)** Venn diagram of 2,331 differential metabolites (DMs) responsive to genotype, stage, and their interaction.

**(B)** Subset of 1,494 DMs. These metabolites include DMs between two inbred lines, between $F_1$ hybrids and parents, and between reciprocal $F_1$ hybrids. The 245 DMs specific to parents and reciprocal $F_1$ hybrids were excluded.

**(C)** Venn diagram of 1,249 DMs upregulated in $F_1$ hybrids relative to their parents.

**(D)** Overlap of 645 metabolites significantly correlated with seedling length (Pearson, $P < 0.05$) and 645 top contributing metabolites from partial least squares (PLS) analysis (average values of variable importance in projections).

**(E)** Contributing order of 645 metabolites in PLS analysis. Eight metabolites from three significantly enriched pathways are marked in red.

**(F-H)** Correlations between metabolite levels and seedling length in parents and $F_1$ hybrids. *P* values indicate Pearson correlation.

**(I)** Upregulated metabolites predominantly had positive correlations with seedling length.

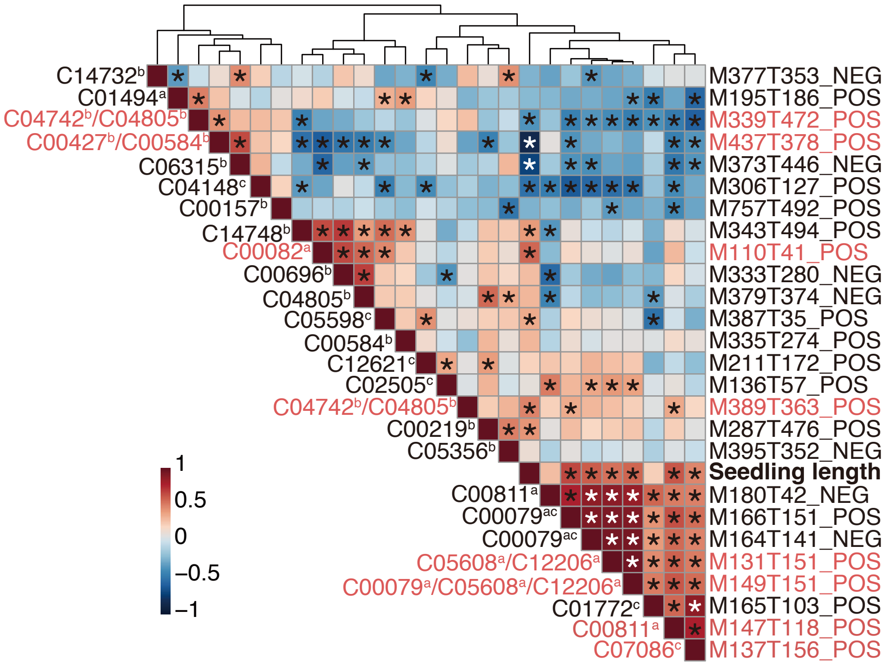


**Supplemental Figure 8. Correlation heatmap of seedling length and heterosis-associated metabolites from three significantly enriched pathways.** Eight peaks annotated by *mummichog* algorithm are labeled in red, and chemically annotated peaks are labeled in black. Metabolites from the three pathways were marked with letters (a, b, and c) according to Figure 2E. Asterisks indicate significant Pearson correlations ($P < 0.05$)

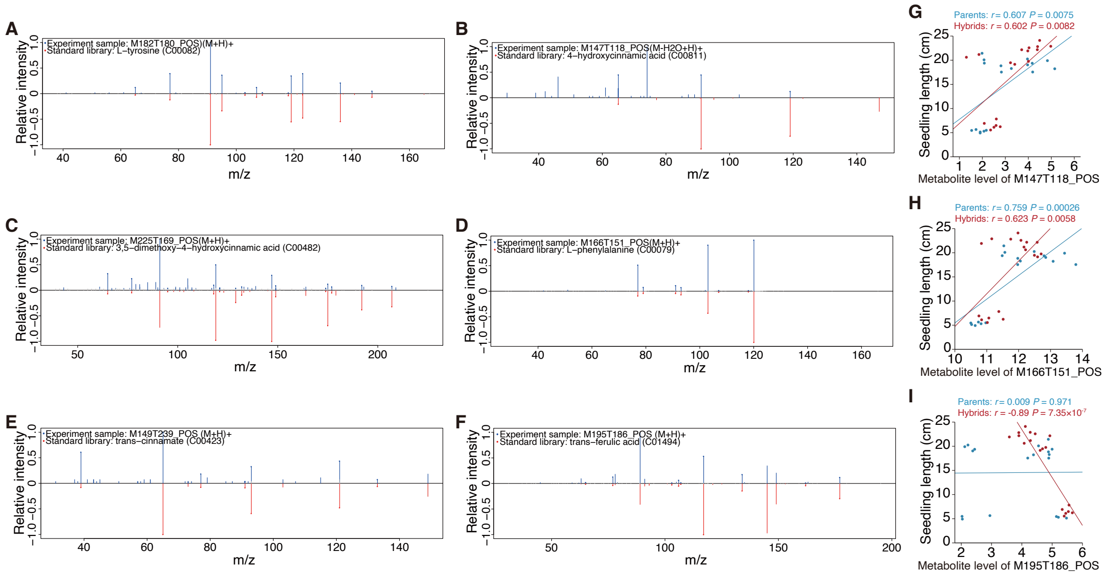


**Supplemental Figure 9. MS/MS spectra of six metabolites and their correlations with seedling length.**

**(A-F)** MS/MS spectra (experimental samples above, standards below horizontal line) of six metabolites.

**(G-I)** Correlations between three chemically identified metabolites and seedling length. *P* values indicate Pearson correlation.

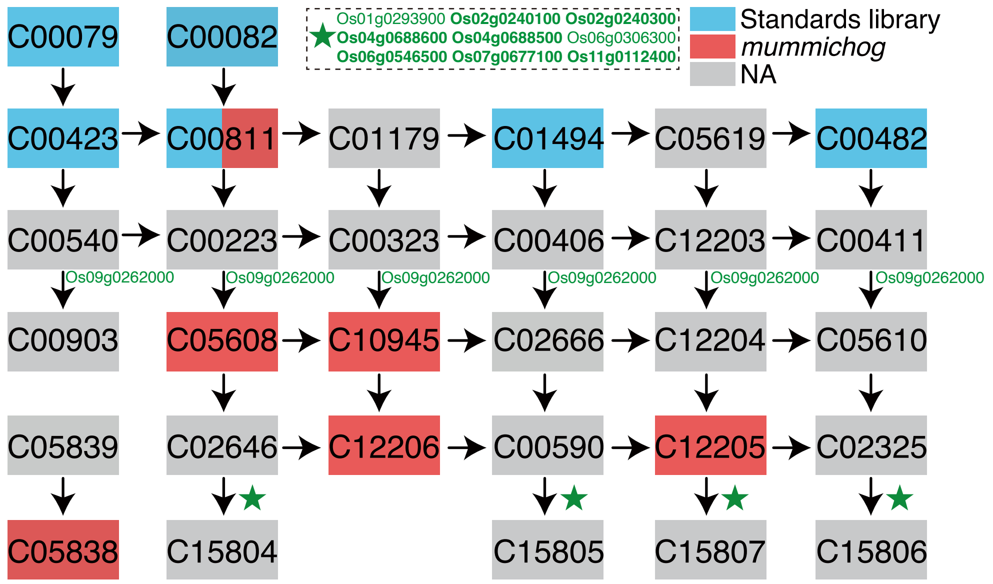


**Supplemental Figure 10. Heterosis-associated genes and metabolites in phenylpropanoid biosynthesis.** Red: *mummichog* algorithm-annotated metabolites; blue: chemical standards-annotated metabolites.

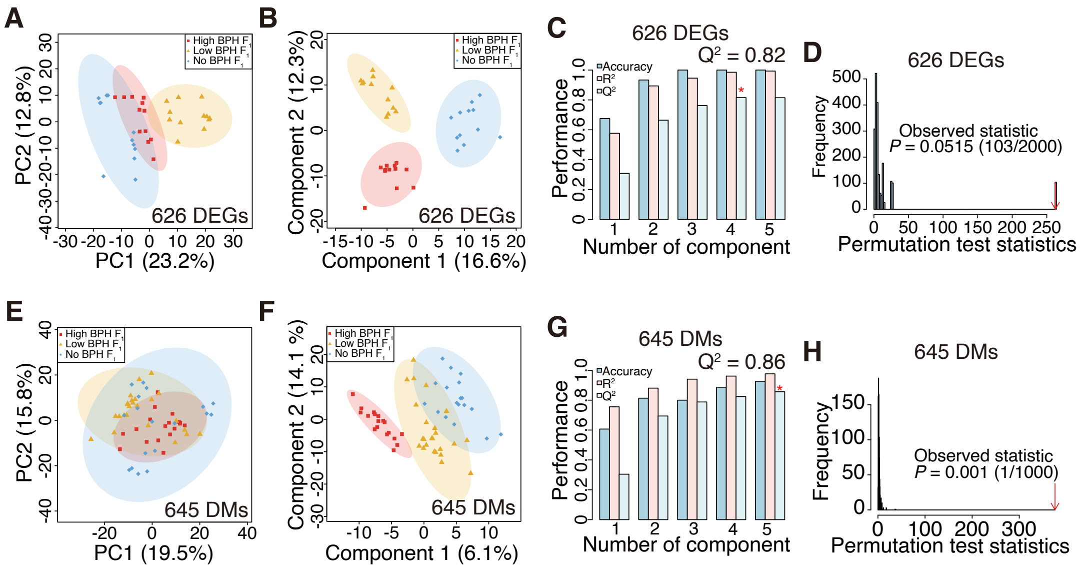


**Supplemental Figure 11. Validation of 626 heterosis-associated genes and 645 heterosis-associated metabolites for seedling length.**

**(A-D)** Analyses of $F_1$ hybrids exhibiting high-, low-, or no better-parent heterosis (BPH) for seedling length using 626 heterosis-associated genes. Shown are PCA score plots (**A**), PLS-DA score plots (**B**), PLS-DA performance (**C**), and PLS-DA permutation test results (**D**).

**(E-H)** Analyses of $F_1$ hybrids exhibiting high-, low-, or no BPH for seedling length using 645 heterosis-associated metabolites. Shown are PCA score plots (**E**), PLS-DA score plots (**F**), PLS-DA performance (**G**), and PLS-DA permutation test results (**H**). *P* values in **D** and **H** are for permutation test.

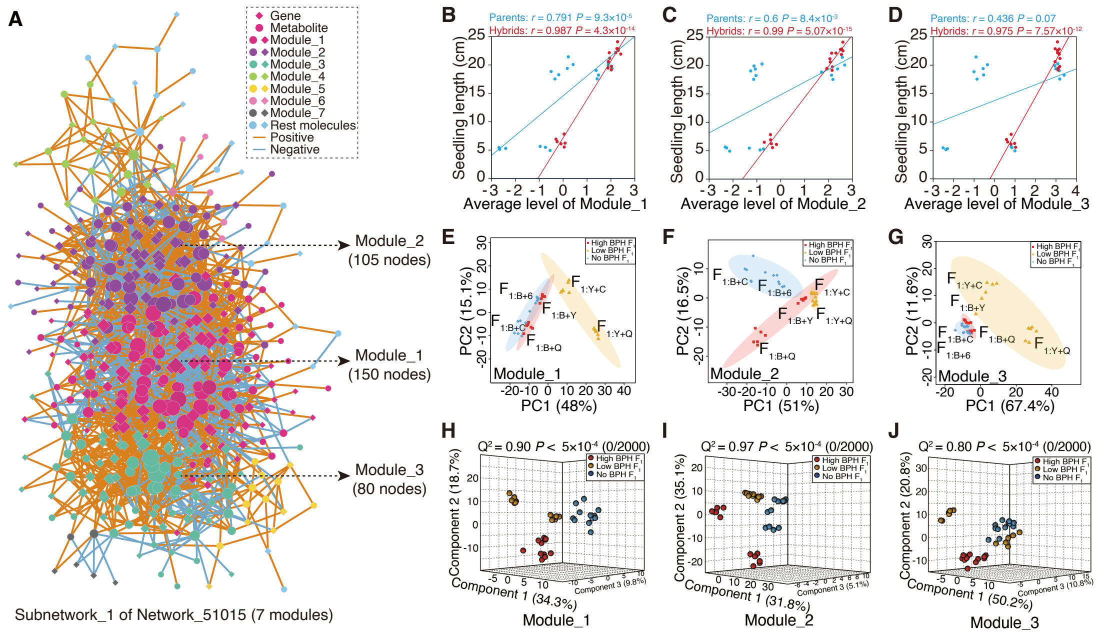


**Supplemental Figure 12. Validation of Network_51015 to heterosis for seedling length.**

**(A)** Subnetwork_1 of Network_51015 integrating 626 heterosis-associated genes and 645 heterosis-associated metabolites (2,138 edges). Node size reflects node degree.

**(B-D)** Pearson correlations between molecular levels of the top three modules and seedling length in parents and hybrids.

**(E-G)** PCA score plots of $F_1$ hybrids with high-, low-, or no better-parent heterosis (BPH) based on molecular levels of the top three modules.

**(H-J)** PLS-DA score plots of $F_1$ hybrids with high-, low-, or no BPH based on the top three modules. *P* values are for permutation test.

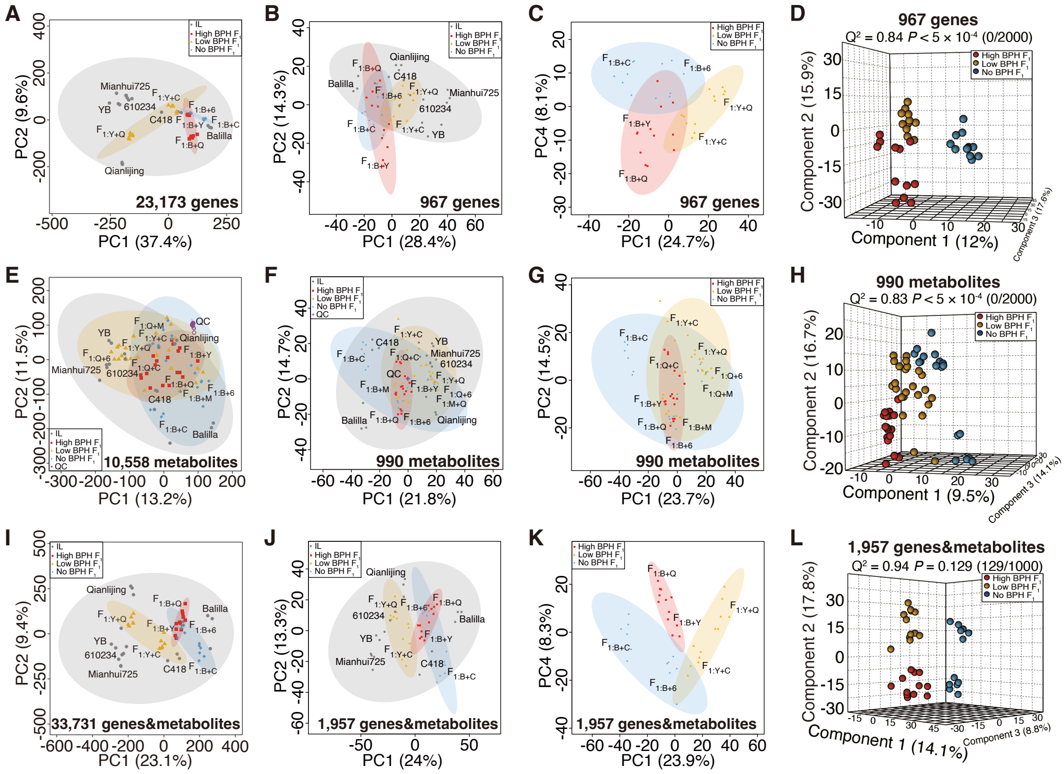

**Supplemental Figure 13. Score plots of parental inbred lines (ILs), $F_1$ hybrids, and quality controls (QCs) based on transcriptomic and metabolomic data.**

**(A-B)** PCA score plots of six ILs and six pairs of reciprocal $F_1$ hybrids on PC1 and PC2 using global gene expression levels (**A**; 23,173 genes) or 967 differentially expressed genes (DEGs; **B**) from the ILs.

**(C-D)** PCA (**C**) and PLS-DA (**D**) score plots of six pairs of reciprocal $F_1$ hybrids using 967 DEGs.

**(E-F)** PCA score plots of six ILs, ten pairs of reciprocal $F_1$ hybrids, and quality controls on PC1 and PC2 using all metabolites (**E**; 10,558 metabolites) or 990 differential metabolites (DMs; **F**) from the ILs.

**(G-H)** PCA (**G**) and PLS-DA (**H**) score plots of ten pairs of reciprocal $F_1$ hybrids using 990 DMs.

**(I-J)** PCA score plots of six ILs and six pairs of reciprocal $F_1$ hybrids on PC1 and PC2 using integrated molecular levels of all genes and metabolites (**I**; 23,173 genes + 10,558 metabolites) or integrated DEGs and DMs (**J**; 967 DEGs + 990 DMs).

**(K-L)** PCA (**K**) and PLS-DA (**L**) score plots of six pairs of reciprocal $F_1$ hybrids using integrated molecular levels of 967 DEGs and 990 DMs. *P* values in **D**, **H**, and **L** are for permutation test.

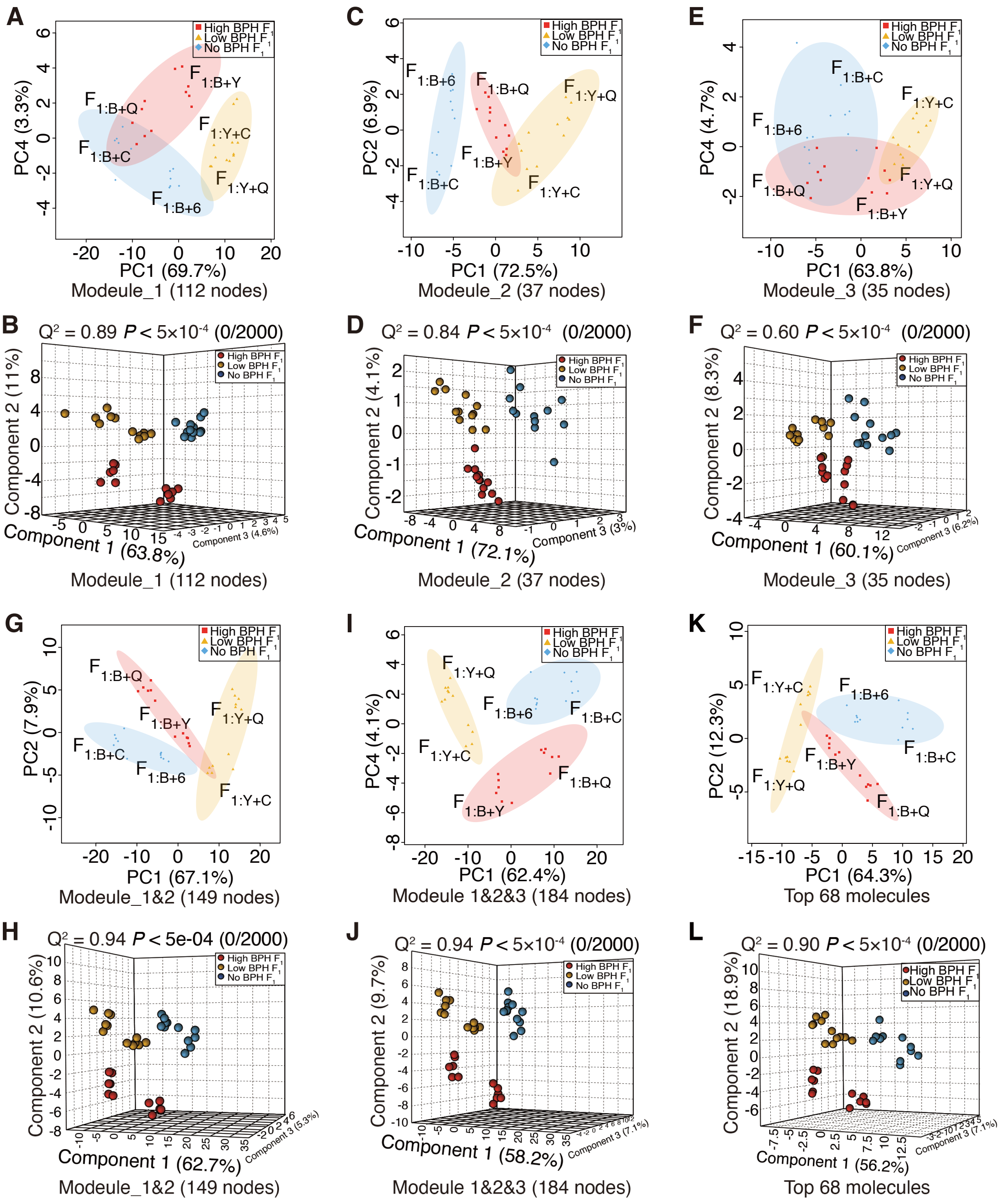


**Supplemental Figure 14. Identification of top-contributing molecules from Subnetwork_1 modules in Network_HLN.** PCA and PLS-DA score plots of six pairs of reciprocal $F_1$ hybrids based on molecular levels of: Module_1 (**A-B**), Module_2 (**C-D**), Module_3 (**E-F**), Module_1 + Module_2 (**G-H**), integrated Module_1-3 (**I-J**), and the top 68 molecules (**K-L**) are shown. *P* values are for permutation test.

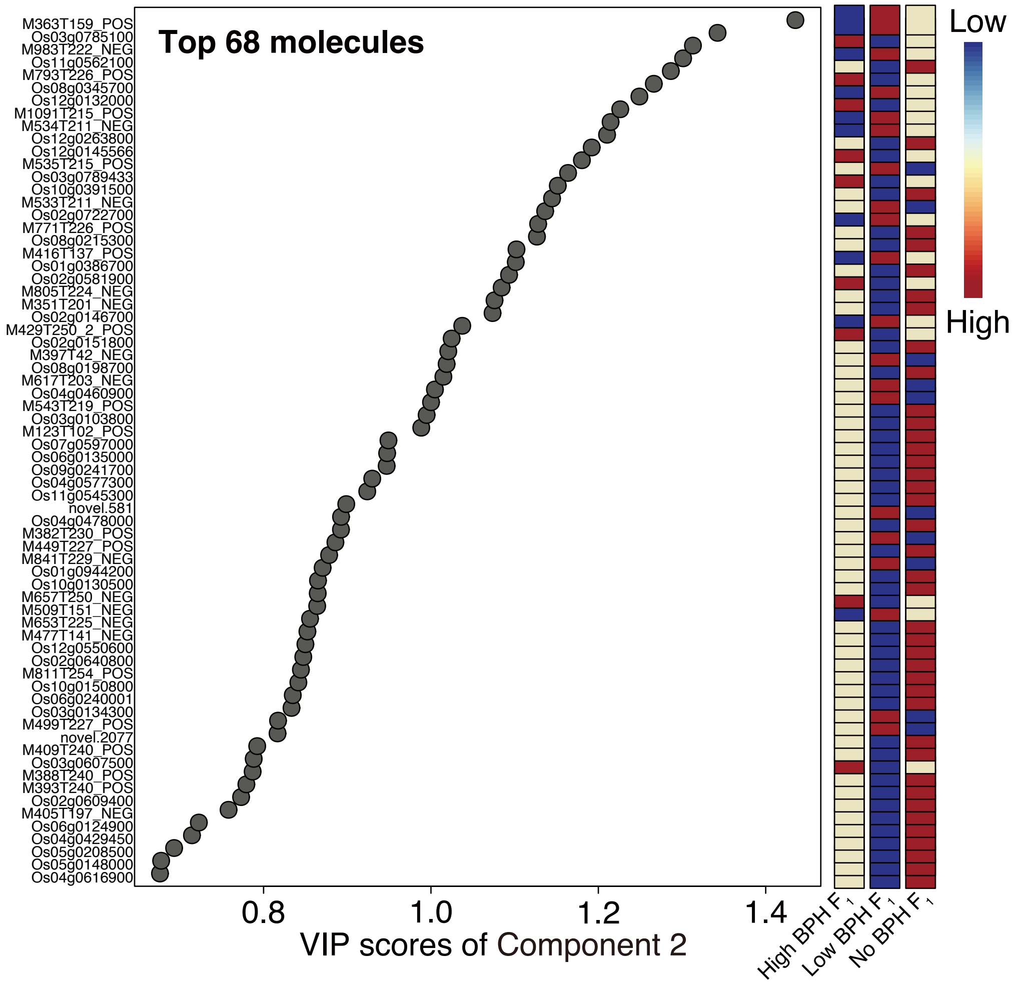


**Supplemental Figure 15. Scores of variable importance in projection (VIP) for Component 2 in partial least squares-discriminant analysis.** Molecular levels are shown for the top 68 heterosis-associated molecules across high-, low-, and no better-parent heterosis (BPH) groups.

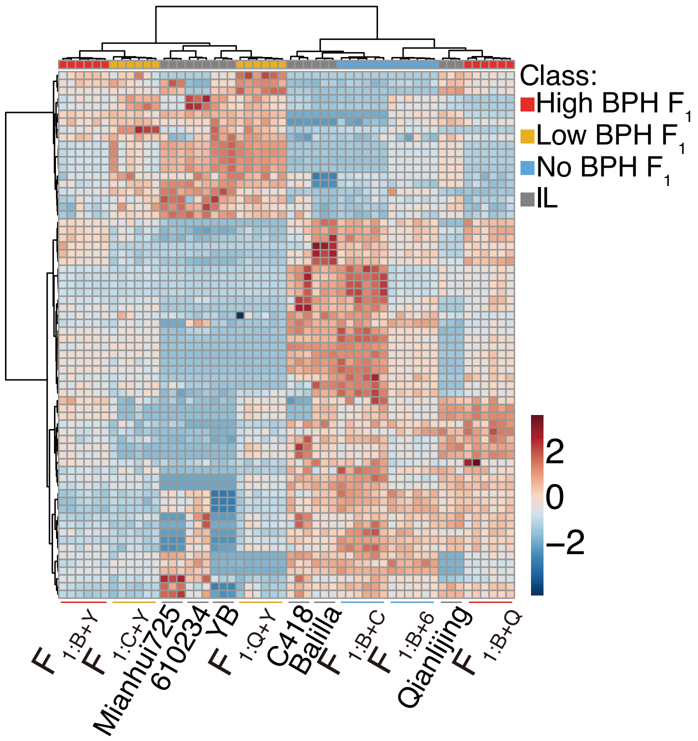


**Supplemental Figure 16. Heatmap of expression level/abundance for 68 heterosis-associated molecules in six inbred lines and $F_1$ hybrids.**

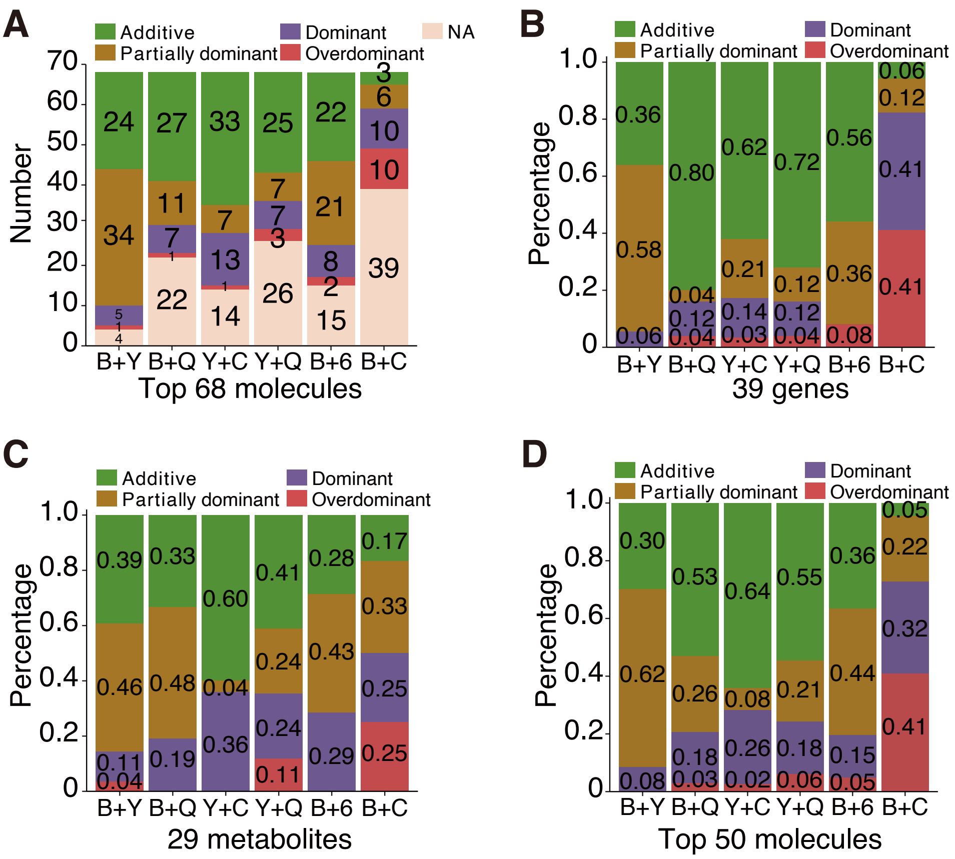


**Supplemental Figure 17. Inheritance patterns of heterosis-associated molecules for seedling length.**

**(A)** Count of inheritance patterns for 68 heterosis-associated molecules.

**(B-C)** Percentage distribution of inheritance patterns for 39 heterosis-associated genes (**B**) and 29 heterosis-associated metabolites (**C**).

**(D)** Percentage distribution of inheritance patterns for the top 50 heterosis-associated molecules.

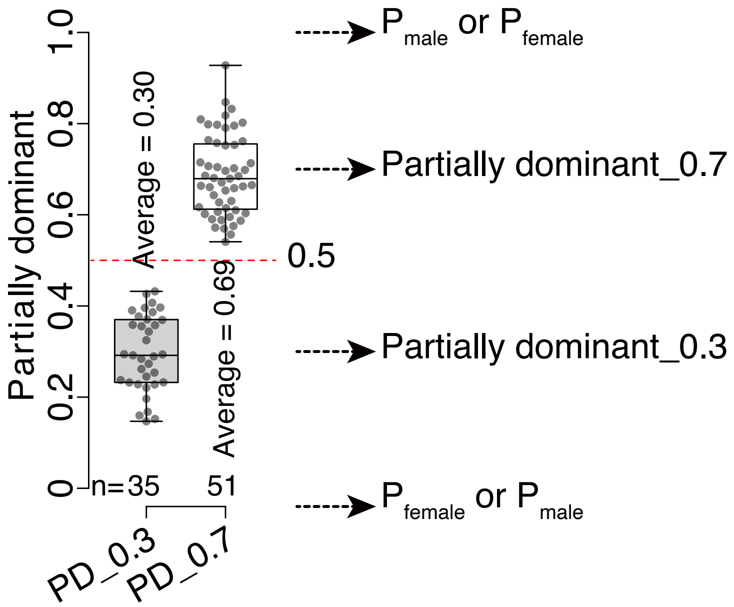


**Supplemental Figure 18. Classification of partially dominant effects.** Using a threshold of 0.5, partially dominant (PD) effect was subdivided into PD_0.3 and PD_0.7 subtypes. The associations of PD_0.3, PD_0.7, and parental molecular levels (female and male parents) are indicated.

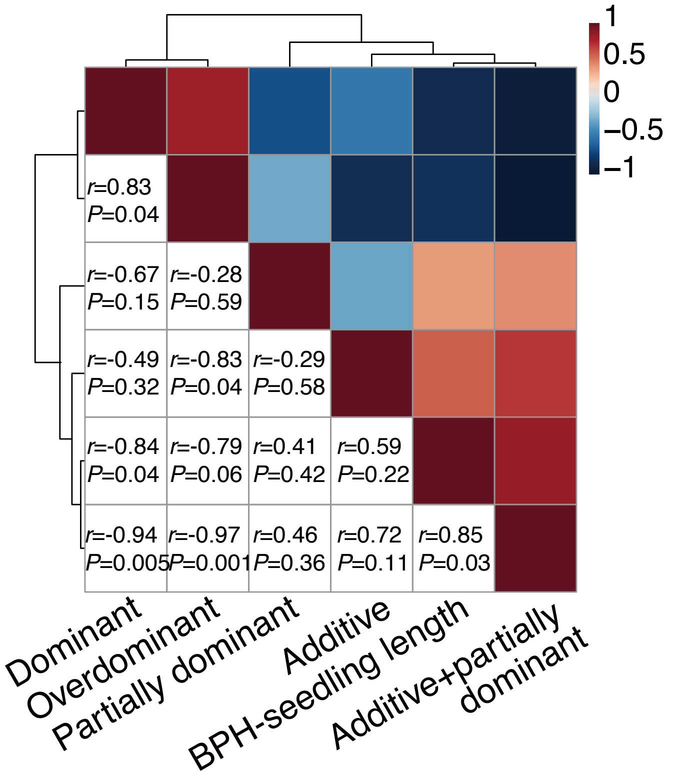


**Supplemental Figure 19. Heatmap for Pearson correlations of better-parent heterosis (BPH) for seedling length and the percentages of inheritance patterns.** *P* values indicate Pearson correlation.

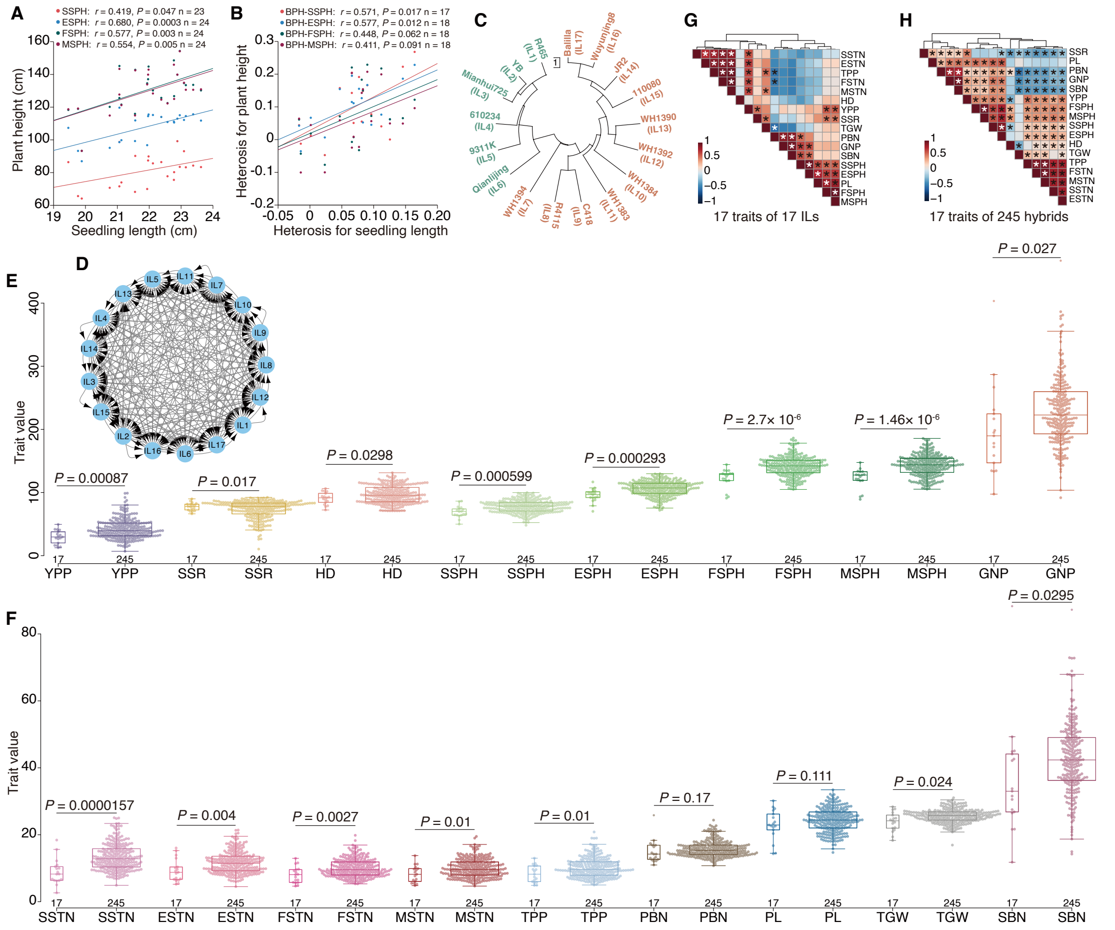

**Supplemental Figure 20. Trait correlations and values for 17 agronomic traits in parental lines and $F_1$ hybrids (a diallel-cross population).**

**(A)** Pearson correlations between seedling length (15 days after sowing) and plant height (across four developmental stages) in parental inbred lines (ILs) and $F_1$ hybrids.

**(B)** Pearson correlations between seedling length heterosis and plant height heterosis. The $F_1$ hybrid C418×Qianlijing was excluded due to large reciprocal differences.

**(C)** Neighbour-joining tree of 17 ILs based on InDel markers.

**(D)** Diallel cross design of the 17 ILs.

**(E-F)** Values of 17 agronomic traits in 17 ILs and 245 $F_1$ hybrids. *P* values indicate independent samples *t* tests.

**(G-H)** Heatmaps for correlations of 17 agronomic traits in 17 ILs (**G**) and 245 $F_1$ hybrids (**H**). Asterisks indicate significant Pearson correlations ($P < 0.05$). Abbreviations: better-parent heterosis = BPH, yield per plant = YPP, secondary branch number = SBN, grain number per panicle = GNP, tiller number per plant = TPP, seed setting rate = SSR, heading date = HD, thousand grain weight = TGW, seedling stage tiller number = SSTN, elongation stage tiller number = ESTN, flowering stage tiller number = FSTN, maturation stage tiller number = MSTN, seedling stage plant height = SSPH, elongation stage plant height = ESPH, flowering stage plant height = FSPH, maturation stage plant height = MSPH, panicle length = PL, and primary branch number = PBN.

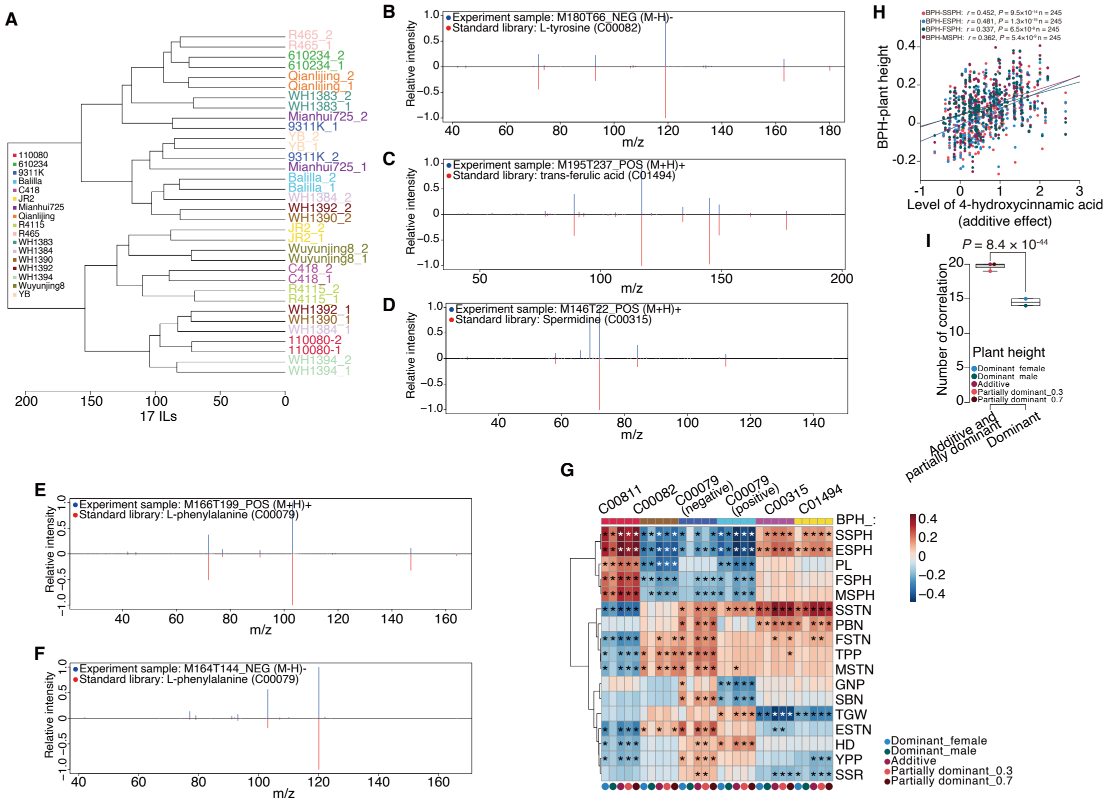

**Supplemental Figure 21. Metabolomic analysis of 17 inbred lines (ILs) and associations between phenylpropanoid biosynthesis metabolites and heterosis of 17 agronomic traits.**

**(A)** Dendrogram of 17 ILs based on untargeted metabolite profiles (two biological replicates per line).

**(B-F)** MS/MS spectra of four metabolites from phenylpropanoid biosynthesis. Experimental sample spectra are shown above the horizontal line; chemical standard spectra are shown below. L-phenylalanine was annotated in both positive and negative ion modes.

**(G)** Heatmap showing correlations between different inheritance patterns (additive, partially dominant, dominant effects) of five chemically annotated metabolites and heterosis of 17 agronomic traits. Asterisks indicate significant Pearson correlations ($P < 0.05$).

**(H)** Correlations between the levels of 4-hydroxycinnamic acid (additive effect) and heterosis for plant height across developmental stages. *P* values are for Pearson correlation.

**(I)** Number of significant correlations between different inheritance patterns (additive/partially dominant vs dominant effects) of five chemically annotated metabolites from phenylpropanoid biosynthesis and heterosis for plant height across four developmental stages. *P* value indicates independent samples *t* tests (weighted). Abbreviations: better-parent heterosis = BPH, yield per plant = YPP, secondary branch number = SBN, grain number per panicle = GNP, tiller number per plant = TPP, seed setting rate = SSR, heading date = HD, thousand grain weight = TGW, seedling stage tiller number = SSTN, elongation stage tiller number = ESTN, flowering stage tiller number = FSTN, maturation stage tiller number = MSTN, seedling stage plant height = SSPH, elongation stage plant height = ESPH, flowering stage plant height = FSPH, maturation stage plant height = MSPH, panicle length = PL, and primary branch number = PBN.

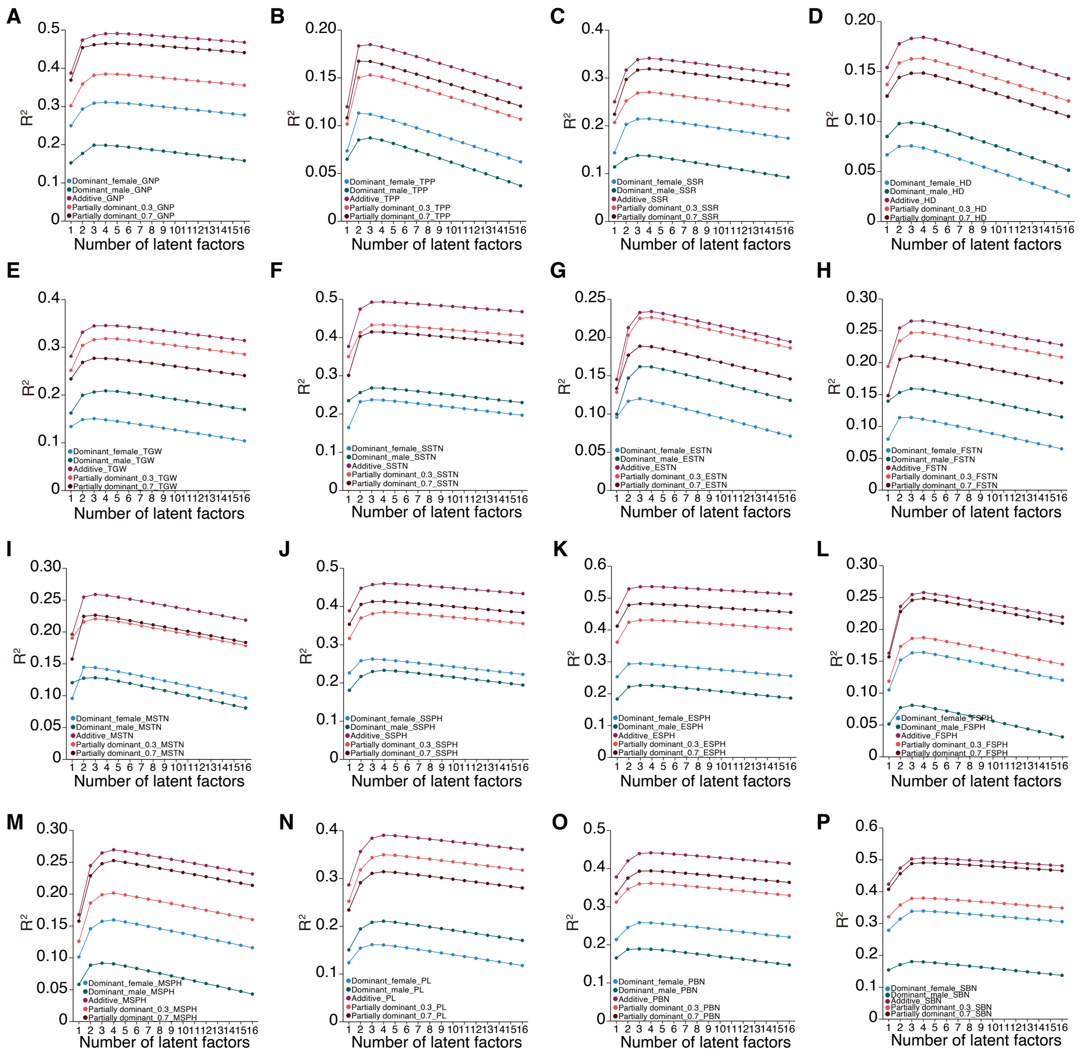


**Supplemental Figure 22. $R^2$ values with varying number of latent factors in partial least squares analysis on heterosis of 16 agronomic traits in 245 $F_1$ hybrids.** The regression models used 1,306 differential metabolites from 17 inbred lines as predictors for better-parent heterosis of 16 traits. Abbreviations: secondary branch number = SBN, grain number per panicle = GNP, tiller number per plant = TPP, seed setting rate = SSR, heading date = HD, thousand grain weight = TGW, seedling stage tiller number = SSTN, elongation stage tiller number = ESTN, flowering stage tiller number = FSTN, maturation stage tiller number = MSTN, seedling stage plant height = SSPH, elongation stage plant height = ESPH, flowering stage plant height = FSPH, maturation stage plant height = MSPH, panicle length = PL, and primary branch number = PBN.

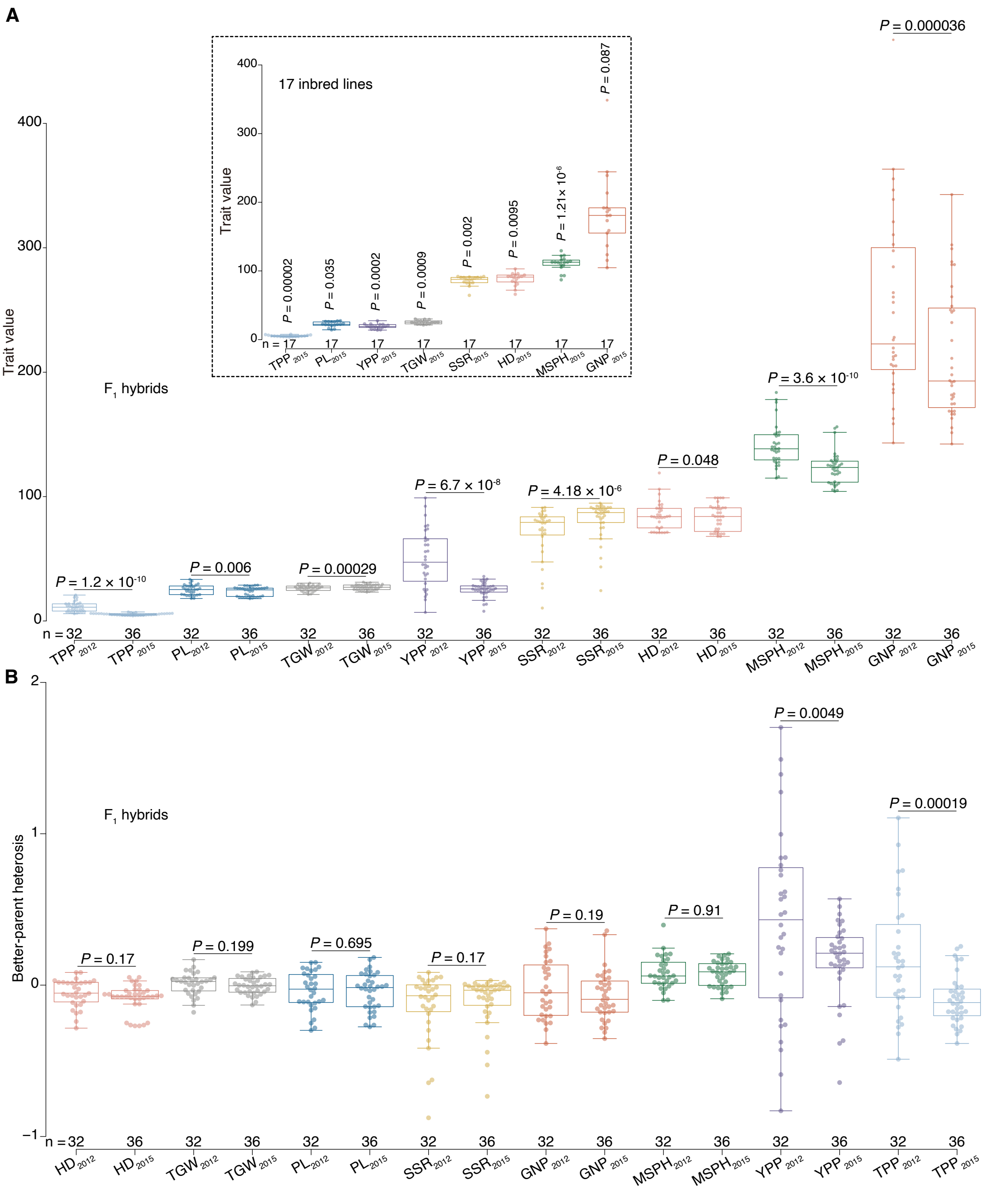


**Supplemental Figure 23. Trait values and better-parent heterosis of eight traits across growth conditions.**

**(A)** Values of eight traits measured in 2012 and 2015 for 17 inbred lines and 36 $F_1$ hybrids. Trait values were available for only 32 $F_1$ hybrids in 2012.

**(B)** Better-parent heterosis of eight traits measured in 2012 and 2015. *P* values are for independent samples *t* tests. Abbreviations: heading date = HD, thousand grain weight = TGW, panicle length = PL, seed setting rate = SSR, grain number per panicle = GNP, maturation stage plant height = MSPH, grain yield per plant = YPP, and tiller number per plant = TPP.

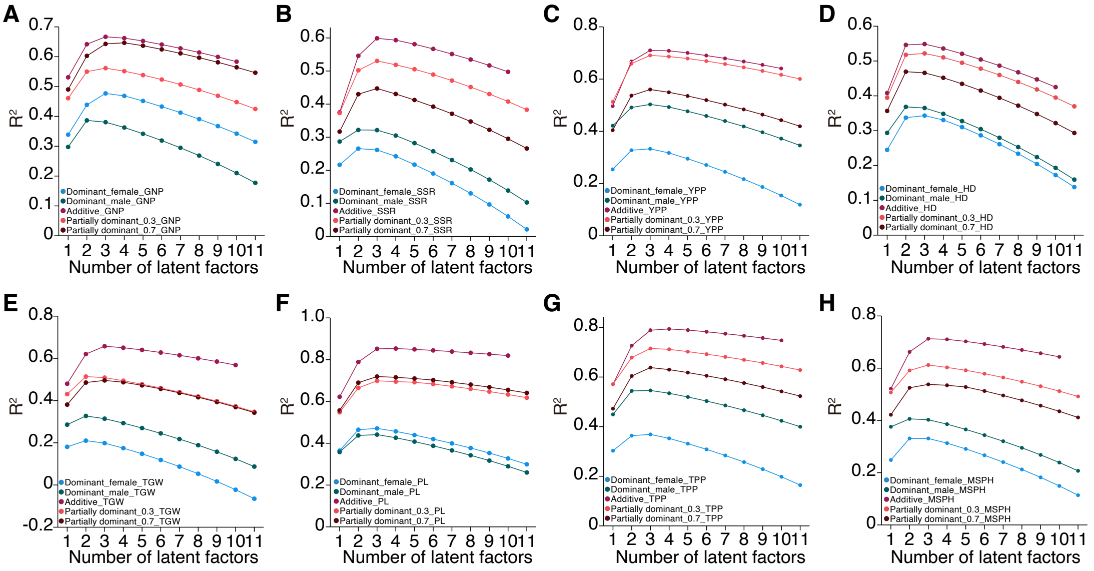


**Supplemental Figure 24. $R^2$ values with varying number of latent factors in partial least squares analysis on heterosis of eight agronomic traits in 36 $F_1$ hybrids.** The regression models based on 1,306 differential metabolites (three inheritance patterns) predicted heterosis of eight traits measured in 2015. The $R^2$ values (adjusted) with 1-11 latent factors are shown for better-parent heterosis for grain number per panicle (GNP; **A**), seed setting rate (SSR; **B**), grain yield per plant (YPP; **C**), heading date (HD; **D**), thousand grain weight (TGW; **E**), panicle length (PL; **F**), tiller number per plant (TPP; **G**), and maturation stage plant height (MSPH; **H**).

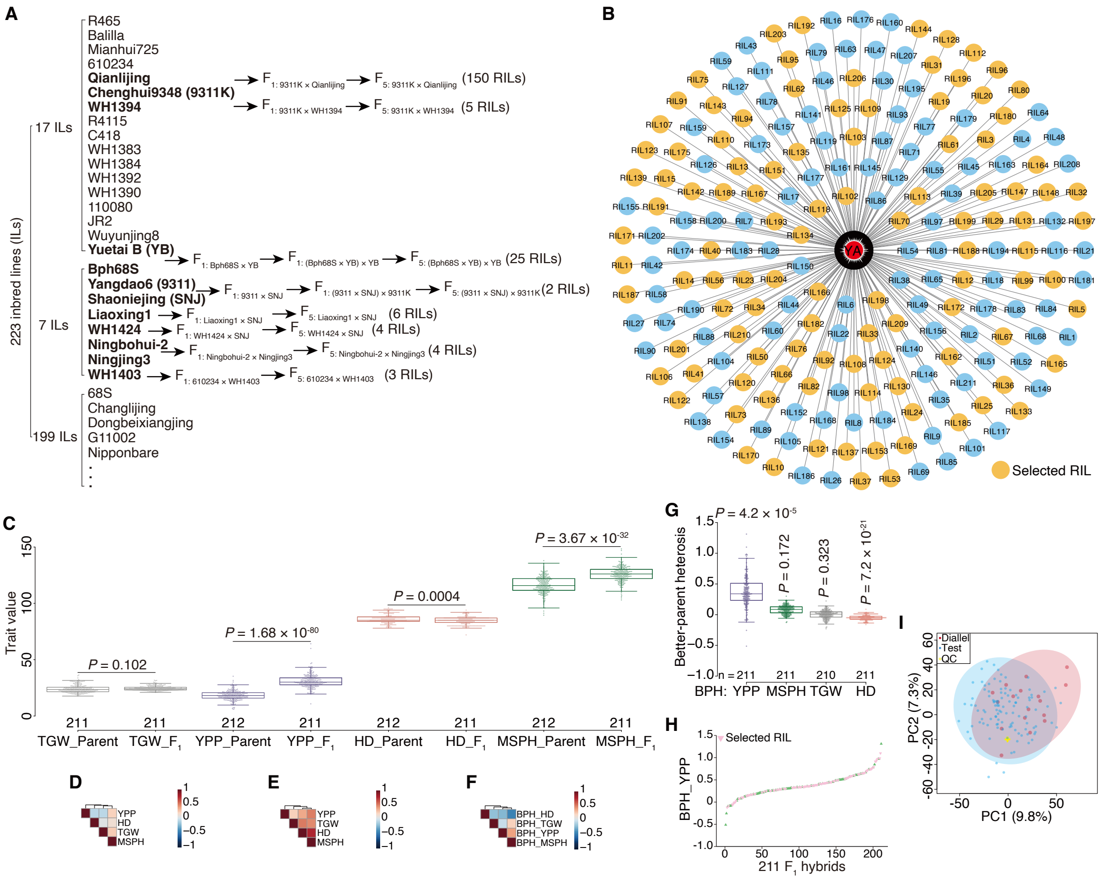


**Supplemental Figure 25. Information of test-cross population and its connections to the diallel-cross population.**

**(A)** Breeding scheme of 211 recombinant inbred lines (RILs) from 223 inbred lines (ILs) via five generations of self-crossing. Male parents of the test-cross population also contained 12 RILs without parental information.

(**B**) Design of test cross population. A total of 211 RILs were crossed with a Honglian-type cytoplasmic male sterile line Yuetai A (YA). RILs selected for metabolite profiling analysis are highlighted in yellow.

**(C)** Value of four traits for test-cross $F_1$ hybrids and parents. Due to sterile character of YA, trait values of its restorer line (Yuetai B) were used for analysis. *P* values are for independent samples *t* tests.

**(D-F)** Heatmaps for correlations of four traits in parental lines (**D**), test-cross $F_1$ hybrids (**E**), and better-parent heterosis in $F_1$ hybrids (**F**). Pearson correlation was performed.

**(G)** Better-parent heterosis (BPH) of four traits. *P* values indicate independent samples *t* tests between the diallel- and test-cross populations.

**(H)** Better-parent heterosis for grain yield per plant in test-cross $F_1$ hybrids from 211 RILs. The RILs selected for metabolite profiling analysis are marked in pink.

**(I)** PCA score plot of parent lines for the diallel- (17 ILs) and test-cross (107 RILs and Yuetai A) populations based on untargeted metabolomic data. Abbreviations: grain yield per plant = YPP, maturation stage plant height = MSPH, thousand grain weight = TGW, heading date = HD, and quality control = QC.

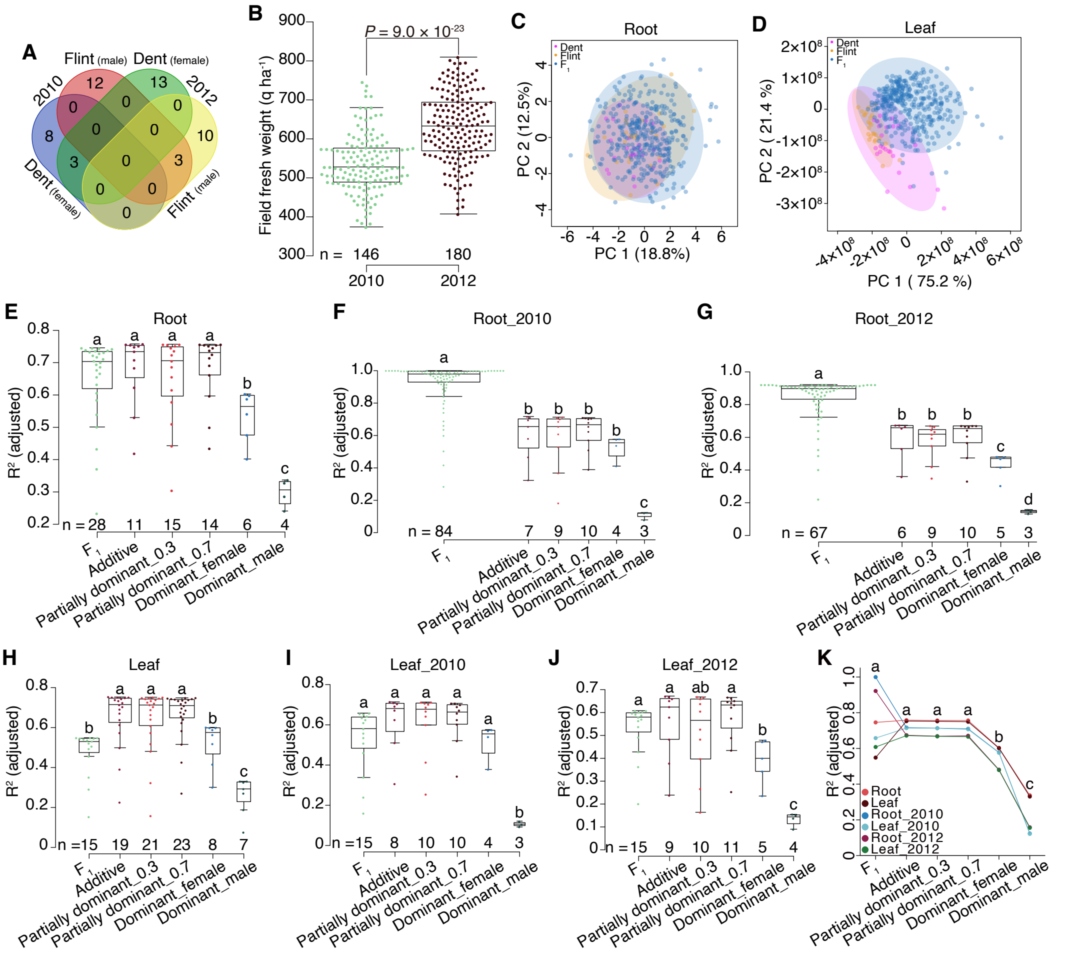


**Supplemental Figure 26. Predicting field fresh weight in 326 maize $F_1$ hybrids using metabolomic data and partial least squares (PLS).**

**(A)** Venn diagram of 24 Dent (female) and 25 Flint (male) lines for two hybrid populations phenotyped in 2010 (n = 146 hybrids) and 2012 (n =180 hybrids).

**(B)** Field fresh weight of the two maize hybrid populations. *P* value indicates independent samples *t* test.

**(C-D)** PCA score plots of 24 Dent lines, 25 Flint lines, and their 326 $F_1$ hybrids (Dent × Flint) based on metabolomic data from young roots and leaves.

**(E-G)** $R^2$ values of PLS-based models for predicting field fresh weight using hybrid metabolite profiles ($F_1$) and different inheritance patterns from young root data (165 metabolites). Panels show combined populations (**E**), 2010 population (**F**) and 2012 population (**G**). n = number of latent factors.

**(H-J)** $R^2$ values of PLS-based models for predicting field fresh weight using leaf metabolic data (81 metabolites). Panels show combined populations (**H**), 2010 population (**I**), and 2012 population (**J**).

**(K)** Top $R^2$ values of models predicting field fresh weight with hybrid profiles and different inheritance patterns using metabolomic data of roots and leaves. Analysis of variance was performed with the least significant difference used in post hoc test.

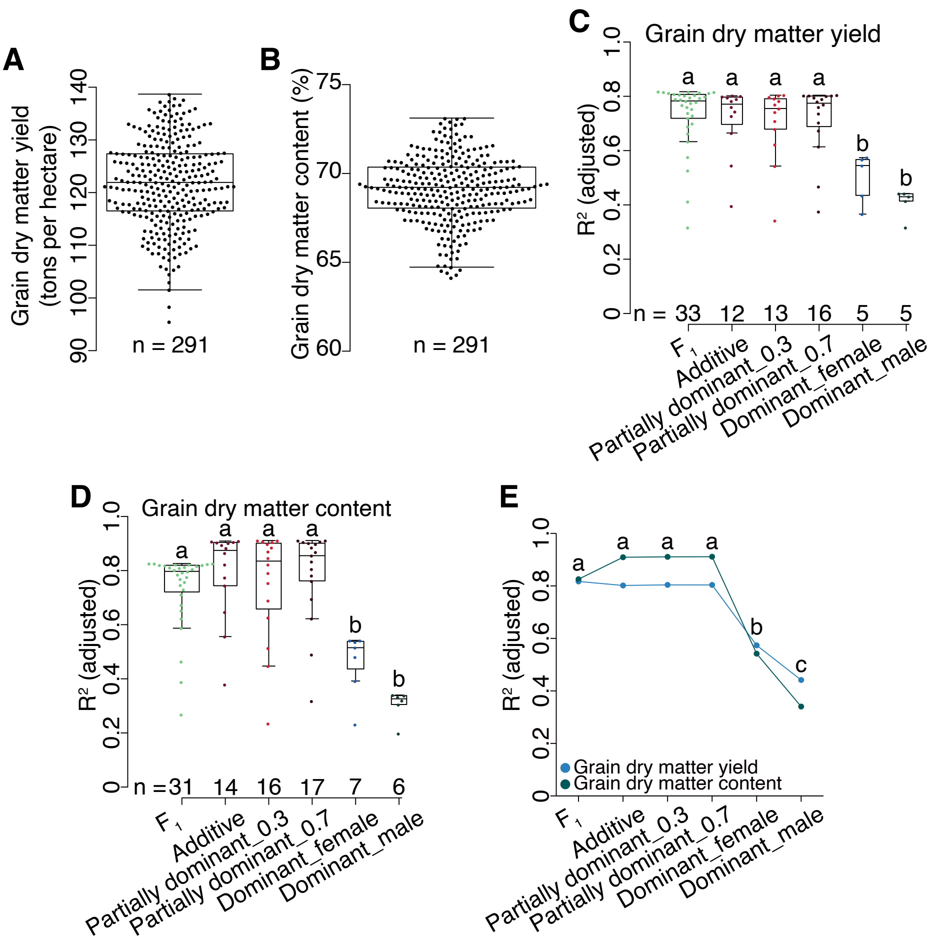


**Supplemental Figure 27. Predicting grain dry matter yield and content in 291 maize $F_1$ hybrids using metabolomic data and partial least squares (PLS).**

**(A-B)** Grain dry matter yield and grain day matter content of 291 $F_1$ hybrids.

**(C-D)** Values of $R^2$ for predicting grain dry matter yield (**C**) and grain day matter content (**D**) using hybrid profiles ($F_1$) and different inheritance patterns of root metabolic data (165 metabolites). n = number of latent factors.

**(E)** Top $R^2$ values for predicting grain dry matter yield and grain dry matter content using root metabolic data. Analysis of variance was performed with the least significant difference used in post hoc test.

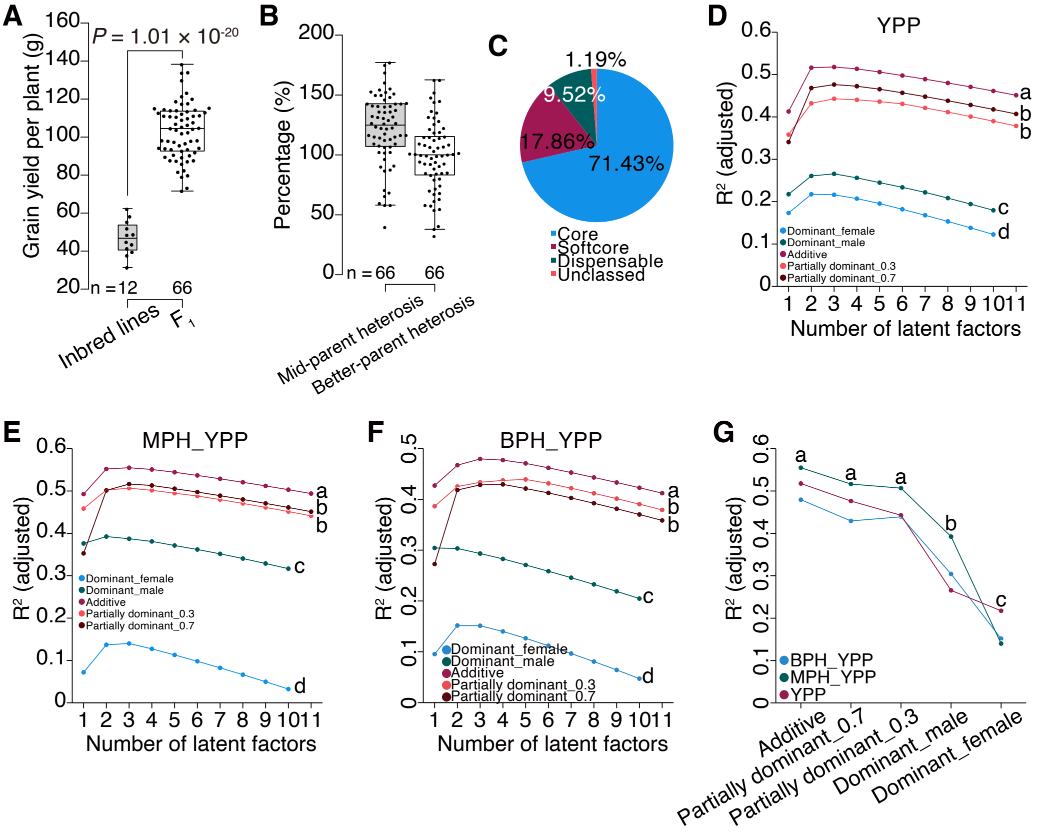


**Supplemental Figure 28. Predicting grain yield and heterosis for grain yield in 66 maize $F_1$ hybrids using gene expression data and partial least squares (PLS).**

**(A)** Grain yield per plant of 12 inbred lines and 66 $F_1$ hybrids. *P* value indicates independent samples *t* test.

**(B)** Mid-parent and better-parent heterosis for grain yield per plant.

**(C)** Distribution of 84 differentially expressed genes (classified by pan-genome) in 12 inbred lines.

**(D-F)** Values of $R^2$ with varying number of latent factors in predicting grain yield per plant (YPP; **D**), mid-parent heterosis for grain yield per plant (MPH_YPP; **E**), and better-parent heterosis for grain yield per plant (BPH_YPP; **F**) using different inheritance patterns of 84 genes.

**(G)** Top $R^2$ values for predicting YPP, MPH_YPP, and BPH_YPP. Analysis of variance was performed with the least significant difference used in post hoc test.

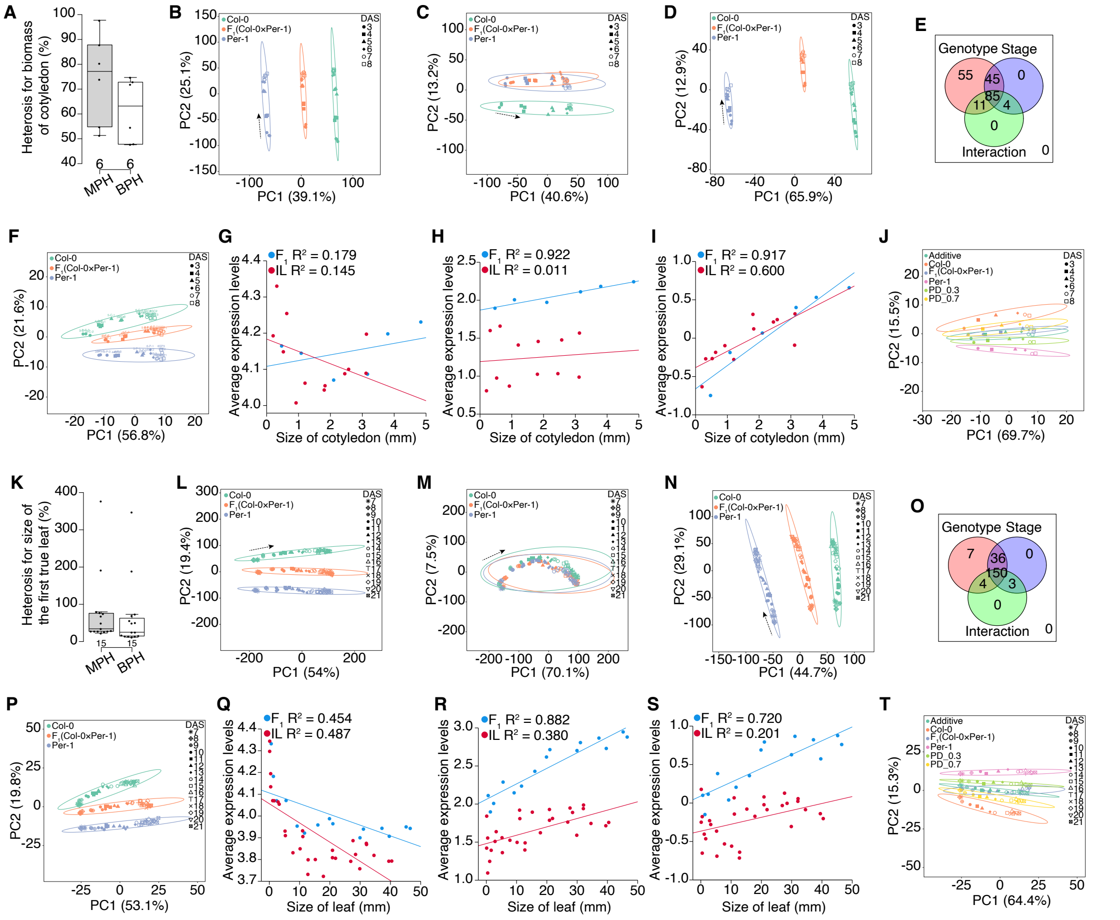

**Supplemental Figure 29. Analyses of transcriptomic and phenotypic data in *Arabidopsis*.**

**(A)** Mid-parent heterosis (MPH) and better-parent heterosis (BPH) for cotyledon biomass across stages in Col-0 × Per-1 $F_1$ hybrids.

(**B-D**) PCA score plots of Col-0, Per-1, and their $F_1$ hybrids based on cotyledon gene expression data at 3-8 days after sowing (DAS). All genes (16,109 genes; **B**), 15,179 non-differentially expressed genes (NDEGs; **C**), and 930 differentially expressed genes (DEGs; **D**) are analyzed.

**(E)** Analysis of variance of 930 DEGs responsive to genotype, stage, and their interaction.

(**F**) PCA score plots using 31 heterosis-associated genes for cotyledon biomass.

**(G-I)** Correlations between cotyledon size and average gene expression levels for all 16,109 genes (**G**), 930 genes (**H**), and 31 genes (**I**). $R^2$ values indicate Pearson correlation.

(**J**) PCA score plots of Col-0, Per-1, their $F_1$ hybrids, "additive hybrids", and "partially dominant hybrids" using 31 genes.

**(K)** MPH and BPH for size of the first true leaf across stages in Col-0 × Per-1 $F_1$ hybrids.

(**L-N**) PCA score plots based on expression data of leaves at 7-21 DAS using all genes (16,384 genes; **L**), 14,839 NDEGs (**M**), and 1,545 DEGs (**N**).

**(O)** Analysis of variance of 1,545 DEGs.

(**P**) PCA score plots using 91 heterosis-associated genes.

**(Q-S)** Correlations between leaf size and average expression levels of all 16,384 genes (**Q**), 1,545 DEGs (**R**), and 91 genes (**S**).
(**T**) PCA score plots of Col-0, Per-1, their $F_1$ hybrids, "additive hybrids", and "partially dominant hybrids" using 91 genes.

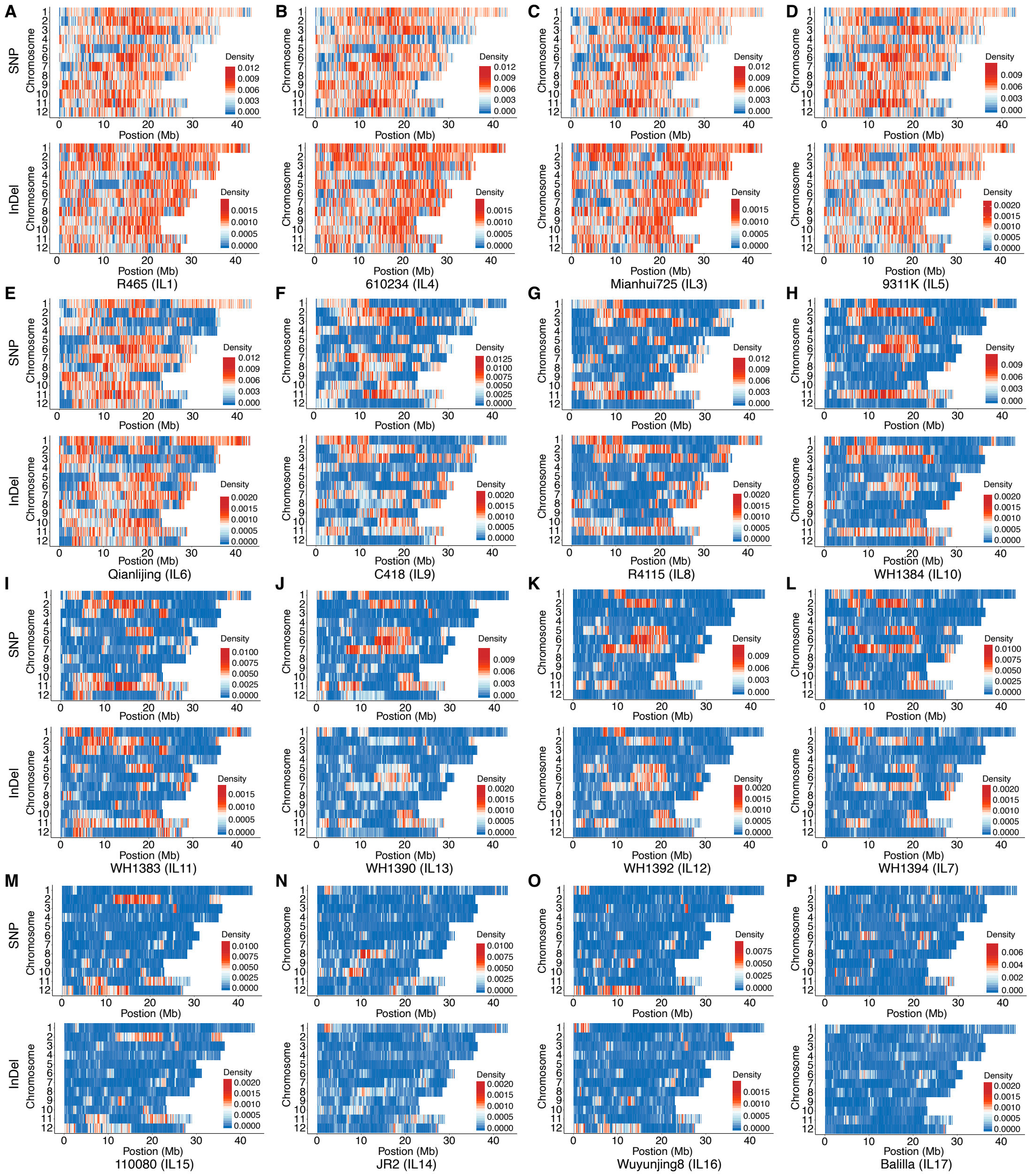


**Supplemental Figure 30. SNP and InDel density across 12 chromosomes in 16 rice inbred lines (ILs).** Heatmaps show SNP and InDel density for: R465 (IL1; **A**), 610234 (IL4; **B**), Mianhui725 (IL3; **C**), 9311K (IL5; **D**), Qianlijing (IL6; **E**), C418 (IL9; **F**), R4115 (IL8; **G**), WH1384 (IL10; **H**), WH1383 (IL11; **I**), WH1390 (IL13; **J**), WH1392 (IL12; **K**), WH1394 (IL7; **L**), 110080 (IL15; **M**), JR2 (IL14; **N**), Wuyunjing8 (IL16; **O**), and Balilla (IL17; **P**).

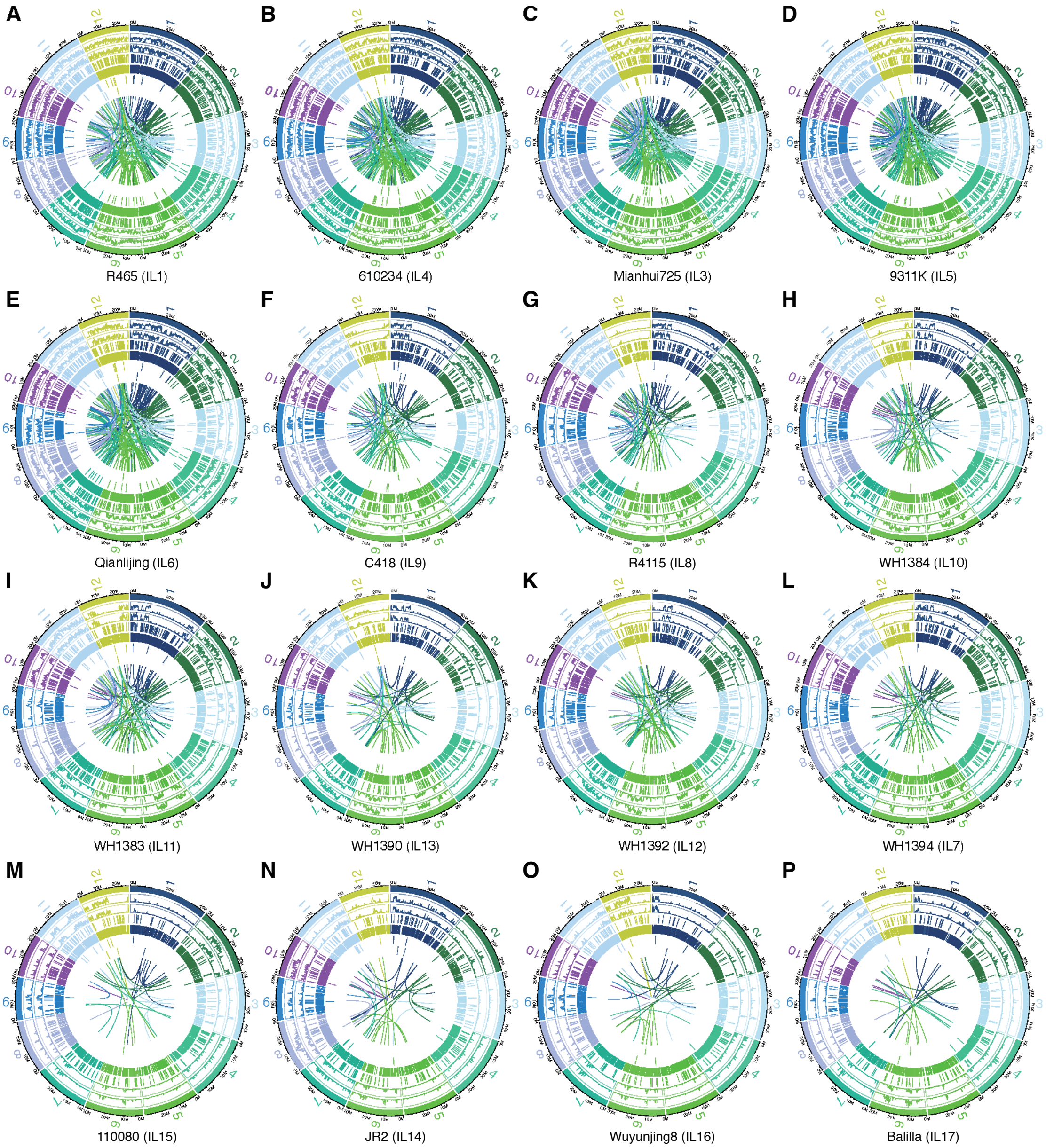


**Supplemental Figure 31. Circos visualization of genomic variants in 16 rice inbred lines (ILs).** Variants (SNPs, InDels, CNVs, and SVs) were identified by genomic sequencing analysis using the Nipponbare reference genome. Circos plots display variants for the following ILs: R465 (IL1; **A**), 610234 (IL4; **B**), Mianhui725 (IL3; **C**), 9311K (IL5; **D**), Qianlijing (IL6; **E**), C418 (IL9; **F**), R4115 (IL8; **G**), WH1384 (IL10; **H**), WH1383 (IL11; **I**), WH1390 (IL13; **J**), WH1392 (IL12; **K**), WH1394 (IL7; **L**), 110080 (IL15; **M**), JR2 (IL14; **N**), Wuyunjing8 (IL16; **O**), and Balilla (IL17; **P**). From outer to inner circles: chromosome position, SNP density, InDel density, CNV duplication, CNV deletion position, SV insertion position, SV deletion position, and SV inversion position.

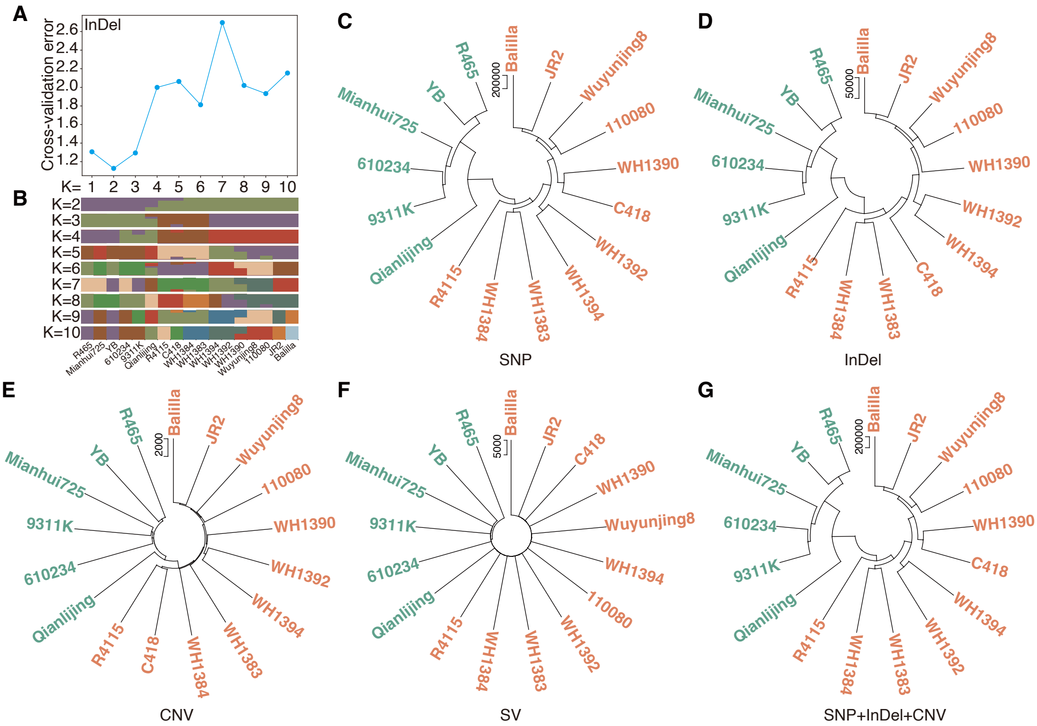


**Supplemental Figure 32. Population structure and neighbor-joining trees of 17 rice inbred lines based on genomic variants.**

**(A)** Cross-validation errors for K values ranging from 1 to 10 using InDels.

**(B)** Population structure inferred from InDels.

**(C-G)** Neighbor-joining trees based on different types of genomic variants. Genetic distances of the 17 inbred lines were represented by the number of SNPs (**C**), InDels (**D**), CNVs (**E**), SVs (**F**), and combined SNP+InDel+CNV (**G**).

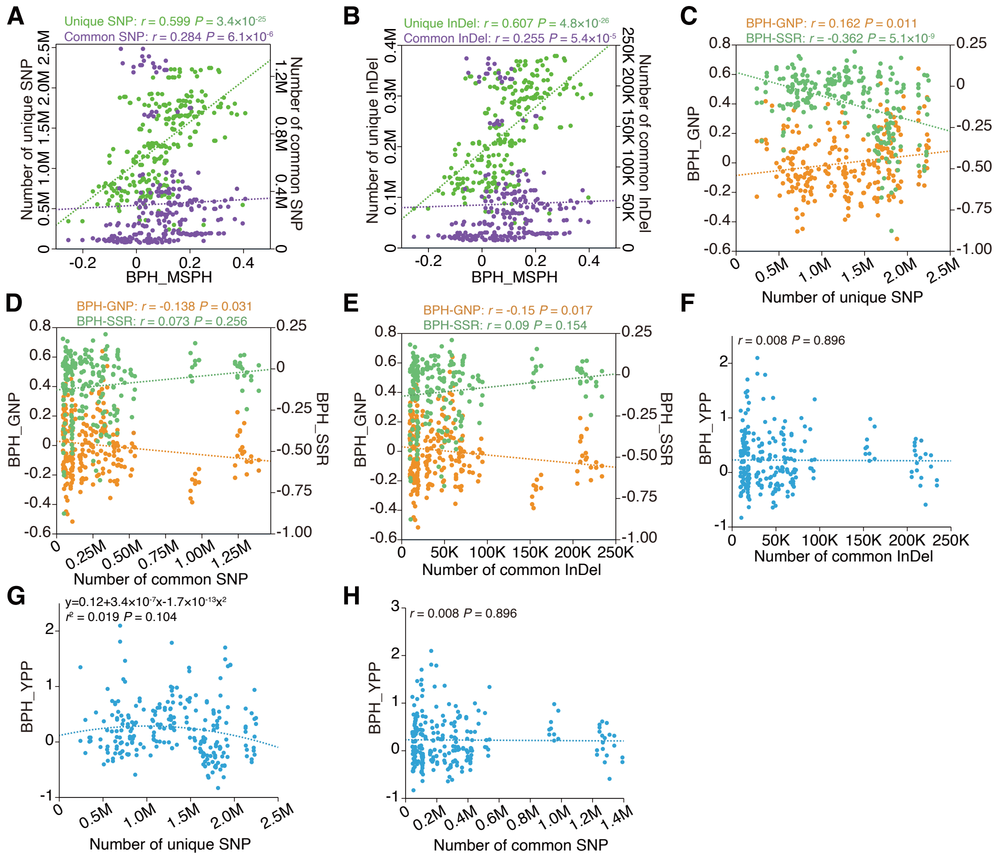


**Supplemental Figure 33. Correlations between parental genomic variants and heterosis of agronomic traits in rice.**

**(A-B)** Correlations of better-parent heterosis for plant height at maturation stage (BPH_MSPH) with the number of parental common and unique SNPs (**A**) or InDels (**B**).

**(C-D)** Correlations of the number of parental unique (**C**) or common (**D**) SNPs with better-parent heterosis for grain number per panicle (BPH_GNP) and better-parent heterosis for seed setting rate (BPH_SSR).

**(E)** Correlations of the number of parental common InDels with heterosis for grain number per plant and heterosis for seed setting rate.

**(F)** Correlation between the number of parental common InDels and better parent heterosis for grain yield per plant (BPH_YPP).

**(G-H)** Correlations between the number of parental unique (**G**) or common (**H**) SNPs and better-parent heterosis for grain yield per plant (BPH_YPP). *P* values are for Spearman rank correlation in **A-E**, **F**, and **H**. *P* value is for quadratic regression in **G**.

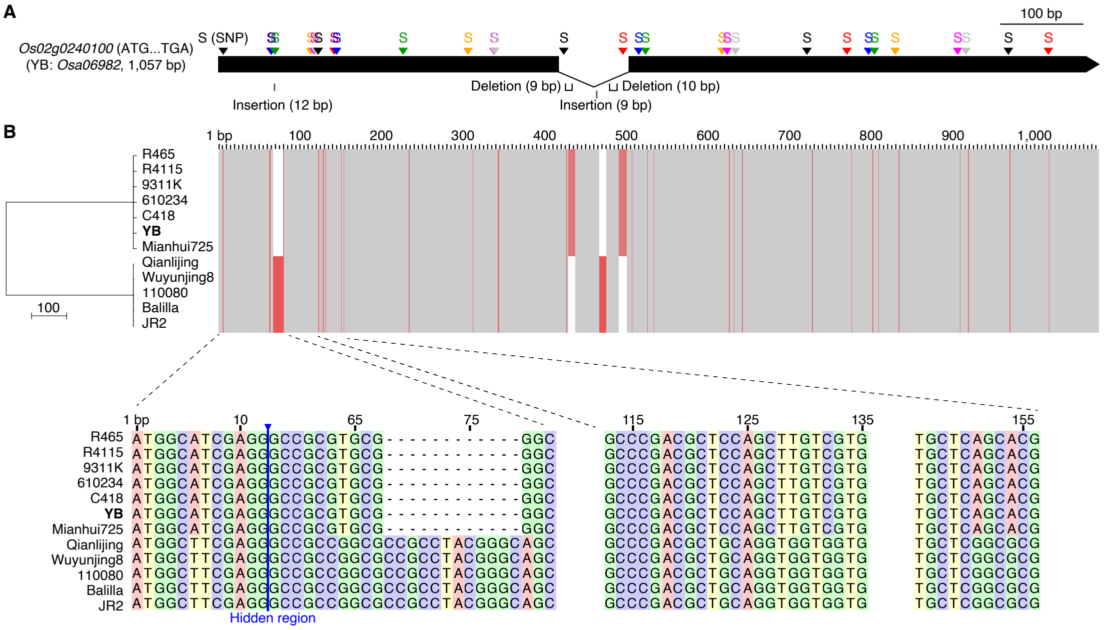


**Supplemental Figure 34. Variation profiles of *Os02g0240100* in 12 rice inbred lines.**

**(A)** Gene structure of *Os02g0240100*. With Yuetai B (YB) as the reference, genomic variants (SNPs, insertions, deletions) detected in 11 inbred lines are indicated.

**(B)** Clustering of the 12 inbred lines based on sequence variants of *Os02g0240100*. Red blocks represent the presence of variants. Detailed sequences of two specific regions in the first exon are shown.

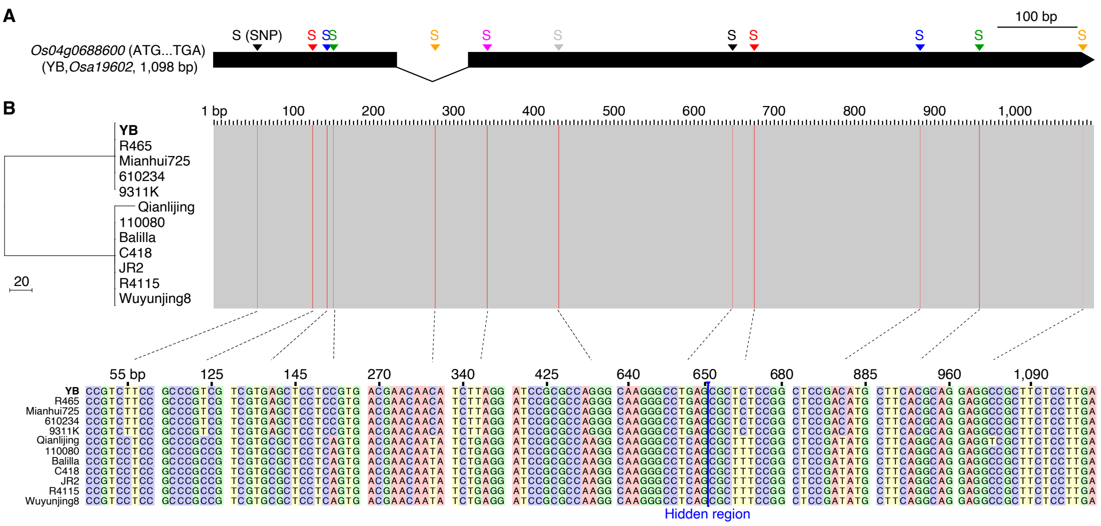


**Supplemental Figure 35. Variation profiles of *Os04g0688600* in 12 rice inbred lines.**

**(A)** Gene structure of *Os04g0688600*. With Yuetai B (YB) as the reference, 12 SNPs detected in 11 inbred lines are indicated.

**(B)** Clustering of the 12 inbred lines based on sequence variants of *Os04g0688600*. Red blocks represent the presence of SNPs.

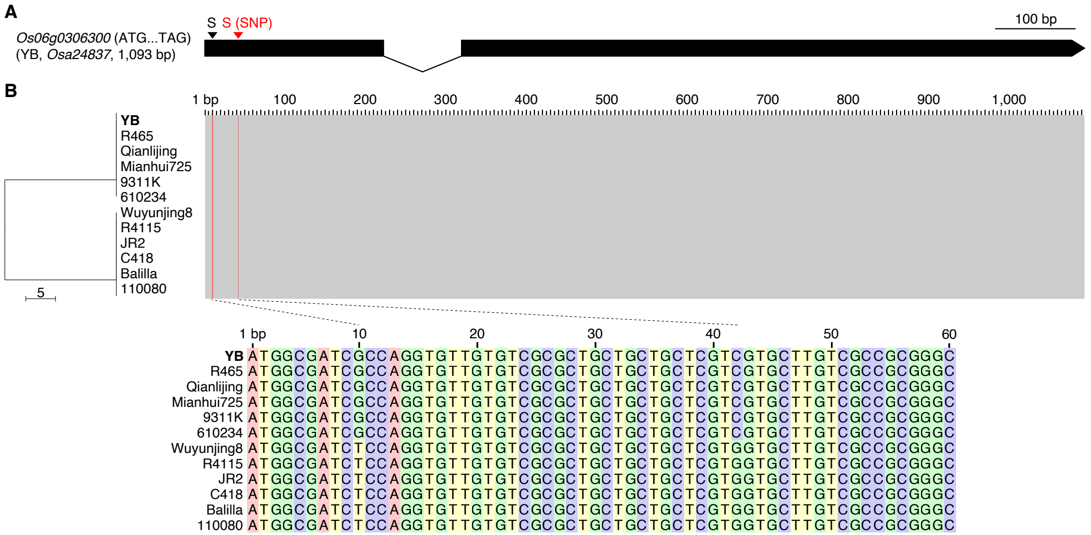


**Supplemental Figure 36. Variation profiles of *Os06g0306300* in 12 rice inbred lines.**

**(A)** Gene structure of *Os06g0306300*. With Yuetai B (YB) as the reference, two SNPs detected in 11 inbred lines are indicated.

**(B)** Clustering of the 12 inbred lines based on sequence variants of *Os06g0306300*. Red blocks represent the presence of SNPs.

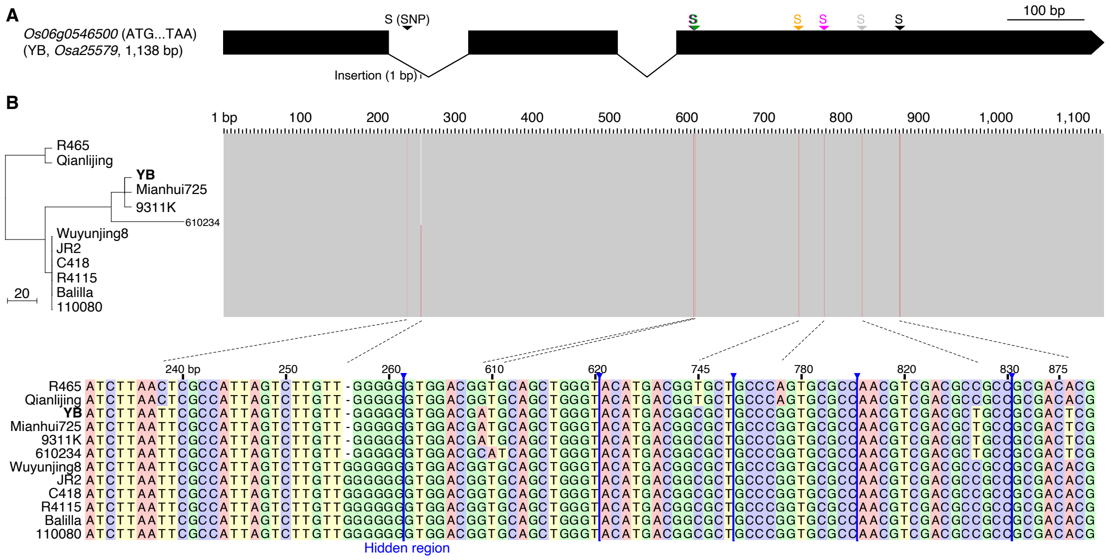


**Supplemental Figure 37. Variation profiles of *Os06g0546500* in 12 rice inbred lines.**

**(A)** Gene structure of *Os06g0546500*. With Yuetai B (YB) as the reference, genomic variant detected in 11 inbred lines are indicated.

**(B)** Clustering of the 12 inbred lines based on sequence variants of *Os06g0546500*. Red blocks represent the presence of variants.

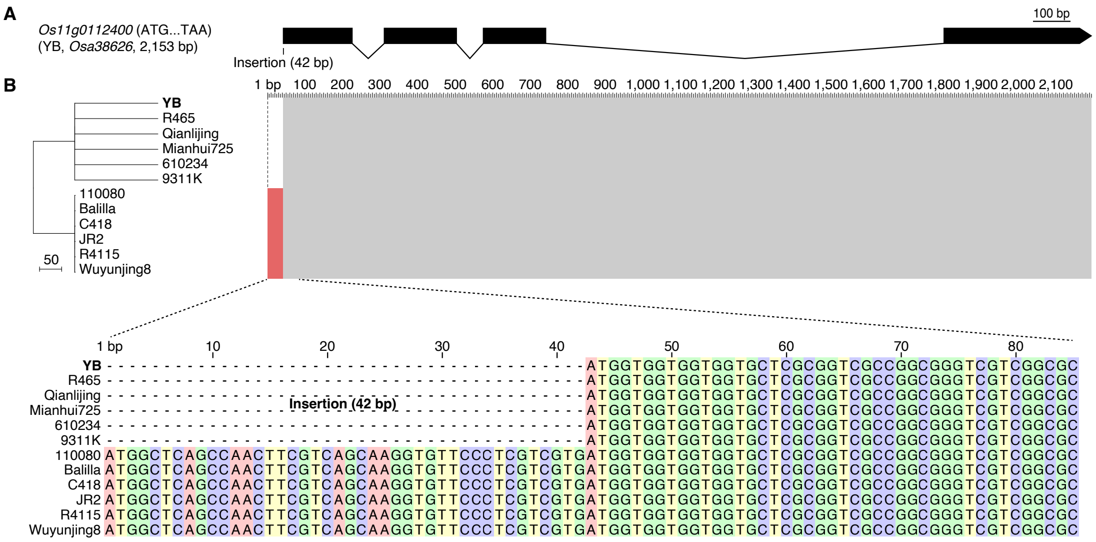


**Supplemental Figure 38. Variation profiles of *Os11g0112400* in 12 rice inbred lines.**

**(A)** Gene structure of *Os11g0112400*. With Yuetai B (YB) as the reference, genomic variant detected in 11 inbred lines are indicated.

**(B)** Clustering of the 12 inbred lines based on sequence variants of *Os11g0112400*. Red blocks represent the presence of a 42 bp insertion.

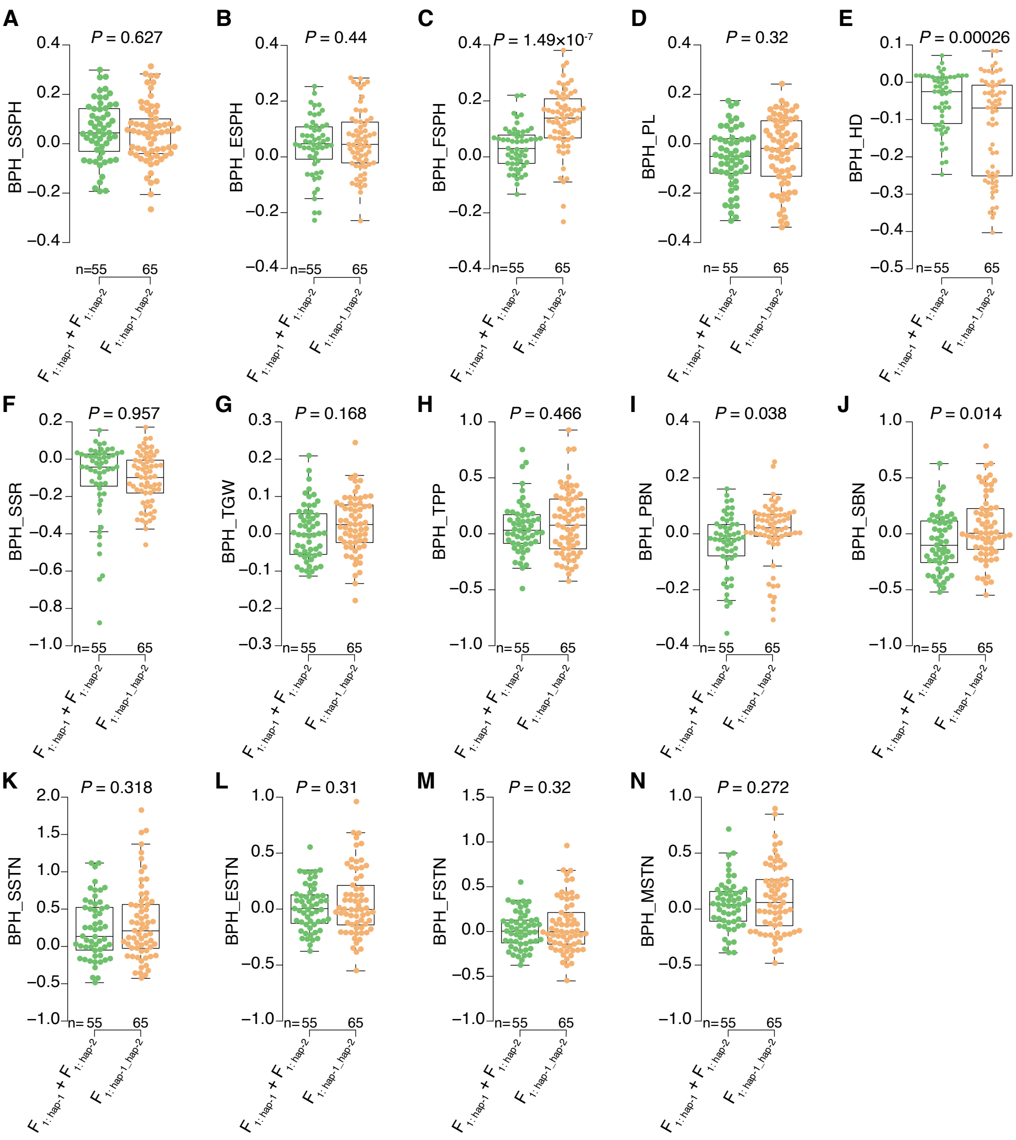


**Supplemental Figure 39. Comparison of heterosis between $F_1$ hybrids with heterozygous haplotype and their homozygous counterparts.** Sequences of six phenylpropanoid biosynthesis genes were used for haplotype analysis of 12 rice inbred lines. Heterosis of 14 traits was compared between $F_1$ hybrids with heterozygous haplotype (haplotype-1_haplotype-2) and their homozygous ones (haplotype-1 and haplotype-2; **A-N**). *P* values are for independent samples *t* test. Abbreviations: better-parent heterosis = BPH, secondary branch number = SBN, tiller number per plant = TPP, seed setting rate = SSR, heading date = HD, thousand grain weight = TGW, seedling stage tiller number = SSTN, elongation stage tiller number = ESTN, flowering stage tiller number = FSTN, maturation stage tiller number = MSTN, seedling stage plant height = SSPH, elongation stage plant height = ESPH, flowering stage plant height = FSPH, panicle length = PL, primary branch number = PBN, and haplotype = hap.

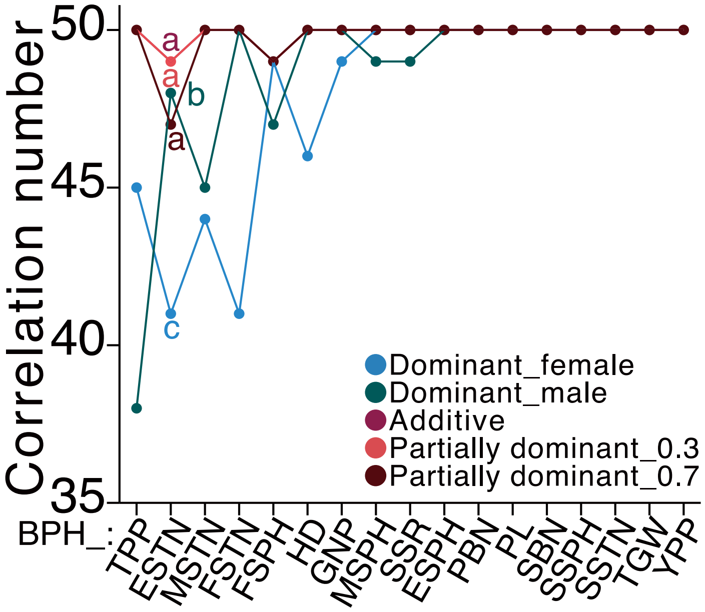


**Supplemental Figure 40. Number of significant correlations between top 50 heterosis-associated metabolites (with different inheritance patterns) and heterosis of 17 agronomic traits.** Analysis of variance (weighted) was performed with the least significant difference used in post hoc test. Abbreviations: better-parent heterosis = BPH, yield per plant = YPP, secondary branch number = SBN, grain number per panicle = GNP, tiller number per plant = TPP, seed setting rate = SSR, heading date = HD, thousand grain weight = TGW, seedling stage tiller number = SSTN, elongation stage tiller number = ESTN, flowering stage tiller number = FSTN, maturation stage tiller number = MSTN, seedling stage plant height = SSPH, elongation stage plant height = ESPH, flowering stage plant height = FSPH, maturation stage plant height = MSPH, panicle length = PL, and primary branch number = PBN.
